# Distinct transcriptomic and hierarchical organization associated with brain hyperconnectivity and hypoconnectivity in autism spectrum disorder

**DOI:** 10.64898/2026.07.31.742105

**Authors:** Abinaya Vairam, Km Bhavna, Lucina Q. Uddin, Bratislav Misic, Dipanjan Roy

## Abstract

Functional hyperconnectivity and hypoconnectivity in autism spectrum disorder (ASD) are typically treated as opposing expressions of a single circuit-level disturbance, but their molecular and hierarchical basis remains unclear. We combined resting-state fMRI from 1,737 individuals from the Autism Brain Imaging Data Exchange (ABIDE I/II) with gene-expression maps from the Allen Human Brain Atlas and show that hyperconnectivity and hypoconnectivity are dissociable neurobiological phenomena, differing in molecular signatures, cortical-hierarchical embedding, age-group profile, and cognitive associations, rather than a single connectivity axis. Hyperconnectivity was concentrated in higher-order cortical and cerebellar regions and was greater in older participants, while hypoconnectivity was consistent across age groups and localized to subcortical and orbitofrontal systems. ASD was associated with reorganization of the sensory-to-transmodal cortical gradient, most pronounced in association networks. Hyperconnectivity- and hypoconnectivity-associated genes showed partially distinct neurotransmitter profiles and differential embedding within cortical hierarchy, both enriched in transmodal cortex and linked to social-cognitive, perceptual, attentional, and reward-related functions. This dissociation was preserved across developmental stage, sex, and symptom severity. These findings indicate hyperconnectivity and hypoconnectivity are not two poles of one process but two separable components of a reproducible molecular-hierarchical architecture, offering a multi-scale framework linking transcriptomic organization to systems-level brain dysfunction in ASD.

## Introduction

Functional hyperconnectivity and hypoconnectivity are consistently reported in autism spectrum disorder (ASD) and are often interpreted as two ends of a single dimension of atypical connectivity, varying in sign but reflecting one underlying process. Recent cross-species studies have demonstrated that distinct patterns of hyperconnectivity and hypoconnectivity can emerge from different autism-associated genetic and molecular mechanisms in animal models, supporting the existence of biologically meaningful connectivity phenotypes ^12^. However, has rarely been tested within human ASD: no study to date has jointly asked whether hyper- and hypoconnectivity differ in their molecular signatures, cortical hierarchial embedding, age profile, and cognitive associations, or instead represent a single organizational axis that merely differs in direction.

Brain structure and function emerge from coordinated gene-expression programs that regulate neuronal development, synaptic signaling, and large-scale network organization ^34,10,74^. Transcriptomic mapping has revealed spatial gradients of gene expression that align with cortical organization, functional specialization, and developmental trajectories ^34,10,32^, suggesting close coupling between molecular architecture and brain function. However, how alterations in these molecular systems relate to macroscale brain dysfunction in neurodevelopmental conditions such as ASD remains incompletely understood.

ASD is a heterogeneous neurodevelopmental condition characterized by differences in social communication alongside restricted and repetitive behaviors ^4^. Neuroimaging studies consistently report atypical functional connectivity in ASD, including both hyper-and hypoconnectivity across cortical and subcortical systems ^76,71,17,37^, that vary across development and suggest disruptions in large-scale network organization rather than uniform connectivity deficits ^56,62^, including altered core–periphery organization and segregation–integration balance ^31,62^. Yet the biological correlates of these alterations remain poorly understood.

Large-scale brain organization can be captured by continuous functional gradients spanning unimodal sensory to transmodal association cortices ^45,38^, providing a low-dimensional link between molecular architecture and distributed functional systems ^51,10,32^. Transmodal regions integrate sensory, affective, and cognitive information and support social cognition, executive function, and adaptive behavior ^51,69^. In ASD, atypical gradient organization, including reduced sensory–transmodal differentiation, has been reported ^36,77^, suggesting that functional connectivity alterations reflect broader disruptions in cortical hierarchy.

Imaging–transcriptomic approaches have begun to bridge molecular and systems-level brain data, showing that functional organization aligns with transcriptional gradients ^60,3^. In ASD, transcriptomic signatures related to synaptic signaling, neuronal development, and neurotransmission have been linked to altered brain structure and function ^29,6,44^, though how transcriptomic organization relates to cortical hierarchy and cognitive systems remains largely unexplored.

Hong et al. ^36^ showed that ASD involves alterations in the principal functional gradient, reflecting atypical cortical hierarchy, while Berto et al. ^6^ linked regional connectivity alterations to cortical gene-expression patterns. However, these studies addressed cortical hierarchy and transcriptomic organization largely independently, leaving unclear whether hyper- and hypoconnectivity exhibit distinct neurochemical signalling associated transcriptomic signatures, differential hierarchical embedding, or divergent cognitive associations, and whether such molecular–functional relationships generalize across development, sex, and clinical heterogeneity.

Among implicated molecular systems, neurotransmitter pathways play a central role. Beyond the historically dominant glutamatergic and GABAergic models ^63,55^, emerging evidence implicates broader neuromodulatory systems, including dopaminergic, cholinergic, noradrenergic, oxytocinergic, vasopressinergic, and endocannabinoid pathways ^27,2,35^, in synaptic plasticity, reward processing, attention, social cognition, learning, and adaptive behavior. This suggests ASD-related brain dysfunction may reflect coordinated variation across multiple neurotransmitter networks rather than a single excitation–inhibition imbalance, and that understanding how these systems are organized across large-scale brain networks may provide important insight into ASD neurobiology. ASD-related connectivity alterations also show substantial developmental variability, with evidence suggesting age-dependent changes in cortical organization, network integration, and functional specialization ^56,62^. Given the marked heterogeneity of ASD, it remains unclear whether molecular signatures associated with large-scale brain dysfunction are conserved across developmental stage, sex, and symptom severity, or whether they instead reflect subgroup-specific biological processes, a question important for parsing ASD heterogeneity and identifying organizational principles that generalize across diverse ASD phenotypes.

In sum, prior work has examined functional connectivity, cortical hierarchy, and transcriptomic organization largely in isolation, each addressing one piece of a puzzle whose pieces have not been shown to fit together. Hong et al. established that ASD involves atypical cortical-gradient organization; Berto et al. established that regional connectivity alterations track cortical gene-expression patterns^36,6^. Neither, nor the broader literature, has tested whether hyper- and hypoconnectivity are two expressions of the same underlying organization or two dissociable phenomena with distinct molecular, hierarchical, and cognitive correlates, and whether any such dissociation generalizes across development, sex, and clinical subgroups.

Here, we tested the hypothesis that ASD-associated hyper- and hypoconnectivity reflect partially distinct molecular, hierarchical, and cognitive organizational patterns rather than a single uniform connectivity alteration. We integrated resting-state functional connectivity, cortical gradient analysis, imaging–transcriptomics, neurochemical signalling system organization, and meta-analytic cognitive decoding within a unified framework, using resting-state fMRI from the Autism Brain Imaging Data Exchange (ABIDE I and II) and gene-expression maps from the Allen Human Brain Atlas (AHBA). We examined how transcriptomic signatures relate to hyper- and hypoconnectivity patterns, how these molecular systems are embedded within cortical hierarchy, how they map onto cognitive functional architecture, and whether these organizational principles are preserved across developmental stage, sex, and clinical severity. We predicted that hyper-and hypoconnectivity would exhibit partially distinct transcriptomic signatures, differential alignment with the sensory–transmodal hierarchy, and distinct cognitive associations, consistent with separable neurobiological processes rather than opposite ends of a single connectivity continuum. Unlike prior work examining these components separately, our framework evaluates how neurochemical signalling associated molecular architecture, functional connectivity, cortical hierarchy, and cognitive systems are jointly organized across biological scales.

## Results

### Developmental patterns of hyperconnectivity and hypoconnectivity in ASD

Following the imaging–transcriptomic workflow illustrated in (Figure 1), we characterized the spatial organization of functional connectivity alterations, by comparing ASD and control participants across the whole brain **(Figure 2)**. ASD exhibited coexisting patterns of hyperconnectivity and hypoconnectivity distributed across cortical and subcortical regions (Figure 2a), consistent with previous reports ^17,37,71,76^ . Hyperconnectivity predominantly involved frontal, cingulate, parietal, and cerebellar cortices, whereas hypoconnectivity localized primarily to subcortical structures, including the nucleus accumbens, thalamus, putamen, and orbitofrontal-temporal regions.

**Figure 1.**
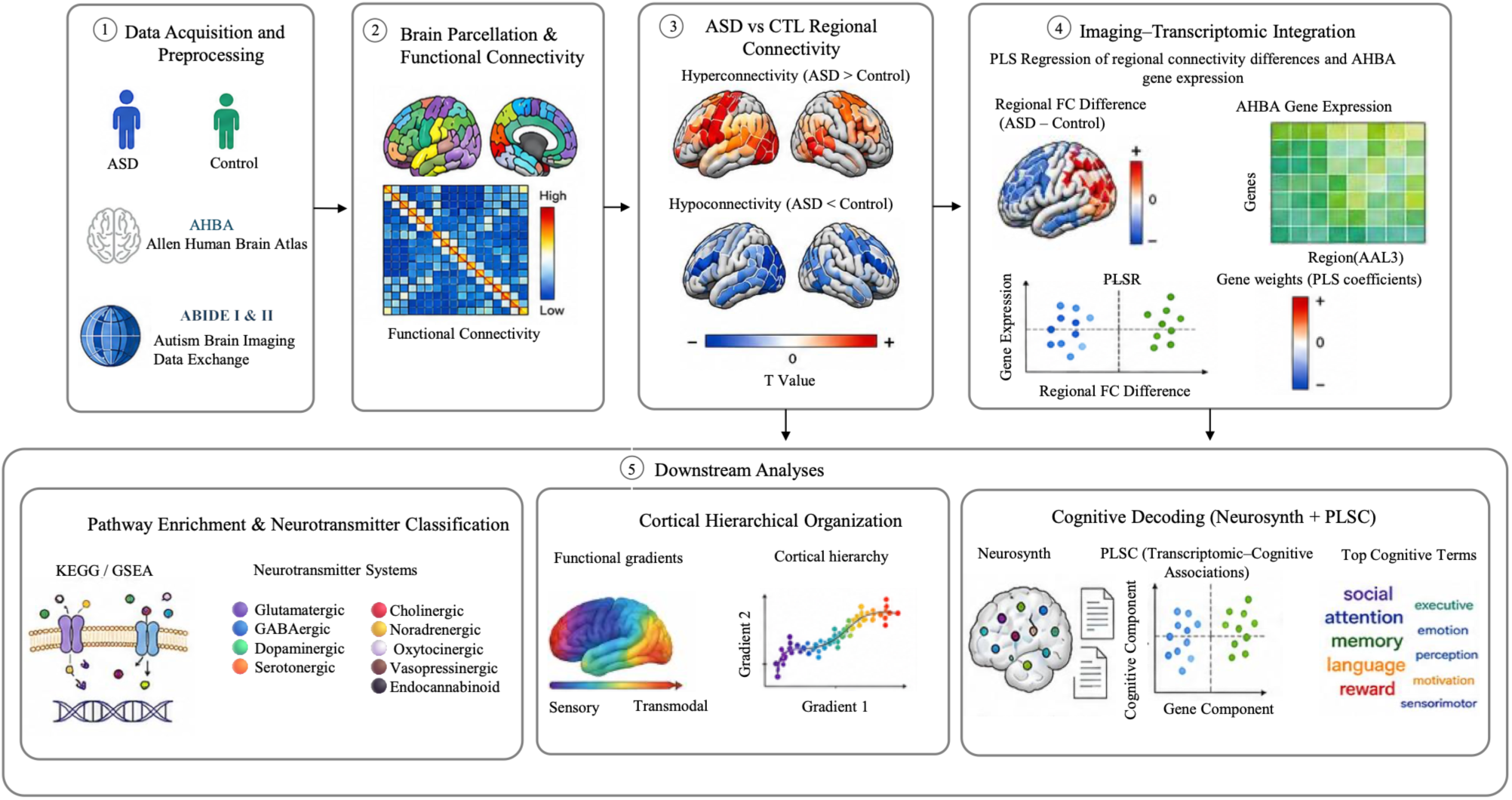
Overview of the imaging–transcriptomic analysis pipeline. Resting-state functional MRI data from the ABIDE I and II cohorts were preprocessed and parcellated using the AAL3 atlas to generate regional functional connectivity matrices. Regional functional connectivity differences between individuals with autism spectrum disorder (ASD) and healthy controls (CTL) were integrated with Allen Human Brain Atlas (AHBA) gene expression profiles using partial least squares regression (PLSR) to identify transcriptomic signatures associated with altered connectivity. The resulting gene signatures were characterized through pathway enrichment and neurochemical signalling system classification, mapped onto the cortical functional hierarchy and linked to cognitive functions using Neurosynth-based cognitive decoding and partial least squares correlation (PLSC) analysis.

**Figure 2:**
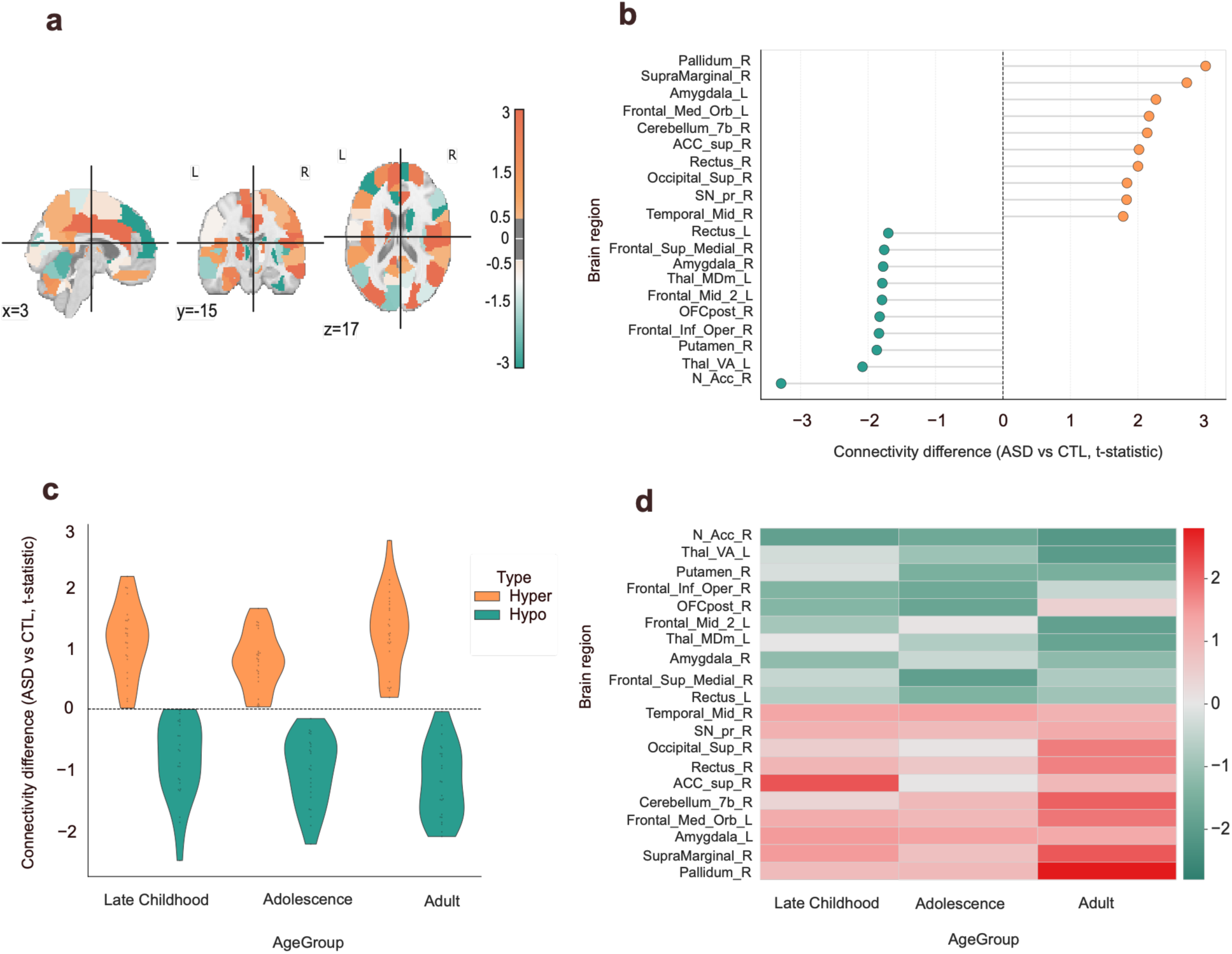
Functional connectivity alterations in ASD. (a) Whole-brain maps of functional connectivity differences (ASD vs CTL). (b) Region-wise connectivity differences across brain regions. (c) Distributions of connectivity differences across age groups. (d) Heatmap of regional connectivity differences across developmental stages.

Regional ranking of the ten strongest positive and negative connectivity differences **(Figure 2b)** showed the largest positive effects in the supramarginal cortex, pallidum, and cerebellum, whereas the strongest negative effects occurred in the nucleus accumbens and multiple thalamic nuclei. The ranked regions demonstrated a consistent spatial organization, with hyperconnectivity preferentially involving higher-order cortical and cerebellar regions and hypoconnectivity concentrated within subcortical structures.

Across the three cross-sectional age groups (**Figures 2c,d**), the overall spatial organization of connectivity alterations was preserved. Hyperconnectivity showed descriptively larger positive connectivity differences in adulthood (mean t = 1.274; SD = 0.661) than in late childhood (t = 1.128; SD = 0.600), whereas hypoconnectivity remained consistently negative across all age groups (late childhood: −0.851; SD = 0.657; adolescence: −1.016; SD = 0.570; adulthood: −1.213; SD = 0.605). However, these age-group differences did not reach statistical significance (P = 0.536), indicating that the observed trends were descriptive rather than reflecting significant age-related effects.

Sex-stratified analyses (**Figures S12–S13**) revealed broadly similar connectivity architectures in males and females, although males showed more spatially distributed alterations, whereas females exhibited relatively focal patterns. Across all subgroup analyses (**Figures S9–S13**), hyperconnectivity consistently involved higher-order cortical association regions, while hypoconnectivity remained localized to the nucleus accumbens, thalamus, putamen, and amygdala—together indicating that the spatial organization of connectivity alterations was preserved across demographic subgroups.

### Gene expression organization and pathway enrichment across neurochemical systems

To characterize the molecular architecture underlying ASD-associated functional connectivity alterations, we examined the 172 leading-edge genes identified via imaging–transcriptomic analysis (69 associated with hyperconnectivity and 103 with hypoconnectivity; **Figure 3**). Hierarchical clustering of these genes revealed coordinated enrichment across several neurotransmitter and synaptic signalling pathways—including glutamatergic, GABAergic, dopaminergic, serotonergic, and cholinergic synapses—alongside calcium signalling, axon guidance, synaptic vesicle cycling, long-term potentiation, and neuroactive ligand–receptor interaction pathways (**Figure 3a**). Based on the significantly enriched KEGG pathways (P_FDR_ < 0.05), eight neurochemical signalling systems were retained for further analyses (**Figure 3c,d**). Notably, the substantial overlap in gene membership across these pathways indicates that diverse neurotransmitter and neuropeptide systems converge on shared intracellular signalling mechanisms, rather than operating through distinct, pathway-specific molecular programmes. The hierarchical clustering solution showed high internal consistency, with cophenetic correlation coefficients of 0.852 for genes and 0.943 for pathways, supporting the robustness of this transcriptomic architecture.

**Figure 3:**
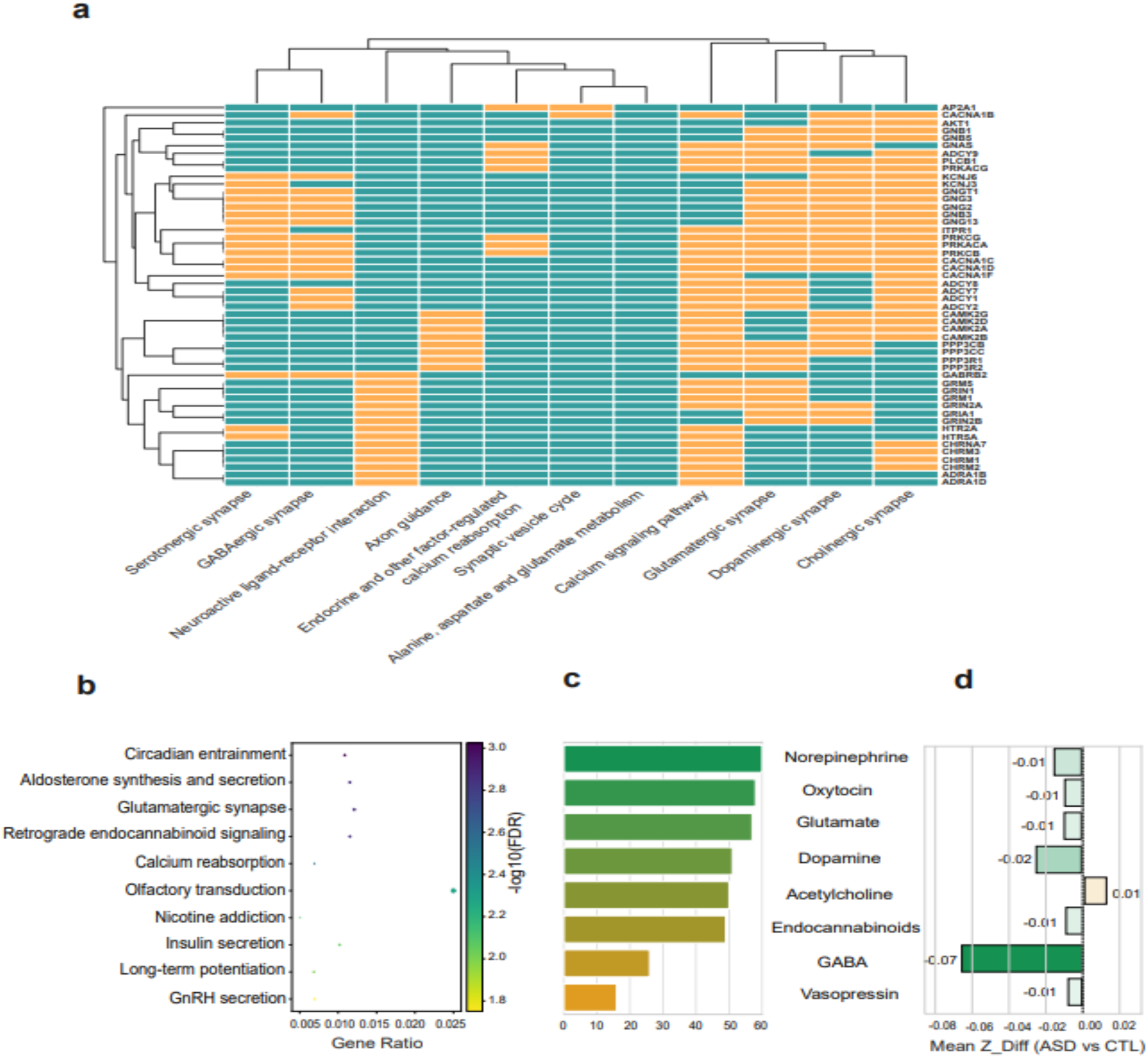
Pathway enrichment and neurochemical signalling system associations of ASD-associated genes. (a) Colormap of gene–pathway relationships across enriched pathways for selected ASD-associated genes. (b) Functional enrichment plot showing pathways by gene ratio and statistical significance. (c) Number of ASD relevant genes associated with each neurotransmitter system. (d) Mean standardized functional connectivity differences (ASD vs CTL) across neurotransmitter systems.

Several core signalling genes recurred across pathways, including calcium channel subunits (CACNA1 family), G-protein components (GNB and GNG families), adenylyl cyclases (ADCY family), phospholipase signalling molecules (PLCB family), and protein kinases (PRK and CAMK2 families), reflecting extensive representation of second-messenger signalling and calcium-dependent processes. Glutamate receptor genes (GRIN, GRIA, and GRM families) and GABA receptor genes (GABRB family) were also prominently enriched, implicating both excitatory and inhibitory synaptic systems. The recurrence of these genes across multiple pathways further underscores the substantial molecular overlap among diverse neurotransmitter systems.

Pathway enrichment analysis identified significant enrichment of circadian entrainment (NES = 2.043, P_FDR_ < 0.001), glutamatergic synapse (NES = 2.064, P_FDR_ = 0.0014), retrograde endocannabinoid signalling (NES = 2.005, P_FDR_ = 0.0014), long-term potentiation (NES = 1.865, P_FDR_ = 0.010), and aldosterone synthesis and secretion pathways (**Figure 3b**). Axon guidance and synaptic vesicle cycling were also enriched, highlighting molecular processes involved in circuit formation, maintenance, and remodelling. Collectively, these pathways span neurotransmission, synaptic plasticity, neuroendocrine regulation, and homeostatic signalling, indicating that ASD-associated transcriptomic signatures are distributed across multiple biological processes rather than confined to a single neurotransmitter system. The concurrent enrichment of glutamatergic, endocannabinoid, and activity-dependent signalling pathways further supports coordinated neuromodulatory and synaptic mechanisms as a basis for large-scale functional connectivity alterations in ASD.

Quantification of neurochemical signalling system representation revealed a broad distribution of ASD-associated genes across multiple pathways (**Figure 3c**). Noradrenergic-associated genes constituted the largest group (n = 60, 16.4%), followed by oxytocinergic (n = 58, 15.8%), glutamatergic (n = 57, 15.5%), dopaminergic (n = 51, 13.9%), cholinergic (n = 50, 13.6%), endocannabinoid (n = 49, 13.3%), GABAergic (n = 26, 7.1%), and vasopressinergic genes (n = 16, 4.4%). This representation differed significantly across neurochemical systems (P < 0.01), indicating a non-uniform distribution of ASD-associated genes among neurochemical and signalling pathways.

Directional analysis demonstrated only modest differences in connectivity-associated effects across neurochemical signalling systems (**Figure 3d**). GABAergic pathways exhibited the largest negative mean Z-difference (−0.07), whereas dopaminergic pathways showed a smaller negative deviation (−0.02). Endocannabinoid, glutamatergic, noradrenergic, oxytocinergic, and vasopressinergic pathways each exhibited slight negative shifts (approximately −0.01), while cholinergic pathways showed a modest positive deviation (+0.01). Overall, the magnitude of directional differences was small across neurotransmitter systems, suggesting broadly distributed transcriptomic contributions to ASD-associated functional connectivity alterations.

Developmental analyses demonstrated highly consistent pathway enrichment profiles across late childhood, adolescence, and adulthood (**Figures S1–S3**). Calcium signalling, glutamatergic synapse, dopaminergic synapse, cholinergic synapse, neuroactive ligand–receptor interaction, and endocannabinoid signalling remained recurrently enriched across all age groups, whereas pathways related to circadian entrainment, endocrine regulation, and long-term potentiation exhibited greater age-dependent variation in enrichment magnitude. Similar pathway architectures were observed across male and female participants (**Figures S4–S5**), although female participants showed relatively greater enrichment of immune- and cell-adhesion-related pathways, whereas males exhibited stronger enrichment of neurotransmission- and plasticity-related pathways. Severity-stratified analyses further demonstrated persistent enrichment of neurotransmitter release, synapse assembly, presynaptic organization, postsynaptic organization, axonal transport, neuronal projection, receptor trafficking, and plasticity-related pathways across lower-, intermediate-, and higher-symptom-burden groups **(Figures S6–S8)**.

Across all subgroup analyses, dopaminergic, glutamatergic, cholinergic, and endocannabinoid pathways consistently exhibited the strongest enrichment, while circadian entrainment, neuroactive ligand–receptor interaction, and endocrine signalling remained prominent neuromodulatory components of the ASD-associated molecular profile. The reproducibility of these enrichment patterns across developmental stage, sex, and symptom-burden groups indicates a stable molecular architecture in which multiple neurotransmitter systems converge on shared signalling and plasticity-related pathways underlying large-scale functional connectivity alterations in ASD.

### Neurochemical signalling system specific associations of hyper- and hypoconnectivity

To investigate the molecular architecture underlying ASD-related functional connectivity alterations, we compared regional gene-expression profiles associated with hyperconnectivity and hypoconnectivity across cortical and subcortical systems (**Figure 4**). Heatmap visualization revealed a clear spatial organization of transcriptomic signatures, with relatively higher normalized expression across association cortical networks—including the default mode, frontoparietal, and limbic networks—whereas subcortical regions, including the thalamus, caudate, putamen, and amygdala, generally showed lower expression (**Figure 4a**). Overall cortical expression exhibited a slightly higher median than subcortical expression (0.771 vs. 0.763), although this difference was not statistically significant (P_FDR_ = 0.067).

**Figure 4:**
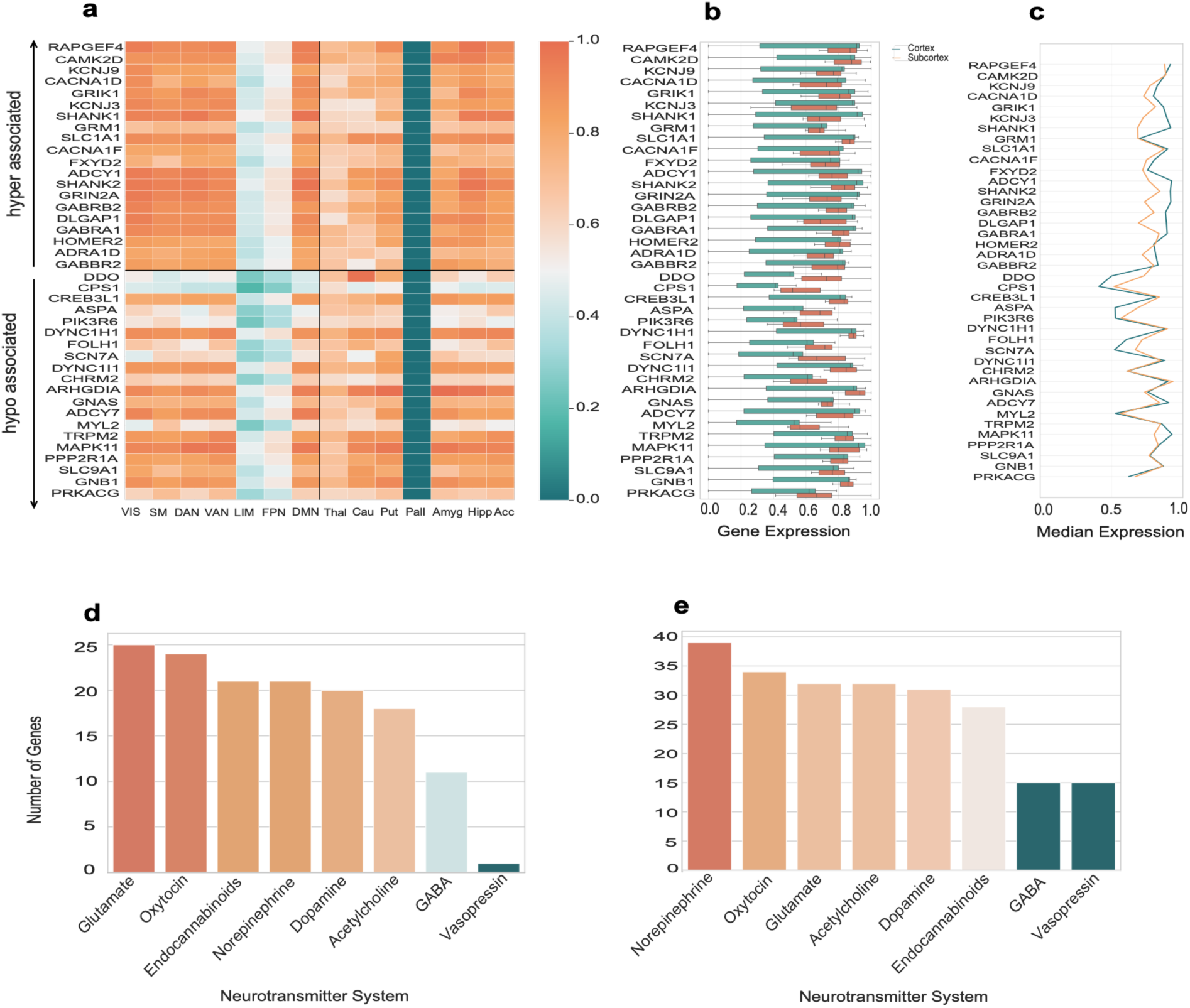
Gene expression profiles of top hyper- and hypoconnectivity-associated genes. (a) Heatmap of gene expression across brain regions for the top 10 hyper- and top 10 hypo-associated genes. (b) Gene expression distribution across cortical and subcortical regions. (c) Median gene expression profiles across regions. (d) Distribution of top hyper-associated genes across neurotransmitter systems. (e) Distribution of top hypo-associated genes across neurotransmitter systems.

Direct comparison of cortical and subcortical gene-expression profiles revealed broadly similar patterns between the two tissue compartments (**Figure 4b**). Median regional expression profiles were moderately correlated between cortex and subcortex (**Figure 4c**), with an average cortex–subcortex expression difference of 0.0229 (95% bootstrap CI: −0.0059 to 0.0513; P_FDR_ = 0.138). Together, these findings point to broadly similar transcriptomic organization across cortical and subcortical regions, with a tendency toward higher expression in cortical association areas.

Hyperconnectivity- and hypoconnectivity-associated genes showed broadly similar neurotransmitter pathway distributions (**Figure 4d,e**). Hyperconnectivity-associated genes were most frequently assigned to glutamatergic (n = 25), oxytocinergic (n = 24), endocannabinoid (n = 21), noradrenergic (n = 21), dopaminergic (n = 20), and cholinergic (n = 18) pathways, with lower representation in GABAergic (n = 11) and vasopressinergic (n = 1) pathways. Hypoconnectivity-associated genes showed the greatest representation in noradrenergic (n = 39), oxytocinergic (n = 34), glutamatergic (n = 32), cholinergic (n = 32), dopaminergic (n = 31), and endocannabinoid (n = 28) pathways, with fewer genes assigned to GABAergic (n = 15) and vasopressinergic (n = 15) pathways. Despite these numerical differences, overall neurotransmitter pathway composition did not differ significantly between hyperconnectivity- and hypoconnectivity-associated gene sets (P = 0.257), indicating substantial overlap in neurotransmitter-system involvement.

This transcriptomic architecture was highly reproducible across subgroup analyses (**Figures S14–S21**). Cortical enrichment of both hyperconnectivity- and hypoconnectivity-associated genes was preserved across childhood, adolescence, and adulthood, although the magnitude of enrichment differed significantly among developmental stages (P = 0.011). Expression profiles were similar between male and female participants, with no significant sex-related differences (P_FDR_ = 0.080). Across lower-, intermediate-, and higher-symptom-burden groups, the overall molecular architecture remained conserved, although enrichment magnitude differed significantly with symptom burden (P_FDR_ = 0.020). Across all subgroup analyses, glutamatergic, endocannabinoid, oxytocinergic, dopaminergic, and noradrenergic pathways consistently represented the predominant neurotransmitter systems.

Notably, genes involved in neurotransmitter signalling, synaptic scaffolding, receptor trafficking, and calcium-dependent intracellular signalling repeatedly emerged among the strongest contributors across all subgroup analyses (**Figures S14–S21**), pointing to consistent involvement of molecular processes underlying synaptic communication, plasticity, and neuromodulatory signalling.

To assess whether the AHBA-derived transcriptomic signatures were broadly represented across the developmental age range of the ABIDE cohort, we performed a supplementary sensitivity analysis using the BrainSpan developmental transcriptomic atlas **(Figure S32 and Table S4)**. All 40 hyperconnectivity- and hypoconnectivity-associated genes identified from the AHBA analysis were represented in the BrainSpan dataset (100% data completeness). Overall, 95% of genes exhibited less than two-fold developmental variation (median coefficient of variation = 0.177; median adult-to-childhood fold change = 1.221), and 80% were consistently expressed across childhood, adolescence, and adulthood. Developmental trajectory analysis revealed heterogeneous expression patterns within both gene sets, with no significant difference in trajectory distributions between hyperconnectivity- and hypoconnectivity-associated genes (Fisher’s exact test, *P* = 0.235). These findings support the developmental applicability of the identified transcriptomic signatures across the age range represented in the ABIDE cohort.

As additional robustness analyses, BrainSMASH spatial-null testing demonstrated that latent variables LV2 (*P_10K SPIN_* = 0.006), LV4 (*P_10K SPIN_* = 0.018), and LV5 (*P_10K SPIN_* = 0.003) remained significant after accounting for spatial autocorrelation, whereas LV1 (*P_10K SPIN_* = 0.696) and LV3 (*P_10K SPIN_* = 0.063) did not exceed the significance threshold (Figure S31). Furthermore, leave-one-site-out (LOSO) and leave-one-region-out (LORO) analyses showed that the direction of hyperconnectivity- and hypoconnectivity-associated gene signatures was highly preserved across repeated analyses. LOSO analyses demonstrated a mean direction stability of 98.4% across all study genes, whereas LORO analyses showed a similarly high mean direction stability of 96.7%. Hypoconnectivity-associated genes exhibited complete directional stability across region-dropped analyses (20/20 genes, 100%), while hyperconnectivity-associated genes also remained highly consistent (mean direction stability = 93.3%). Together, these analyses demonstrate that the identified imaging–transcriptomic associations were robust to spatial autocorrelation, imaging-site heterogeneity, and regional exclusion, indicating that the observed molecular signatures were not driven by individual sites or brain regions.

Collectively, these findings demonstrate that ASD-related hyperconnectivity and hypoconnectivity share overlapping neurochemical-system signatures with preferential cortical enrichment. Their reproducibility across developmental stage, sex, symptom burden, and multiple sensitivity analyses supports a robust relationship between transcriptomic organization and large-scale functional connectivity alterations in ASD.

### Reorganization of cortical functional gradients and transcriptomic organization

To investigate how these alterations are organized at the macroscale level of brain hierarchy, we examined cortical functional gradients (**Figure 5**). In controls, cortical regions formed a compact and coherent distribution in gradient space (**Figure 5a**), whereas the ASD group showed a more dispersed configuration, particularly along Gradient 1 (variance: 0.00175 vs. 0.00058; variance ratio = 3.04; P_FDR_ = 0.0002). Density distributions further revealed a shift toward higher Gradient 1 values in ASD, indicating expansion along the sensory-to-transmodal axis.

**Figure 5:**
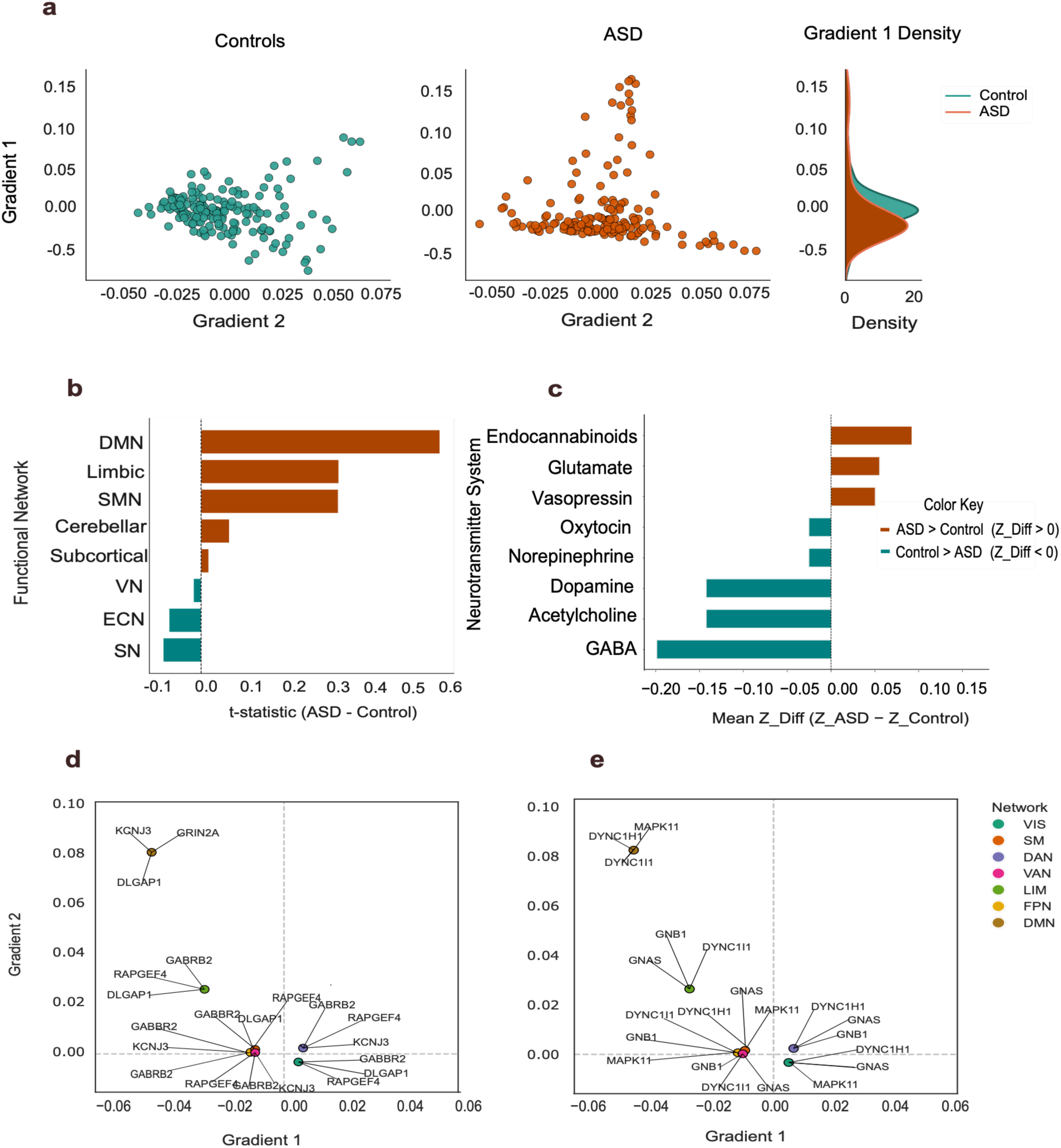
Functional gradient alterations and network-level differences in ASD. (a) Scatter plots of brain regions in Gradient 1–Gradient 2 space for controls and ASD, with Gradient 1 density distributions. (b) Bar plot of t-statistics (ASD − Control) showing gradient differences across functional brain networks. (c) Bar plot of mean Z-score differences (ASD − Control) across neurotransmitter systems. (d) Gradient space distribution of hyper-correlated genes associated with functional organization. (e) Gradient space distribution of hypo-correlated genes associated with functional organization.

At the network level (**Figure 5b**), gradient reorganization was concentrated within higher-order association systems. The default mode network showed the largest positive shift (mean t = 0.616; SD = 0.368; P_FDR_ = 0.0040), followed by the limbic (mean t = 0.355; SD = 1.257) and sensorimotor (mean t = 0.354; SD = 0.708) networks. In contrast, salience (mean t = −0.098; SD = 1.369) and executive control (mean t = −0.083; SD = 1.210) networks showed significant reductions relative to controls, whereas visual (mean t = −0.020; SD = 1.109), cerebellar (mean t = 0.073; SD = 0.847), and subcortical (mean t = 0.020; SD = 1.219) systems displayed comparatively smaller alterations.

Neurochemical specific differences (**Figure 5c**) revealed a pattern consistent with earlier analyses. Glutamatergic (mean Z = 0.055) and endocannabinoid (mean Z = 0.092) systems showed positive shifts, whereas GABAergic (mean Z = −0.199), cholinergic (mean Z = −0.142), and dopaminergic (mean Z = −0.142) systems showed negative effects. Oxytocinergic (mean Z = −0.025) and noradrenergic (mean Z = −0.025) systems showed smaller negative deviations, whereas the vasopressinergic system (mean Z = 0.050) showed a modest positive effect.

Projection of transcriptomic signatures onto cortical gradient space revealed distinct spatial organization of hyperconnectivity- and hypoconnectivity-associated genes (**Figure 5d,e**). Hyperconnectivity-associated genes were centred toward transmodal cortical territories (median Gradient 1 = −0.0197; centroid = −0.0221), whereas hypoconnectivity-associated genes were centred toward sensory and subcortical territories (median Gradient 1 = 0.0314; centroid = 0.0271). These findings indicate that hyperconnectivity- and hypoconnectivity-associated genes occupy distinct positions along the cortical functional hierarchy.

Age-stratified analyses demonstrated that the overall architecture of cortical gradient reorganization was preserved across late childhood, adolescence, and adulthood (**Figures S22–S24**). Within each developmental stage, the ASD group exhibited significantly greater Gradient 1 variance than controls (late childhood: P_FDR_ = 0.024; adolescence: P_FDR_ = 0.029; adulthood: P_FDR_ = 0.016). Adolescence showed the greatest Gradient 1 variance within the ASD group (variance = 0.004), whereas late childhood exhibited the largest ASD-to-control variance ratio (1.37), indicating developmental differences in cortical hierarchical organization without a monotonic increase in gradient alterations with age. Across all developmental stages, higher-order association networks—including the default mode, limbic, and frontoparietal systems—showed the largest deviations from controls, whereas primary sensory and visual networks showed comparatively smaller shifts. Neurotransmitter-associated gradient effects were similarly consistent across development, with glutamatergic and endocannabinoid systems showing positive associations with gradient expansion, and GABAergic, dopaminergic, and cholinergic systems showing negative associations (**Figures S22–S24**).

Comparable patterns were observed across sex-stratified analyses (**Figures S25–S26**). Both male and female ASD participants exhibited greater Gradient 1 variance than controls, with ASD-to-control variance ratios of 2.23 and 1.96, respectively. Distributional differences remained significant in both males (P_FDR_ = 0.0043) and females (P_FDR_ < 0.001), indicating a conserved spatial organization of cortical gradient alterations despite modest differences in network-specific effect sizes. Hyperconnectivity-associated genes preferentially localized to intermediate and transmodal cortical regions, whereas hypoconnectivity-associated genes extended toward sensory and subcortical territories in both sexes (**Figures S25–S26**).

Symptom-stratified analyses similarly demonstrated preserved cortical gradient organization across lower-, intermediate-, and higher-severity ASD subgroups (**Figures S27–S29**). Gradient 1 variance ratios were 3.04, 2.16, and 7.26 for the lower-, intermediate-, and higher-severity groups, respectively (P_FDR_ < 0.001). Glutamatergic, calcium-signalling, endocannabinoid, and synaptic-plasticity genes remained preferentially localized to transmodal cortical territories, whereas hypoconnectivity-associated genes showed broader distributions across the cortical hierarchy. Collectively, these findings demonstrate systematic reorganization of cortical functional hierarchy in ASD, with distinct hyperconnectivity- and hypoconnectivity-associated transcriptomic signatures preserved across developmental stage, sex, and symptom severity.

### Neurochemical signalling system associated cognitive mapping using Neurosynth

To contextualize the identified transcriptomic signatures in terms of cognitive function, we performed neurotransmitter-specific meta-analytic decoding using Neurosynth (**Figure 6a**). Across all neurotransmitter systems, the dorsolateral prefrontal cortex, cuneus, fusiform gyrus, orbitofrontal cortex, and striatum were consistently represented, indicating a common anatomical substrate underlying neurotransmitter-associated gene-expression profiles. Distinct functional specializations were also evident: glutamatergic and endocannabinoid systems showed stronger associations with visual and associative processing; GABAergic and cholinergic systems were preferentially linked to executive and prefrontal functions; dopaminergic and noradrenergic systems were enriched for reward-related processes; and oxytocinergic and vasopressinergic systems were associated with social cognition and face-processing networks. Collectively, these findings demonstrate that neurotransmitter-associated transcriptomic signatures preferentially map onto cognitive domains relevant to ASD, including social cognition, executive function, visual processing, and motivation and reward.

**Figure 6.**
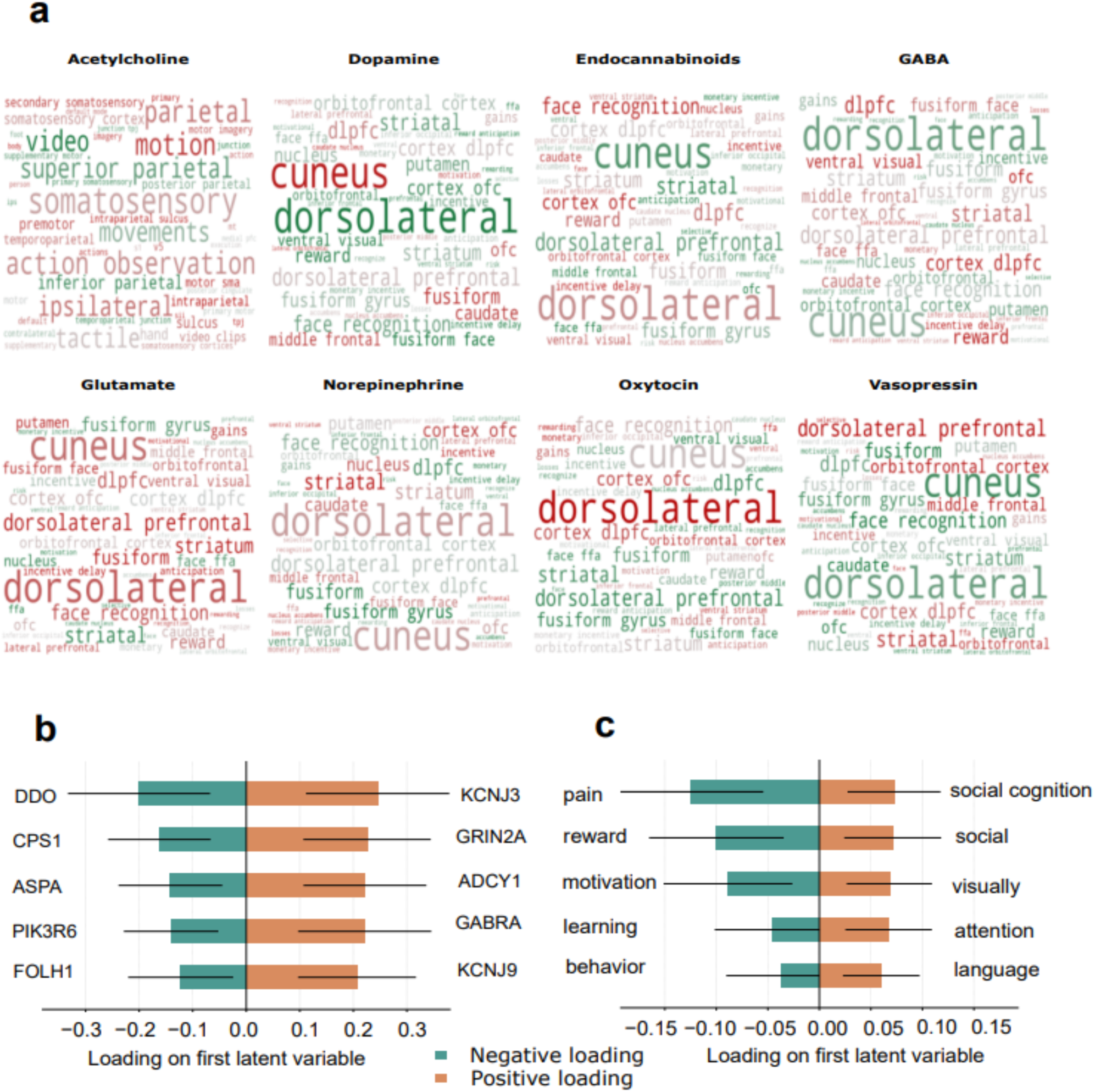
Transcriptomic - cognitive associations revealed by partial least squares correlation (PLSC). (a) Neurosynth cognitive decoding of neurotransmitter-associated transcriptomic signatures. Word size is proportional to association strength (b) Top five positive and negative gene loadings for the first latent variable (LV1). (c) Top five positive and negative cognitive-term loadings for LV1.

To investigate the relationship between transcriptomic organization and cognitive functional architecture, we performed partial least squares correlation (PLSC) between regional gene-expression profiles and Neurosynth-derived cognitive activation maps (**Figure 6b,c**). The first latent variable (LV1) accounted for the largest share of covariance between the transcriptomic and cognitive datasets (67.0%) and showed the strongest association (r = 0.657). LV1 was highly significant following permutation testing (10,000 permutations; P < 0.001) and remained significant after BrainSMASH spatial-null correction (*P_10K SPIN_* = 0.001), indicating that the observed transcriptomic–cognitive correspondence exceeded that expected under spatial autocorrelation alone. Although additional latent variables were also significant (LV2: 18.4% covariance explained, r = 0.447, *P_10K SPIN_* = 0.003; LV3: 7.0%, r = 0.416, *P_10K SPIN_* = 0.016; LV5: 1.4%, r = 0.492, *P_10K SPIN_* = 0.001; all BrainSMASH *P_10K SPIN_* = 0.001), LV1 explained the majority of shared covariance and was therefore selected for biological interpretation.

Gene loadings for LV1 defined a molecular axis along which positively weighted genes were predominantly associated with synaptic signalling, whereas negatively weighted genes were enriched for metabolic and broader cellular processes (**Figure 6b**). The strongest positive loadings were observed for KCNJ3 (loading = 0.245, BSR = 3.59), GRIN2A (0.226, BSR = 3.72), ADCY1 (0.222, BSR = 3.79), GABRA1 (0.221, BSR = 3.49), and KCNJ9 (0.207, BSR = 3.68), whereas the strongest negative loadings were observed for DDO (−0.200, BSR = −2.96), CPS1 (−0.162, BSR = −3.32), ASPA (−0.142, BSR = −2.86), PIK3R6 (−0.140, BSR = −3.12), and FOLH1 (−0.122, BSR = −2.45). Positively weighted genes included several involved in neurotransmission, calcium-dependent signalling, ion-channel regulation, and synaptic plasticity: GRIN2A and ADCY1 participate in glutamatergic and calcium signalling pathways, GABRA1 mediates inhibitory neurotransmission, and members of the KCNJ family regulate neuronal membrane excitability. Negatively weighted genes, by contrast, were primarily associated with amino acid metabolism, intracellular signalling, and broader cellular maintenance. Together, these findings suggest that LV1 captures a transcriptomic continuum spanning synaptic signalling and metabolic biological processes.

The corresponding cognitive loadings defined a complementary functional axis (**Figure 6c**). The strongest positive loadings were observed for social cognition (loading = 0.072, BSR = 3.15), social processing (0.071, BSR = 2.97), visual processing (0.068, BSR = 3.21), attention (0.067, BSR = 3.14), and language (0.060, BSR = 3.17). Among negatively weighted terms, pain (−0.124, BSR = −3.49), reward (−0.100, BSR = −3.00), and motivation (−0.089, BSR = −2.78) showed the strongest negative contributions, whereas learning (−0.045, BSR = −1.75) and behaviour (−0.037, BSR = −1.60) showed comparatively weaker negative loadings. These findings suggest that brain regions enriched for synaptic-signalling genes are preferentially associated with higher-order social, perceptual, attentional, and language-related functional systems, whereas regions enriched for metabolic and broader cellular processes correspond more strongly to reward-, motivation-, and pain-related functional domains.

Notably, several highly weighted genes identified by PLSC—including GRIN2A, ADCY1, GABRA1, KCNJ3, and KCNJ9—were also consistently identified in the imaging–transcriptomic, pathway-enrichment, and cortical-gradient analyses, supporting convergent involvement of glutamatergic, GABAergic, calcium-dependent, and neuromodulatory signalling pathways across analytical levels. Together, these findings provide convergent evidence linking neurotransmitter-associated transcriptomic signatures with large-scale functional connectivity, cortical hierarchy, and cognitive functional architecture.

## Discussion

The present study integrates functional connectivity, cortical hierarchy, imaging–transcriptomics, neurochemical signalling system organization, and cognitive functional mapping to characterize large-scale brain organization in (ASD). Although previous studies have independently reported atypical functional connectivity, altered cortical gradients, and transcriptomic associations in ASD ^36,44,6^, how these levels of organization relate to one another has remained largely unresolved. By integrating connectivity alterations, developmental trajectories, transcriptomic organization, cortical hierarchy, and cognitive functional architecture within a unified framework, the present findings identify a reproducible molecular–hierarchical organization of ASD characterized by consistent spatial associations among neurotransmitter-related gene-expression profiles, large-scale brain organization, and cognitive functional systems across developmental and clinical subgroups.

This dissociation directly distinguishes the present findings from Berto et al.⁶, the most closely related prior imaging–transcriptomic study of ASD connectivity. Berto et al.⁶ linked regional connectivity alterations to cortical gene expression treating ASD-related connectivity change as a single, unsigned quantity, without asking whether increases and decreases in connectivity arise from dissociable molecular and hierarchical substrates. The present study instead analyzes hyperconnectivity and hypoconnectivity as separate phenotypes throughout — in the transcriptomic, gradient, neurotransmitter, and cognitive-decoding analyses alike — and shows that they differ in molecular signature, hierarchical embedding, age profile, and cognitive association. This sign-resolved, multi-scale approach, and its reproducibility across developmental, sex, and severity subgroups, is the central conceptual and analytic advance relative to Berto et al.⁶ and is not addressed by treating connectivity alteration as a unitary construct.

Importantly, the principal contribution of the present study is not the independent observation of altered connectivity, cortical gradients, or transcriptomic associations, each of which has been reported previously in ASD ^36,44,6^. Rather, the findings demonstrate that these phenomena converge onto a common organizational framework spanning multiple levels of brain organization. Specifically, hyperconnectivity and hypoconnectivity exhibited partially distinct neurotransmitter-associated transcriptomic signatures, differential correspondence with cortical hierarchy, and distinct cognitive associations. These organizational relationships were preserved across developmental stages, sex-stratified analyses, and symptom-severity subgroups, suggesting generalizable principles of brain organization despite substantial ASD heterogeneity.

A central finding of this study is that hyperconnectivity and hypoconnectivity in ASD are not two poles of a single underlying process. Rather, they behave as two separable neurobiological entities, each with its own molecular signature, its own age-group profile, its own embedding within cortical hierarchy, and its own cognitive correspondence, and this dissociation was reproducible across age, sex, and symptom severity. Hyperconnectivity and hypoconnectivity differed in their spatial distribution, age-group profiles, and neurochemical signalling associated transcriptomic signatures. Hyperconnectivity preferentially involved higher-order cortical territories, including frontal, parietal, cingulate, and cerebellar regions, and was greater in the adult group than in the late-childhood group. In contrast, hypoconnectivity was relatively consistent across the age groups sampled and remained concentrated within subcortical and orbitofrontal systems. These findings help reconcile longstanding inconsistencies in the ASD connectivity literature by demonstrating that increased and decreased connectivity are not simply opposite manifestations of a single process but instead represent partially distinct patterns of large-scale brain organization ^76,71,37,62^. Importantly, the age-group comparisons indicate that hyperconnectivity and hypoconnectivity exhibit distinct age-related profiles, with relatively stable subcortical alterations coexisting alongside a more pronounced divergence of higher-order cortical systems in the older age groups. The observed age-group differences are consistent with emerging models of ASD emphasizing atypical maturation of association cortex and altered integration of distributed functional networks ^22,72,17^. Association cortices exhibit prolonged developmental trajectories and support integration across multiple cognitive systems, making them particularly sensitive to developmental variation ^69^. The greater prominence of hyperconnectivity within transmodal regions in the older age groups indicates a more pronounced differentiation of connectivity profiles within cortical systems commonly associated with social cognition, executive control, and adaptive behavior.

An additional finding was the consistent involvement of cerebellar systems across the functional connectivity, gradient, transcriptomic, and cognitive analyses. Cerebellar regions were among the most strongly hyperconnected territories and showed a more pronounced ASD–control divergence in the older age groups. Moreover, cerebellar systems repeatedly emerged in cognitive decoding analyses linked to social cognition, executive function, and perceptual processing. Although historically viewed primarily as a structure supporting motor coordination, the cerebellum is increasingly recognized as a critical component of distributed circuits supporting higher-order cognition and social behaviour.^9,73^. Disruptions in cerebro-cerebellar connectivity have been consistently reported in ASD.^24,20,67^, and the present findings extend this literature by demonstrating that cerebellar alterations show similar transcriptomic, hierarchical, and cognitive associations to those observed in cortical association systems. These observations suggest that cerebellar organization should be considered alongside cortical and subcortical systems when investigating large-scale brain organization in ASD.

At the molecular level, these findings indicate that ASD-related functional connectivity alterations are associated with distributed neurotransmitter and intracellular signalling systems rather than a single neurochemical pathway. Pathway enrichment analyses consistently implicated glutamatergic, GABAergic, dopaminergic, cholinergic, oxytocinergic, vasopressinergic, noradrenergic, and endocannabinoid pathways, together with calcium signalling, synaptic plasticity, circadian entrainment, neuroactive ligand–receptor interactions, endocrine signalling, retrograde endocannabinoid communication, and intracellular kinase cascades. These observations align with growing evidence that ASD-related brain alterations reflect distributed molecular signatures spanning multiple neurotransmitter systems rather than isolated abnormalities within a single pathway^13,14,6^. Accordingly, the present findings extend traditional excitation–inhibition imbalance models by demonstrating that ASD-related functional connectivity differences involve multiple interacting excitatory, inhibitory, neuromodulatory, and homeostatic signalling systems. Hyperconnectivity showed stronger associations with glutamatergic, oxytocinergic, and endocannabinoid pathways, whereas hypoconnectivity was relatively enriched for dopaminergic, noradrenergic, and cholinergic pathways. Although substantial overlap existed among neurotransmitter systems, these distinct molecular profiles suggest that hyperconnectivity and hypoconnectivity are not only spatially but also transcriptomically differentiated, extending previous imaging–transcriptomic studies in ASD^44,6^.

A particularly notable finding was the consistency of this molecular organization across developmental stage, sex, and symptom-severity subgroups: pathway-enrichment profiles, neurotransmitter-system contributions, and gradient-associated transcriptomic distributions remained stable despite variation in effect size, indicating that the molecular–hierarchical organization identified here is not an artifact of a particular demographic or clinical subgroup.

Importantly, supplementary analyses using the independent BrainSpan developmental transcriptomic atlas further supported the developmental applicability of the identified transcriptomic signatures. The hyperconnectivity- and hypoconnectivity-associated genes identified using the AHBA were broadly represented across childhood, adolescence, and adulthood, with the majority exhibiting relatively stable postnatal expression. Although BrainSpan represents normative developmental gene expression rather than ASD-specific molecular alterations, these findings suggest that the identified transcriptomic signatures are not restricted to adult cortical gene-expression patterns and remain developmentally represented across the age range encompassed by the ABIDE cohort.

The transcriptomic analyses further revealed strong cortical enrichment of ASD-associated molecular signatures. Across developmental, sex-stratified, and severity-defined analyses, genes associated with connectivity differences showed preferential expression within association cortices relative to subcortical structures. Default mode, frontoparietal, limbic, and other transmodal systems repeatedly emerged as regions exhibiting strong transcriptomic enrichment. These observations indicate that higher-order association cortex represents a region where transcriptomic, connectivity, gradient, and cognitive findings consistently con-verge. The repeated involvement of these systems across molecular, functional connectivity, gradient, and cognitive analyses supports the view that transmodal cortex occupies a central position within the organizational framework identified in the present study.

At the macroscale level, ASD was associated with systematic reorganization of cortical hierarchy. Functional gradient analyses revealed increased dispersion along the principal sensory-to-transmodal axis and altered positioning of higher-order association systems. De-fault mode, limbic, and frontoparietal networks exhibited the largest deviations, whereas primary sensory and visual systems showed comparatively smaller changes. These findings extend previous reports of altered cortical gradients in ASD ^36,57^ and support the view that atypical organization preferentially affects regions occupying the apex of cortical hierarchy, where information from multiple sensory, affective, and cognitive systems is integrated.

Developmental gradient analyses further revealed a greater separation between ASD and control participants in the older than in the younger age groups, particularly within transmodal territories. Together with the greater hyperconnectivity observed in the older age groups, these findings indicate a more pronounced divergence of cortical hierarchical organization in the older age groups. Such effects are consistent with theories proposing altered differentiation and segregation of large-scale cortical systems during neurodevelopment and with hierarchical models of cortical organization in which transmodal association cortices occupy the apex of information integration across the brain ^51,45,69^. More broadly, these results indicate that cortical hierarchy provides a useful intermediate scale for understanding how molecular variation corresponds with systems-level brain organization. Replication with longitudinal neuroimaging datasets is needed to confirm whether these age-group differences reflect genuine developmental trajectories.

Mapping transcriptomic signatures onto gradient space revealed an additional level of organization. Genes associated with hyperconnectivity showed stronger correspondence with transmodal cortical territories and greater clustering within higher-order regions of gradient space, whereas genes associated with hypoconnectivity were more diffusely distributed. These findings indicate that ASD-related transcriptomic signatures exhibit systematic spatial organization across cortical hierarchy rather than random distribution throughout the brain. This organizational pattern remained highly consistent across developmental stages, sex, and symptom severity, suggesting that it represents a reproducible feature of ASD-related brain organization.

Cognitive decoding and PLSC analyses provide an additional level of support for the identified molecular signatures. Meta-analytic decoding consistently implicated domains central to ASD, including social cognition, face processing, language, attention, visual perception, reward, and executive control ^37,47,71,66,19,80^. Furthermore, PLSC identified a significant latent axis linking neurotransmission-related transcriptomic signatures to social-cognitive, perceptual, attentional, and language-related functions. Importantly, these associations remained significant following BrainSMASH spatial-null correction, indicating that they cannot be explained solely by shared spatial organization. These findings demonstrate reproducible correspondence between transcriptomic organization and cognitive functional architecture and suggest that molecular signatures associated with ASD-related connectivity differences correspond to distributed cognitive systems supporting behaviorally relevant functions. Consistent involvement of regions such as the dorsolateral prefrontal cortex, fusiform gyrus, cuneus, striatum, and cerebellum further supports the relevance of the identified molecular signatures to social cognition, reward processing, perceptual integration, and executive control ^66,19,47^.

Several limitations should be considered when interpreting the present findings. First, transcriptomic analyses were based on the AHBA, which comprises transcriptomic profiles from six neurotypical adult donor brains and therefore represents a normative rather than disorder-specific molecular reference ^34^. This approach is widely adopted in imaging–transcriptomic studies investigating both healthy and disease-related brain organization ^60,10,3,46,36,6^, but it does not capture developmental or ASD-specific transcriptional variation. To partially address this limitation, we performed a supplementary sensitivity analysis using the independent BrainSpan developmental transcriptomic atlas. The majority of the identified hyperconnectivity- and hypoconnectivity-associated genes were broadly represented across childhood, adolescence, and adulthood and exhibited relatively stable postnatal expression, supporting the developmental applicability of the identified transcriptomic signatures across the age range represented in the ABIDE cohort. In addition, we performed a supplementary BrainSMASH spatial-null analysis to evaluate the potential influence of spatial autocorrelation on the multivariate imaging–transcriptomic model. The observed multivariate imaging–transcriptomic model exhibited significantly greater associations than expected under spatially constrained null models, providing additional support that the identified imaging–transcriptomic relationships were unlikely to be explained solely by intrinsic spatial autocorrelation ^11,30^. Nevertheless, BrainSpan represents normative developmental gene expression and therefore cannot account for ASD-specific transcriptional alterations, while the BrainSMASH analysis was performed as a supplementary model-level robustness assessment rather than a gene-level spatial-null inference. Future studies integrating age-matched, disorder-specific, single-cell, and spatial transcriptomic datasets together with gene-level spatial-null inference will further refine imaging–transcriptomic investigations of ASD. Second, the cross-sectional design of the ABIDE datasets precludes inference regarding longitudinal developmental trajectories of functional connectivity and transcriptomic organization. Third, despite harmonization using NeuroCombat, residual variability related to multisite image acquisition and scanner characteristics may remain. Finally, the observed imaging–transcriptomic associations are spatial and correlational in nature and therefore should not be interpreted as evidence of causal molecular mechanisms. Future studies integrating longitudinal neuroimaging, single-cell transcriptomics, and independent cohorts will be important for validating and extending the present findings.

A further methodological limitation concerns the assignment of AHBA transcriptomic samples to AAL3 regions, which was performed using nearest-neighbor matching of MNI coordinates rather than the abagen processing pipeline³, the current field-standard workflow for imaging–transcriptomic integration. We chose a volumetric, AAL3-based approach because AAL3 provides direct subcortical and cerebellar coverage, both of which showed prominent hyper- and hypoconnectivity in the present functional connectivity results and would be excluded or substantially under-sampled under the cortical-surface-restricted mapping conventions typical of abagen-based pipelines; however, this choice departs from the more common cortical-surface transcriptomic mapping convention in this literature and may not be directly comparable to surface-based pipelines in probe-to-region assignment or intensity-based structural filtering. Future work directly comparing volumetric nearest-neighbor and abagen-based cortical-surface mapping within the same dataset will help clarify the extent to which the present molecular–hierarchical findings depend on the specific gene-to-region mapping strategy employed.

## Methodology

### Data Acquisition and Preprocessing

Resting-state functional magnetic resonance imaging (rs-fMRI) data and accompanying phenotypic information were obtained from the Autism Brain Imaging Data Exchange (ABIDE) I and II initiatives, two large multicenter repositories established to facilitate reproducible neuroimaging studies of ASD ^17,18^. ABIDE I comprises 1,112 participants (539 individuals with ASD and 573 typically developing controls [CTL]) acquired across 17 international imaging sites, whereas ABIDE II includes 1,114 participants (521 ASD and 593 CTL) collected across 19 imaging sites. Together, these repositories constitute one of the largest publicly available rs-fMRI datasets for investigating large-scale functional brain organization in ASD. Following preprocessing, participants with complete rs-fMRI and phenotypic data required for subsequent analyses were retained, resulting in a final analytical cohort of 1,737 individuals (885 ASD and 852 CTL).

For ABIDE I, preprocessed imaging derivatives distributed through the Preprocessed Connectomes Project (PCP) were used. These derivatives were generated using the Configurable Pipeline for the Analysis of Connectomes (C-PAC), specifically the nofilt_noglobal preprocessing stream ^16^. Preprocessing included slice-timing correction, rigid-body head-motion correction, functional-to-anatomical coregistration, nonlinear normalization to the Montreal Neurological Institute (MNI152) 2-mm template, nuisance regression of 24 head-motion parameters together with white matter and cerebrospinal fluid signals, and spatial smoothing using a 6-mm full-width at half-maximum (FWHM) Gaussian kernel. Temporal band-pass filtering was intentionally omitted, and global signal regression was not performed to preserve low-frequency resting-state fluctuations while avoiding potential alterations in correlation structure associated with global signal removal ^53,59,65,79^.

Raw imaging data from ABIDE II were downloaded from the Functional Connectomes Project/International Neuroimaging Data-sharing Initiative (FCP/INDI) repository and pre-processed locally using C-PAC version 1.8 following the same nofilt noglobal workflow to ensure methodological consistency across both datasets ^16^.

Participant-level phenotypic information, including age, sex, diagnostic status (ASD versus CTL), imaging site, and Autism Diagnostic Observation Schedule (ADOS) scores (where available), was obtained from the PCP portal for ABIDE I and the INDI repository for ABIDE II.

Regional brain signals were extracted using the Automated Anatomical Labeling atlas version 3 (AAL3), comprising 170 anatomically defined regions of interest (ROIs), including 94 cortical, 22 subcortical, and 54 cerebellar regions ^61^. Compared with earlier versions of the AAL atlas, AAL3 provides improved anatomical delineation of cortical association areas, subcortical nuclei, and cerebellar structures, enabling comprehensive characterization of whole-brain functional organization. For these reasons, we chose to use AAL3 parcellation rather than ROIs defined by a functional atlas.

For each participant, the mean blood-oxygen-level-dependent (BOLD) time series was extracted from every AAL3 region. Functional connectivity between all pairs of brain regions was estimated using Pearson correlation coefficients, generating a symmetric 170 *×* 170 functional connectivity matrix for each participant ^7,28^. Correlation coefficients were subsequently transformed using Fisher’s *r*-to-*z* transformation,

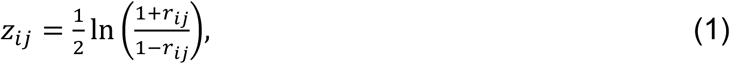

where *r_ij_* denotes the Pearson correlation coefficient between regions *i* and *j*.

To obtain a regional measure of large-scale functional integration, mean functional connectivity (node strength) was calculated for each brain region by averaging the Fisher-transformed connectivity values between that region and all remaining brain regions 64,

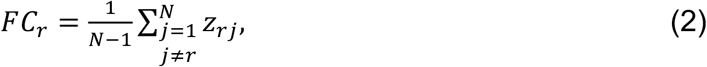

where *FC*_*r*_ denotes the regional mean functional connectivity (node strength) of region *r*, *z_rj_* is the Fisher-transformed connectivity between regions *r* and *j*, and *N* = 170 is the total number of AAL3 regions. These regional node-strength measures constituted the primary functional connectivity phenotype used throughout the study.

To minimize scanner- and site-related variability inherent to multisite neuroimaging datasets, regional node-strength measures were harmonized using NeuroCombat, an empirical Bayes harmonization framework that removes batch effects while preserving biologically meaningful variation associated with diagnosis and demographic variables ^40,26^. Imaging site was specified as the batch variable, whereas diagnosis, age, and sex were included as biological covariates. The harmonized regional functional connectivity measures were subsequently used in all downstream analyses.

To evaluate the generalizability of functional connectivity alterations across clinically relevant subgroups, participants were additionally stratified according to developmental stage, sex, and symptom severity. Developmental analyses included late childhood (6–12 years), adolescence (13–17 years), and adulthood (≥18 years). Symptom severity was assessed using total Autism Diagnostic Observation Schedule (ADOS) scores, a standardized observational measure of autism symptom severity, with participants stratified into lower (ADOS <8), intermediate (ADOS 8–11), and higher (ADOS ≥12) symptom-burden groups for comparative analyses. To ensure balanced comparisons and minimize bias arising from unequal subgroup sizes, participants in the larger subgroups were randomly subsampled to match the size of the smallest subgroup before downstream analyses. Random subsampling for balanced subgroup analyses was performed using a fixed random seed (random state = 42) to ensure reproducibility of participant selection.

### Identification of regional hyperconnectivity and hypoconnectivity

Regional functional connectivity alterations associated with ASD were quantified using the harmonized node-strength measures described above. For each AAL3 region, the mean regional functional connectivity was calculated separately for participants with ASD and typically developing controls (CTL). ASD-associated regional connectivity differences were then estimated as

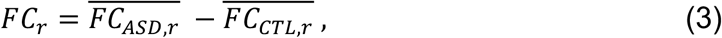

where 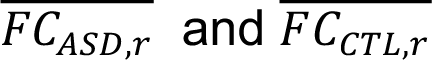 denote the mean regional node strength of brain region *r* in the ASD and CTL groups, respectively.

The resulting regional connectivity-difference map (Δ*FC*) served as the primary imaging phenotype throughout the study. Brain regions exhibiting positive connectivity differences (Δ*FC*_*r*_ > 0) were classified as hyperconnective, whereas regions exhibiting negative connectivity differences (Δ*FC*_*r*_ < 0) were classified as hypoconnective.

enabled characterization of the spatial distribution of increased and decreased functional connectivity across cortical, subcortical, and cerebellar systems.

To investigate the spatial organization of connectivity alterations, the regional ASD–CTL difference maps were projected onto the AAL3 anatomical atlas and visualized using three-dimensional cortical and cerebellar renderings. Region-wise connectivity differences were further summarized according to anatomical location and functional system to characterize the distribution of hyperconnective and hypoconnective regions across the brain.

The regional connectivity workflow was additionally applied to each predefined developmental, sex, and symptom-based subgroup. The resulting hyperconnectivity and hypoconnectivity maps served as the imaging phenotype for all subsequent transcriptomic, pathway enrichment, neurotransmitter-system, functional gradient, and cognitive decoding analyses.

### Imaging–transcriptomic analysis

To investigate the molecular correlates of regional functional connectivity alterations, regional gene-expression profiles were obtained from the AHBA, a comprehensive whole-brain transcriptomic resource containing genome-wide microarray measurements from six neurotypical adult donor brains ^34^. Imaging–transcriptomic analyses were performed following current recommendations for integrating spatial gene-expression data with neuroimaging phenotypes ^3,46,60^. AHBA tissue samples were transformed from their Montreal Neurological Institute (MNI) coordinates into AAL3 atlas voxel space and assigned to the corresponding anatomical parcels using voxel-based nearest-neighbour mapping. Samples located outside atlas boundaries were excluded. For parcels containing multiple tissue samples, probe-level expression values were averaged to obtain a single regional estimate. Probe identifiers were subsequently mapped to official gene symbols using the AHBA probe annotation file, and expression values from multiple probes corresponding to the same gene were averaged, yielding a parcel-wise regional gene-expression matrix aligned with the functional connectivity data.

The regional transcriptomic data was organized into a predictor matrix

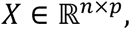

where *n* denotes the number of AAL3 brain regions and *p* represents the number of mapped genes. The corresponding imaging phenotype consisted of the regional ASD-associated functional connectivity differences,

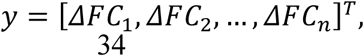

where Δ*FC_r_* denotes the regional functional connectivity difference defined in Eq. (3). Prior to multivariate analysis, gene-expression values were standardized using z-score normalization,

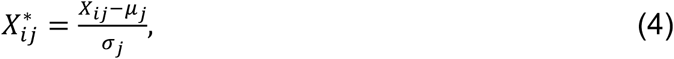

where *X_ij_* denotes the expression level of gene *j* within brain region *i*, and *µ_j_* and *σ_j_* represent the corresponding mean and standard deviation across regions.

Associations between regional transcriptomic organization and functional connectivity alterations were identified using Partial Least Squares Regression (PLSR), a supervised multivariate approach that identifies latent variables maximizing covariance between high-dimensional predictor variables and a continuous imaging phenotype while accounting for substantial multicollinearity among predictors ^48,49,43^. PLS-based approaches have become widely used in imaging–transcriptomic studies because they provide robust identification of molecular gradients associated with macroscale neuroimaging phenotypes ^60,3,10^.

The PLSR model can be expressed as

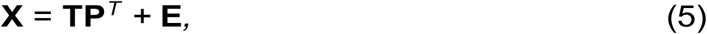

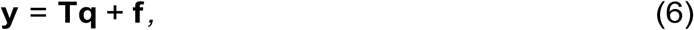

where **T** denotes the latent score matrix, **P** represents the gene loading matrix, **q** contains regression coefficients relating the latent variables to the imaging phenotype, and **E** and **f** denote residual error terms.

PLSR was implemented using two latent components. Regression coefficients were extracted for every gene and used to rank genes according to their spatial association with regional functional connectivity alterations. Genes with positive regression coefficients were interpreted as exhibiting positive spatial associations with regions showing relatively greater functional connectivity differences, whereas genes with negative coefficients exhibited inverse spatial associations. These ranked gene lists were used for all subsequent downstream analyses.

To independently evaluate the robustness of the multivariate findings, non-parametric Spearman rank correlation coefficients were additionally computed between regional gene-expression profiles and regional functional connectivity measures for every gene. Spearman correlation was selected because it is robust to non-Gaussian distributions and monotonic relationships commonly encountered in spatial transcriptomic datasets.

For each gene,

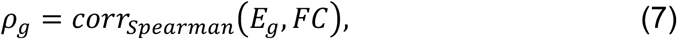

where *E_g_* denotes the regional expression profile of gene *g*, and *FC* represents the regional functional connectivity phenotype.

Differences between ASD and control gene–connectivity associations were quantified using Fisher’s r-to-z transformation ^25^,

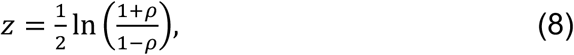

followed by

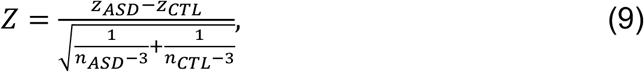

where *n*_*ASD*_ and *n*_*CTL*_ denote the numbers of observations contributing to the ASD and control correlation estimates, respectively. The resulting Z-statistics quantified differences in gene–connectivity associations between diagnostic groups. Genes demonstrating concordant evidence across both analyses were considered robust transcriptomic correlates and retained for downstream analyses.

To account for spatial autocorrelation, supplementary BrainSMASH spatial-null analyses were performed¹². BrainSMASH generated 10,000 spatially constrained surrogate functional connectivity maps preserving the spatial autocorrelation of the observed phenotype. PLSR was repeated for each surrogate map, and the observed latent components were compared with the corresponding null distributions to evaluate whether imaging–transcriptomic associations exceeded those expected under spatially constrained null models³⁰.

### Pathway enrichment and neurotransmitter-system characterization

To identify biological pathways associated with ASD-related functional connectivity alterations, genes were ranked according to their Partial Least Squares Regression (PLSR) coefficients and analysed using pre-ranked Gene Set Enrichment Analysis (GSEA) implemented in the GSEApy Python package against the KEGG 2021 Human pathway database⁶⁸^˒^²³^˒^⁴²^˒^⁴¹. Positive PLSR coefficients indicated preferential association with hyperconnectivity, whereas negative coefficients indicated preferential association with hypoconnectivity. Enrichment significance was assessed using 1,000 phenotype permutations (random seed = 42).

For each KEGG pathway *S*, the enrichment score (ES) was calculated using the weighted running-sum statistic proposed by Subramanian et al. ^68^,

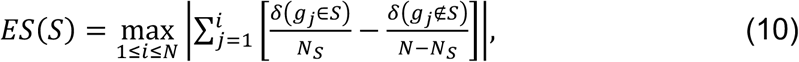

where *N* denotes the total number of ranked genes, *N_S_* represents the number of genes belonging to pathway *S*, and *δ*(*·*) is an indicator function.

To facilitate comparison across pathways of different sizes, enrichment scores were normalized to generate normalized enrichment scores (NES),

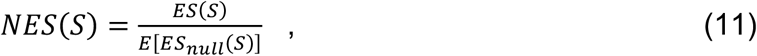

where *E*[*ES*_null_] denotes the expected enrichment score obtained from permutation-derived null distributions. Nominal permutation *p*-values were computed for every pathway, and multiple comparisons were controlled using the Benjamini–Hochberg false discovery rate (FDR) procedure ^5^. Pathways with FDR-adjusted *P_FDR_ <* 0.05 were considered statistically significant.

Leading-edge genes were extracted from all significant pathways, and the proportion of leading-edge genes within each pathway (gene ratio) was calculated to summarize pathway representation. Genes were subsequently classified into neurochemical systems (including neurotransmitters, neuropeptides and neuromodulatory signalling systems) using curated KEGG pathway annotations and established neurochemical terminology ^41,70^. Genes participating in multiple signalling pathways were assigned to all supported categories. Neurochemical system representation was quantified as the proportion of annotated leading-edge genes assigned to each system, and preferential associations with hyperconnectivity or hypoconnectivity were evaluated using the mean differential Fisher *Z*-statistic across genes within each category.

To investigate relationships among enriched biological pathways, a binary gene–pathway membership matrix was constructed and pairwise pathway similarity was quantified using Pearson correlation. Pathway dissimilarity was defined as 1 − Pearson correlation, and average-linkage hierarchical clustering was performed to identify biologically related pathway modules ^21,54^. Clustered heatmaps were generated to visualize coordinated signalling modules and shared gene composition across neurotransmitter-associated pathways.

### Functional gradient analysis

Cortical functional gradients were estimated using diffusion map embedding implemented in BrainSpace toolbox ^77,15,45,38^. For each participant, the regional functional connectivity matrix derived from the AAL3 atlas was used to estimate cortical functional gradients. Pairwise similarity between regional connectivity profiles was quantified using cosine similarity,

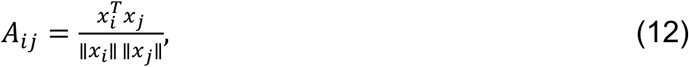

where *x*_*i*_ and *x*_j_denote the functional connectivity profiles of brain regions *i* and *j*, respectively. The resulting affinity matrix was subsequently used as input to diffusion map embedding, which estimates the eigenvectors of the normalized diffusion operator,

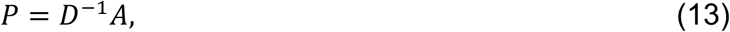

where *A* denotes the affinity matrix and *D* is the corresponding diagonal degree matrix with elements

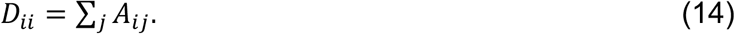

The eigenvectors of the diffusion operator satisfy

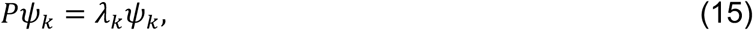

where λ_*k*_ and ψ_*k*_ denote the eigenvalue and eigenvector corresponding to gradient *k*. Gradients were ordered according to the proportion of variance explained by their corresponding eigenvalues. Only the first two gradients were retained for subsequent analyses. Gradient 1 corresponded to the principal sensory-to-transmodal cortical axis, whereas Gradient 2 represented a secondary organizational axis ^45,38,36^.

Group-average gradient maps were subsequently computed separately for ASD and typically developing controls (CTL). To facilitate direct comparison between groups, ASD gradients were aligned to the control reference space using Procrustes alignment as implemented in BrainSpace ^77^. Regional gradient differences were calculated as

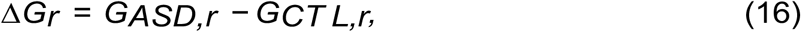

where *G_ASD,r_* and *G_CT L,r_* denote the gradient score of brain region *r* in the ASD and CTL groups, respectively.

To quantify alterations in cortical hierarchy, the distribution of Gradient 1 values was estimated using kernel density estimation, following previous studies investigating cortical hierarchy compression and expansion in neurodevelopmental disorders ^36,78^. Network-level analyses were performed by averaging regional gradient scores within canonical functional systems, including the default mode, frontoparietal, salience, limbic, visual, sensorimotor, cerebellar, and subcortical networks. Group differences in network-level gradient organization were subsequently quantified to identify systems exhibiting preferential alterations in cortical hierarchy.

To investigate the molecular embedding of cortical hierarchy, transcriptomic signatures identified from the imaging–transcriptomic analyses were projected onto gradient space according to their regional expression profiles obtained from AHBA. Hyperconnectivity-associated and hypoconnectivity-associated genes were analysed separately to determine whether their spatial distributions preferentially occupied distinct positions along the principal cortical gradient and to evaluate the relationship between molecular organization and large-scale cortical hierarchy.

### Cognitive decoding using Neurosynth

Meta-analytic functional decoding was performed using the Neurosynth platform to identify cognitive functions associated with ASD-related transcriptomic signatures ^80^. Regional gene-expression profiles were integrated with AAL3-parcellated Neurosynth cognitive activation maps. Genes were grouped according to neurotransmitter systems identified by pathway enrichment analyses ^6,35^ and cognitive functions associated with each neurotransmitter system were identified by comparing regional gene-expression profiles with cognitive activation maps. Cognitive terms were ranked by association strength and compared across neurotransmitter systems.

### Transcriptomic–cognitive association using Partial Least Squares Correlation

To investigate the relationship between regional transcriptomic organization and cognitive functional architecture, Partial Least Squares Correlation (PLSC) was performed between regional gene-expression profiles and parcel-wise Neurosynth cognitive activation maps ^48,49,43^. PLSC identifies latent variables that maximize the covariance between two high-dimensional datasets while accounting for multicollinearity among predictors.

The predictor matrix,

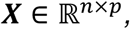

contained regional gene-expression profiles obtained from the Allen Human Brain Atlas, whereas the response matrix,

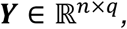

contained parcel-wise Neurosynth cognitive activation scores, where *n* denotes the number of AAL3 regions, *p* the number of genes, and *q* the number of cognitive terms. PLSC identifies latent variables that maximize the covariance between linear combinations of the two datasets,

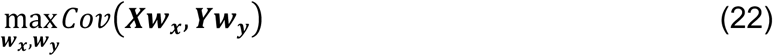

where **w***_x_* and **w***_y_* denote the gene and cognitive loading vectors, respectively.

The corresponding latent variables were calculated as

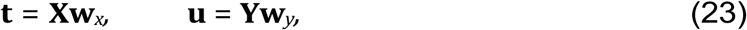

where **t** and **u** represent the transcriptomic and cognitive scores associated with each latent component.

Statistical significance of each latent variable was assessed using permutation testing with 10,000 permutations, in which the rows of one data matrix were randomly permuted to generate a null distribution of covariance values ^49,43^. Latent variables with empirical permutation *P <* 0.05 were considered statistically significant.

To account for spatial autocorrelation inherent in transcriptomic and neuroimaging data, significance was additionally evaluated using BrainSMASH spatially constrained null models^11^. A total of 10,000 surrogate maps preserving the empirical spatial autocorrelation structure were generated and observed covariance values were compared with the corresponding spatial-null distributions.

The stability of gene and cognitive-term loadings was assessed using 10,000 bootstrap resamples ^49^. Bootstrap ratios were calculated as

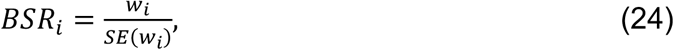

where *w*_*i*_denotes the loading of gene or cognitive term *i* and *SE*(*w*_*i*_) is bootstrap-estimated standard error. Genes and cognitive terms were ranked according to their bootstrap ratios, with positive and negative loadings interpreted separately to identify transcriptomic signatures associated with distinct cognitive domains. Genes with the largest absolute bootstrap ratios were subsequently annotated according to neurotransmitter-system classifications derived from pathway enrichment analyses, enabling interpretation across neurotransmitter systems.

The integration of permutation testing, BrainSMASH spatial-null modelling, bootstrap stability analysis, and neurotransmitter-system annotation provided a robust multiscale framework for relating regional gene-expression patterns to distributed cognitive functional architecture while accounting for statistical uncertainty and spatial autocorrelation.

### Robustness and sensitivity analysis

To evaluate the robustness and generalizability of the identified imaging–transcriptomic associations, supplementary robustness and sensitivity analyses were performed. First, a leave-one-site-out (LOSO) analysis assessed the influence of site-specific variability within the multisite ABIDE cohort. In each iteration, one imaging site was excluded, regional ASD–control functional connectivity differences were recomputed, and the complete imaging–transcriptomic pipeline, including PLSR, was repeated. Robustness was evaluated by determining whether hyperconnectivity-associated genes retained positive PLS regression coefficients and hypoconnectivity-associated genes retained negative coefficients across iterations.

Second, a leave-one-region-out (LORO) analysis evaluated the influence of individual brain regions. Only AAL3 regions exhibiting significant ASD-related functional connectivity differences (*P* < 0.05) in the primary analysis were considered. Each region was sequentially excluded, and the imaging–transcriptomic analysis was repeated. Stability was assessed by determining whether genes retained the same direction observed in the primary analysis.

To assess the developmental applicability of the AHBA-derived transcriptomic signatures, a supplementary sensitivity analysis was performed using the independent BrainSpan developmental transcriptomic atlas spanning prenatal development through adulthood^52^. The top 20 hyperconnectivity-associated and top 20 hypoconnectivity-associated genes identified by PLSR were evaluated across childhood, adolescence, and adulthood. Developmental stability was quantified using the coefficient of variation and adult-to-childhood fold change in mean expression. Genes exhibiting less than two-fold variation were classified as developmentally stable, and expression trajectories were categorized as increasing, stable, or decreasing. Differences between hyperconnectivity- and hypoconnectivity-associated gene sets were evaluated using Fisher’s exact test to determine whether the AHBA-derived signatures remained broadly represented across the developmental age range encompassed by the ABIDE cohort.

### Statistical analysis

All statistical analyses were performed using Python (version 3.11). Data processing and numerical analyses were conducted using NumPy³³, SciPy⁷⁵, and Pandas⁵⁰. Neuroimaging analyses were performed using Nilearn¹, BrainSpace⁷⁷, and NiBabel⁸. Imaging–transcriptomic analyses were implemented using scikit-learn⁵⁸, pathway enrichment analyses using GSEApy²³, spatial-null analyses using BrainSMASH¹², and data visualization using Matplotlib³⁹ and Seaborn. Regional group differences were assessed using two-sided independent-samples Student’s t-tests.

Unless otherwise stated, statistical significance was defined as a two-sided *P* < 0.05. For analyses involving multiple comparisons, *P*-values were adjusted using the Benjamini–Hochberg false discovery rate (FDR) procedure⁵, with FDR-adjusted *Pfdr* < 0.05 considered statistically significant.

Statistical significance of latent variables identified by Partial Least Squares Correlation (PLSC) was assessed using 10,000 permutation tests, and the stability of gene and cognitive-term loadings was evaluated using 10,000 bootstrap resamples⁴⁹. To account for spatial autocorrelation, statistical significance of the PLSC and supplementary PLSR analyses was additionally evaluated using BrainSMASH spatially constrained null models with 10,000 surrogate maps¹²^˒^³⁰.

Because the age, sex, and severity subgroup analyses each generated an independent battery of pathway, gradient, and gene-expression tests, we additionally considered the multiple-comparisons burden across the full set of subgroup analyses rather than within each family alone. Within every individual analysis (pathway enrichment, gene-level correlations, gradient contrasts, and PLSC loadings), Benjamini–Hochberg FDR correction was applied as described above. Across subgroup analyses, we relied on convergence rather than significance counting as the primary evidence standard: a molecular or hierarchical feature was considered reproducible only if it emerged consistently, and in the same direction, across independently FDR-corrected age, sex, and severity strata, a considerably more conservative criterion than treating each subgroup test as an independent test of the same hypothesis. We note that this convergence criterion does not constitute a formal correction for the total number of hypothesis tests performed across the entire subgroup battery, and some nominally significant subgroup-specific effects may not survive a study-wide correction; such effects are interpreted with corresponding caution in the Results and Discussion.

## Data availability

The neuroimaging data analysed in this study were obtained from the Autism Brain Imaging Data Exchange (ABIDE) I and II repositories, which are publicly available through the International Neuroimaging Data-sharing Initiative (INDI) at http://fcon_1000.projects.nitrc.org/indi/abide/.

Human brain gene expression data were obtained from the Allen Human Brain Atlas (AHBA), provided by the Allen Institute for Brain Science, and are publicly available at https://human.brain-map.org/.

Developmental transcriptomic data used for the supplementary sensitivity analysis were obtained from the BrainSpan Atlas of the Developing Human Brain and are publicly available at https://www.brainspan.org/.

No new data were generated in this study. All data supporting the findings are publicly available from the sources listed above.

## Code availability

The custom code used for data preprocessing and analysis in this study is publicly available at https://github.com/AbinayaVairam0903/neuroomics-asd.

## Author contributions statement

Abinaya and DR were involved in primary study design and in planning the conceptual analysis. DR was primarily responsible for supervising and funding this study; BM provided methodological guidance on the imaging-transcriptomic and network-based analyses and critically reviewed the manuscript; Abinaya V, Km Bhavna, LQU, BM, and DR all contributed to writing and editing the original research article.

## Acknowledgements

This research was supported in part by grants from the SERB Core Research Grant (CRG) S/SERB/DPR/20230033 extramural grant from the Department of Science and Technology, Ministry of Science and Technology, Govt. India and we acknowledge the computational support from IIT Jodhpur core research facility.

## 1 Supplementary Materials

### 1.1 Gene Expression and Neurotransmitter Analysis

**Figure S1:**
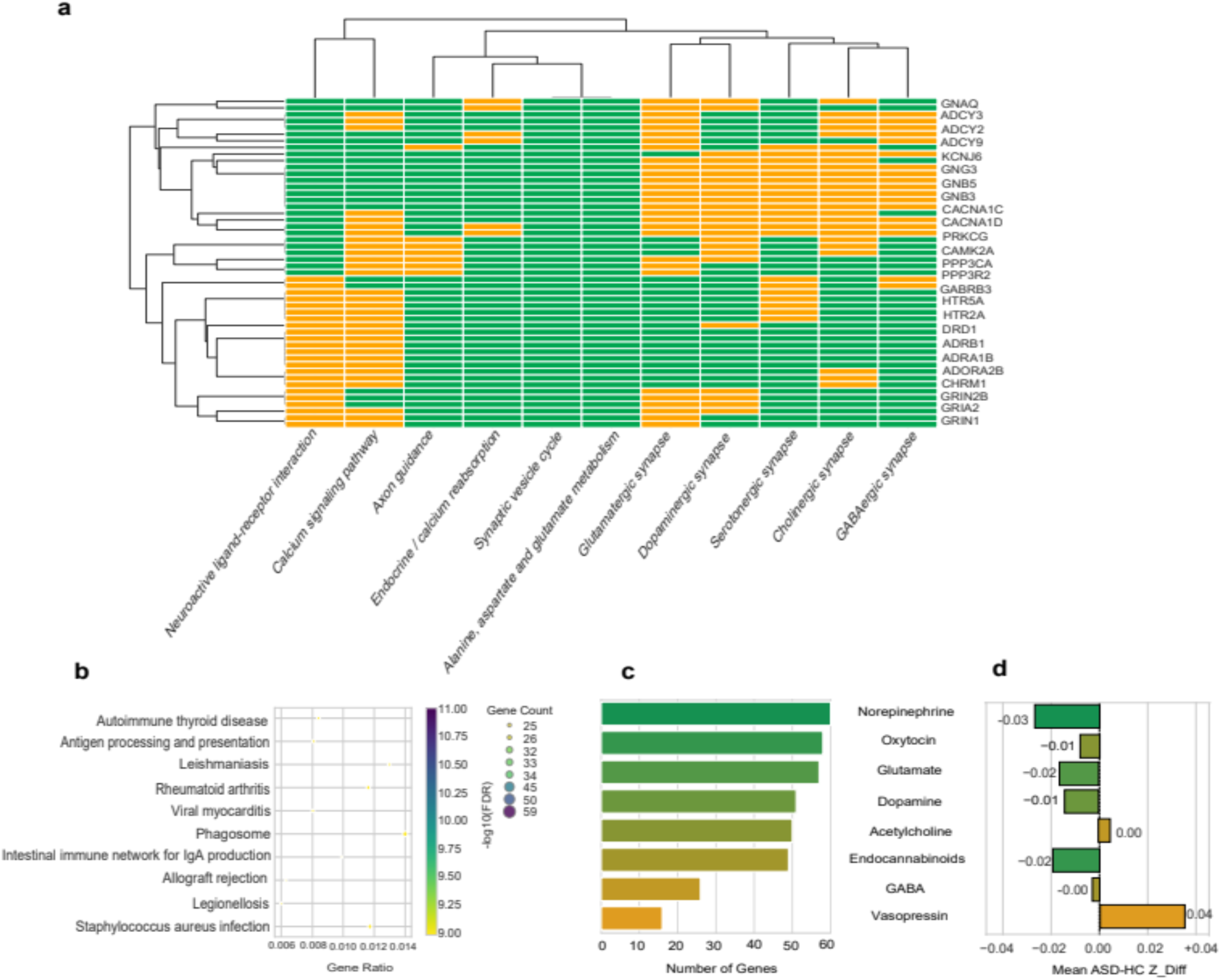
Neurotransmitter-related gene expression patterns in late childhood ASD, showing pathway enrichment, regional expression differences, and neurotransmitter-associated molecular alterations.

**Figure S2:**
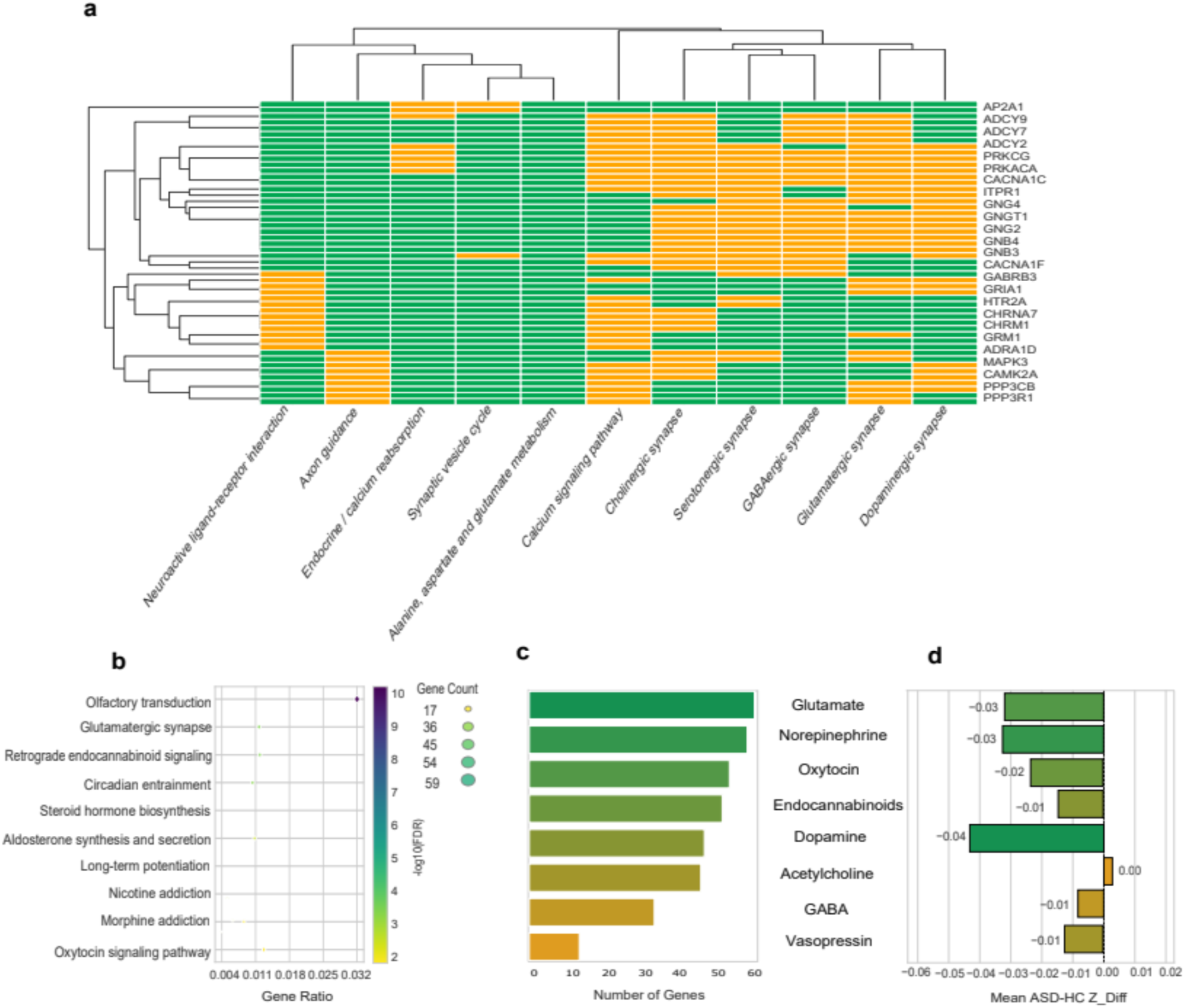
Neurotransmitter-related gene expression patterns in adolescent ASD, highlighting developmental alterations in synaptic and signaling pathways.

**Figure S3:**
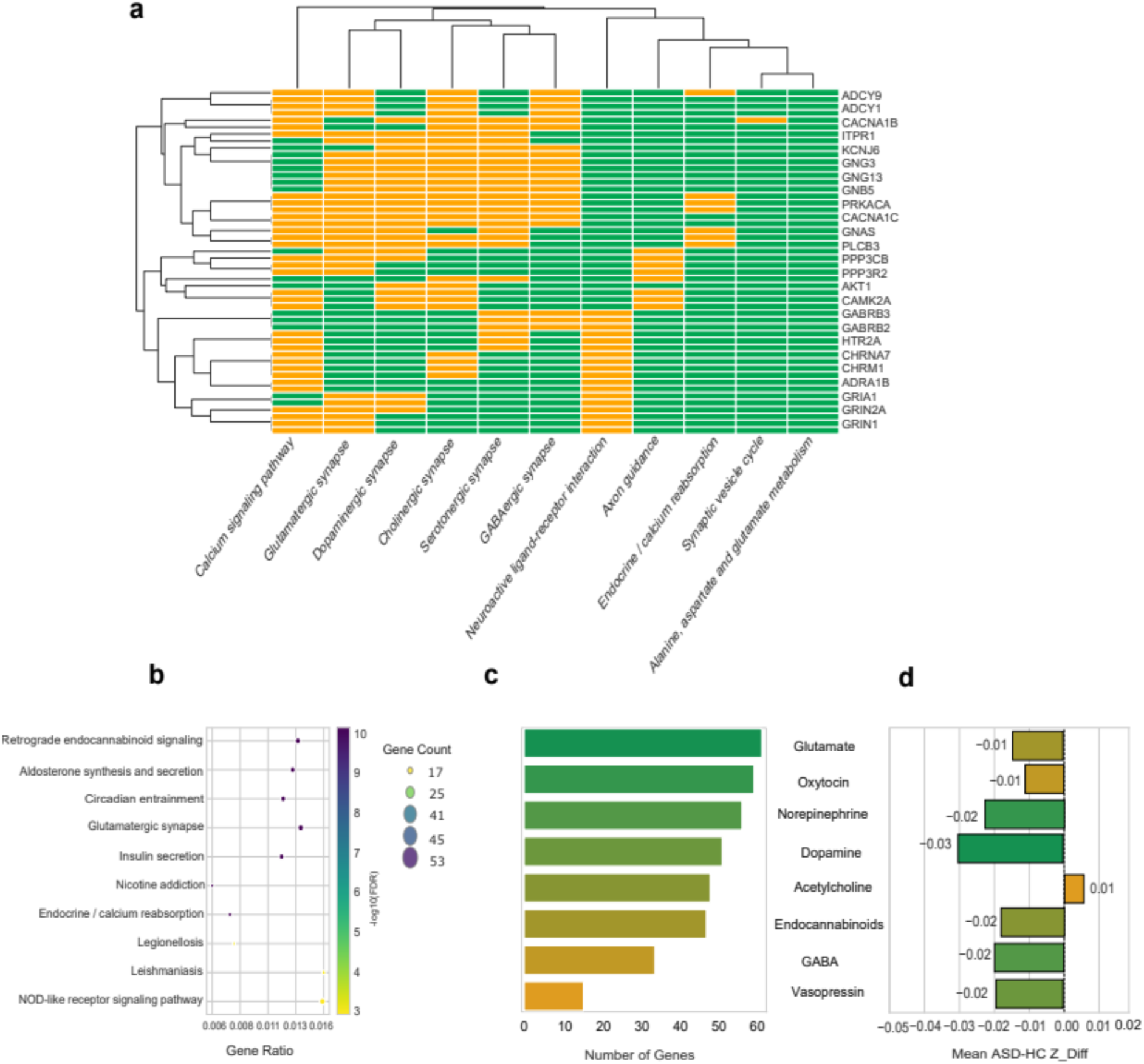
Neurotransmitter-related gene expression patterns in adult ASD, showing persistent and stage-specific molecular pathway alterations.

**Figure S4:**
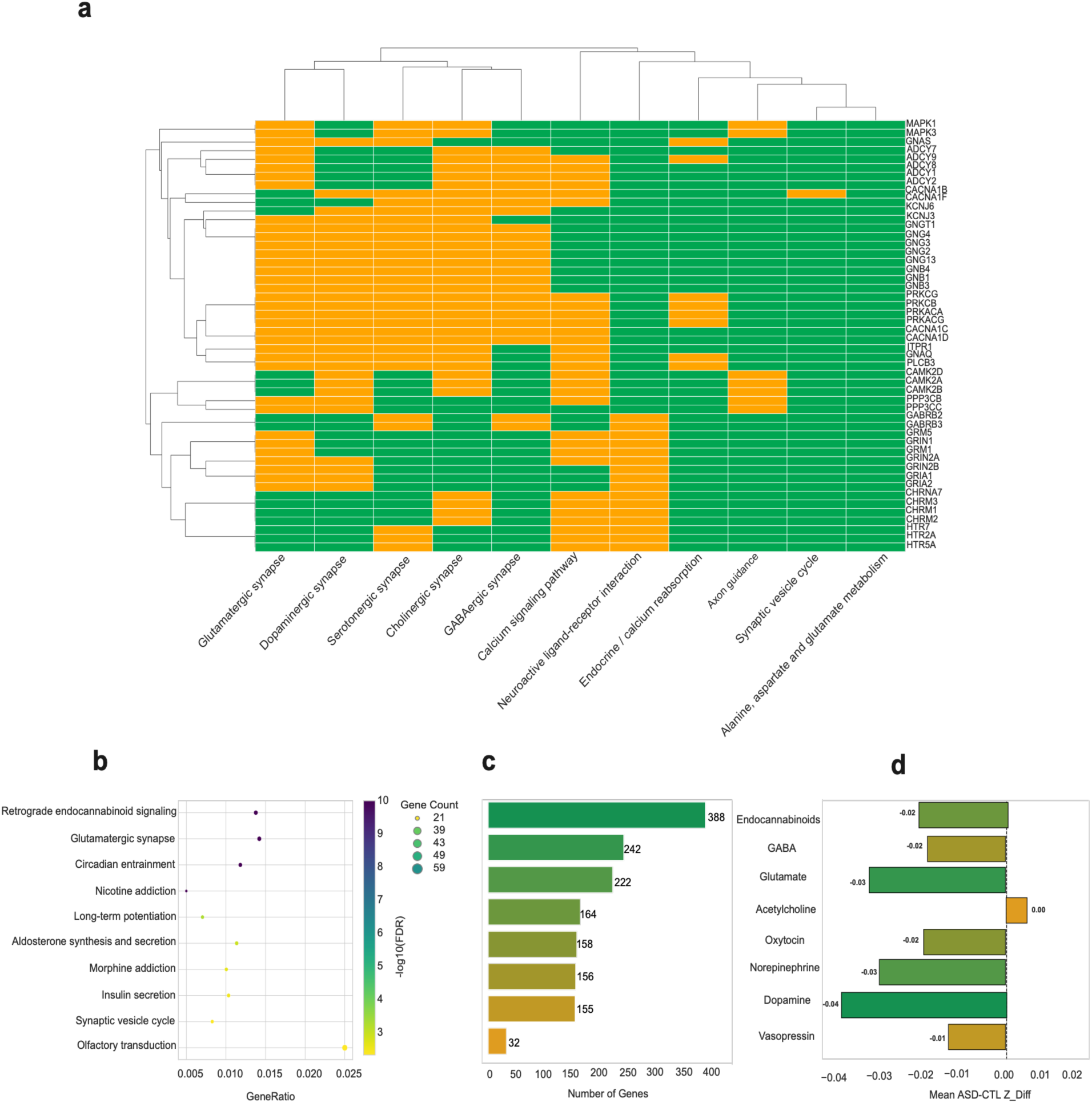
Neurotransmitter-related gene expression patterns in male ASD participants, showing altered synaptic signaling pathways and neurotransmitter-associated molecular expression profiles.

**Figure S5:**
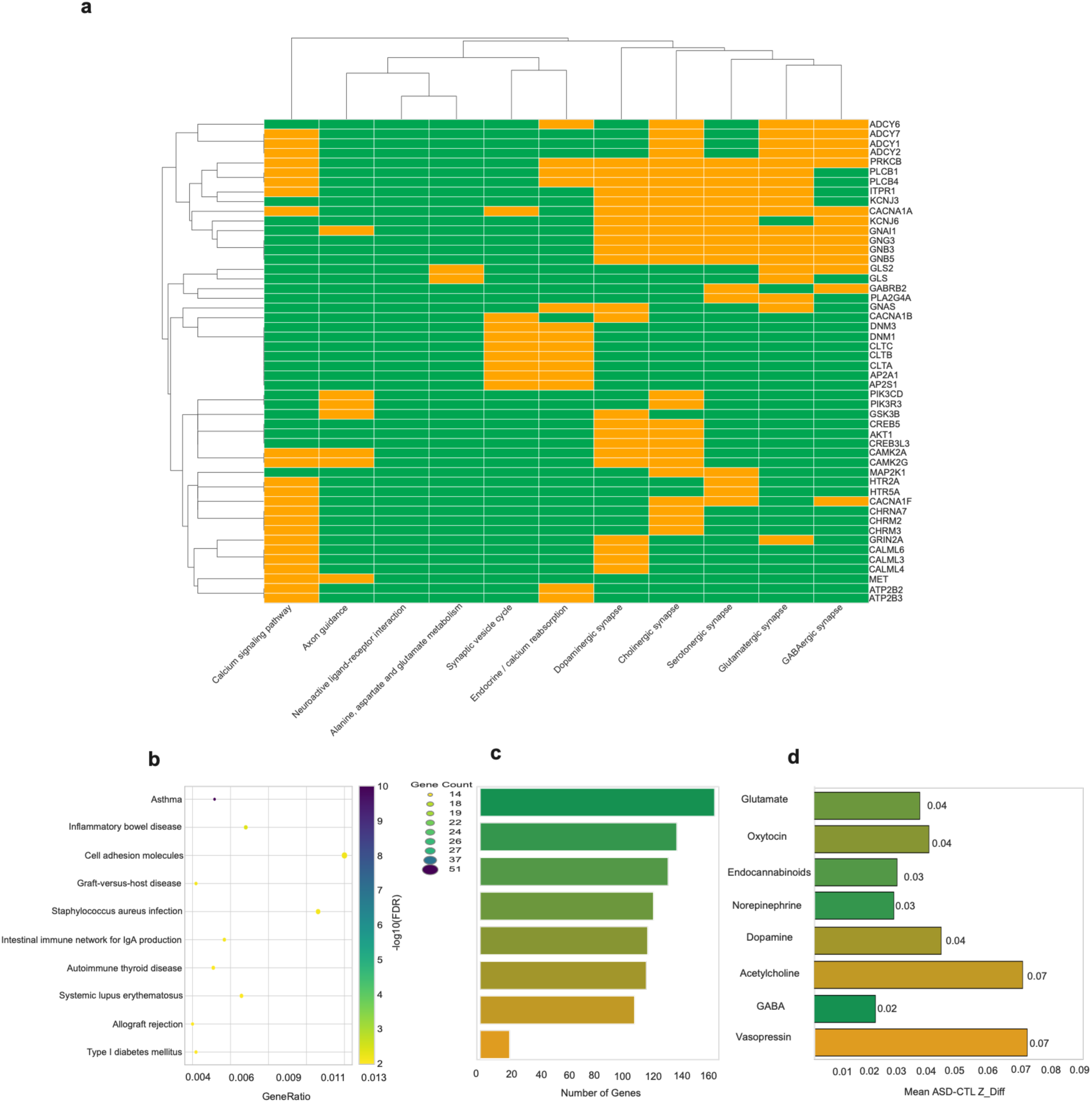
Neurotransmitter-related gene expression patterns in female ASD participants, highlighting sex-specific molecular and neurotransmitter-associated alterations.

**Figure S6:**
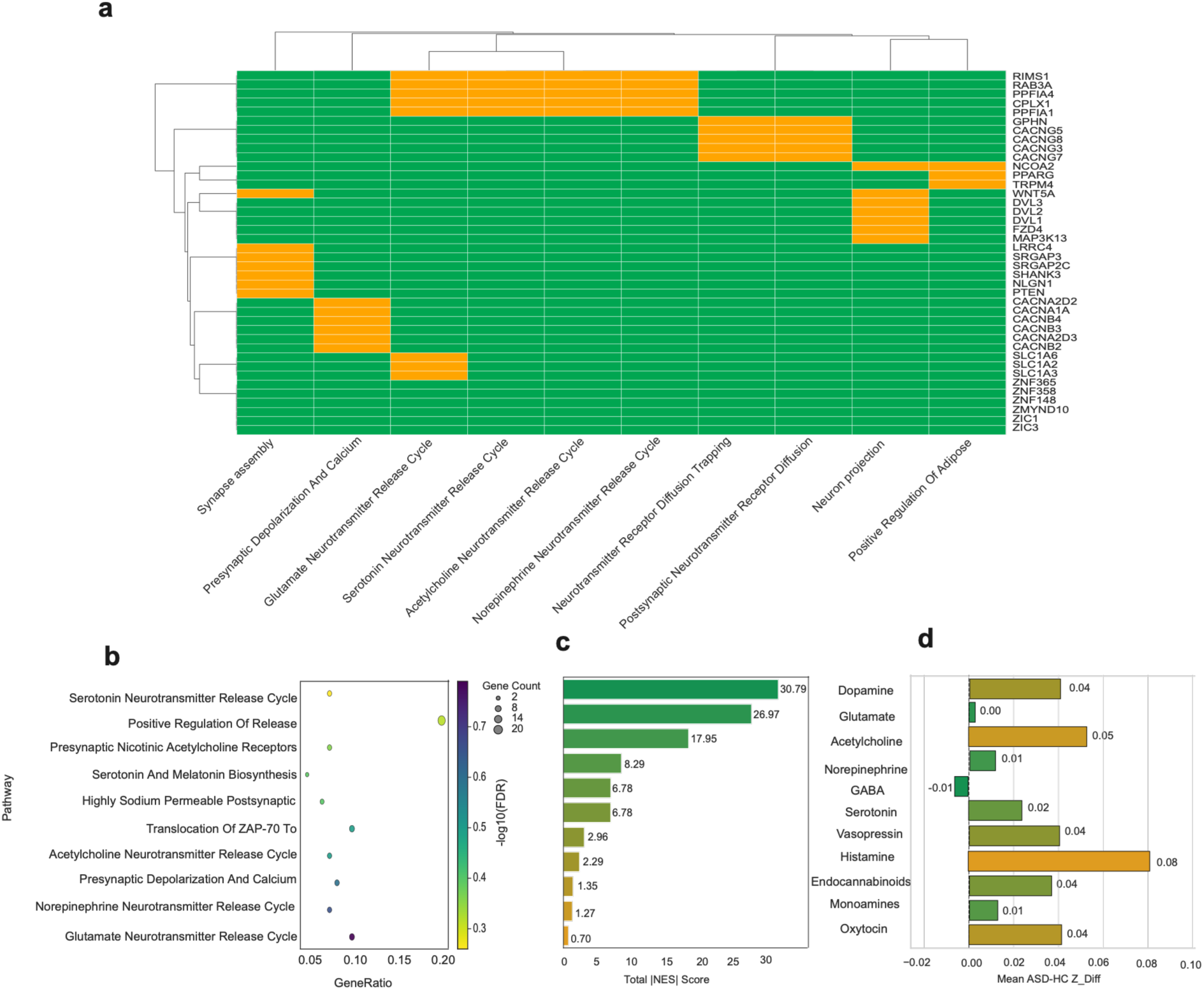
Neurotransmitter-related gene expression patterns in the low symptom group, showing pathway enrichment and neurotransmitter-associated molecular changes linked with lower symptom severity.

**Figure S7:**
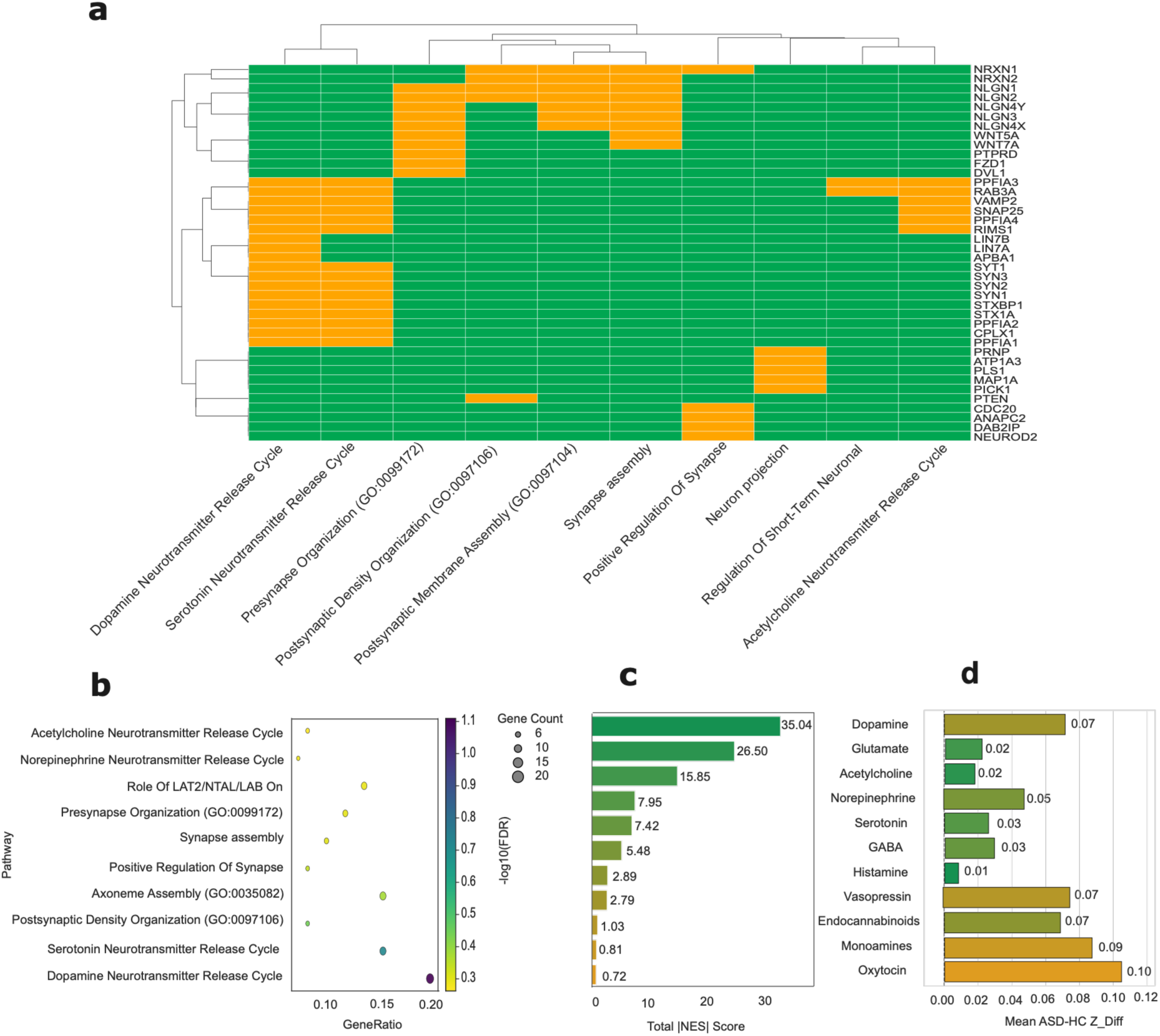
Neurotransmitter-related gene expression patterns in the intermediate symptom group, demonstrating alterations in synaptic signaling and neurotransmitter-associated pathways.

**Figure S8:**
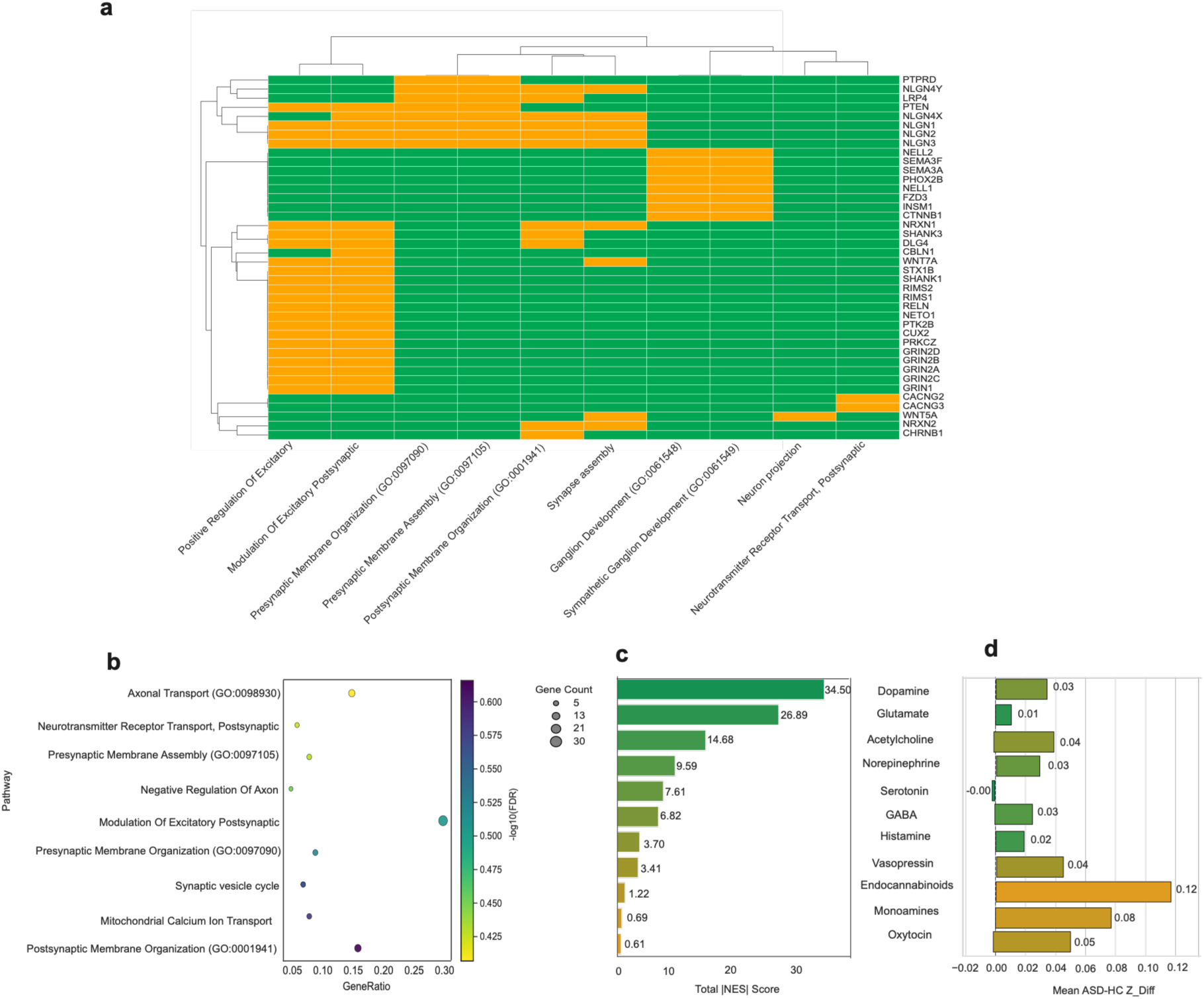
Neurotransmitter-related gene expression patterns in the high symptom group, showing pathway enrichment and neurotransmitter-associated molecular alterations linked with higher symptom severity..

### 1.2 Hyper- and Hypo-Connectivity Analysis

**Figure S9:**
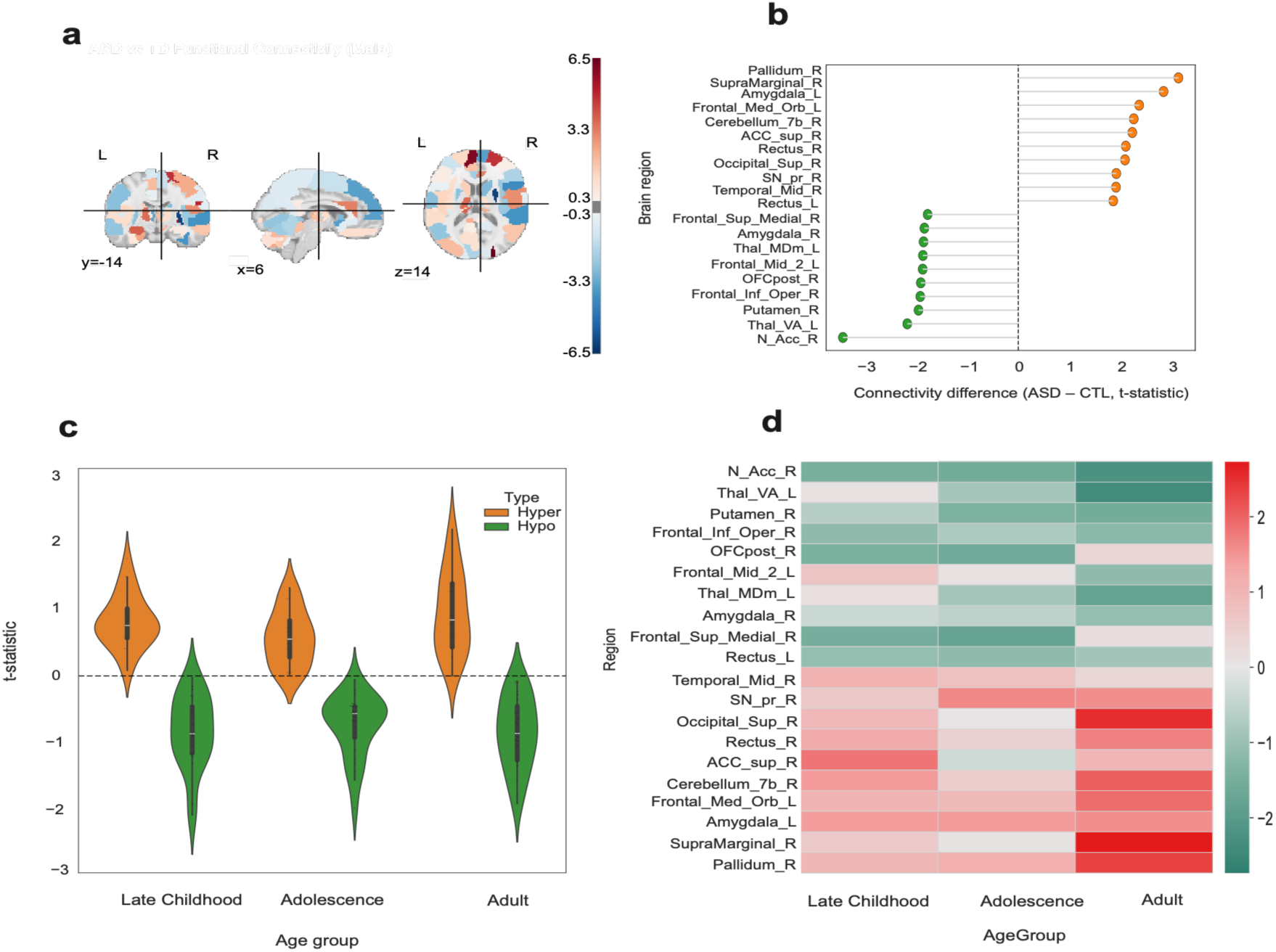
Functional hyper- and hypo-connectivity patterns in male ASD participants, showing sex-specific alterations across cortical and subcortical networks.

**Figure S10:**
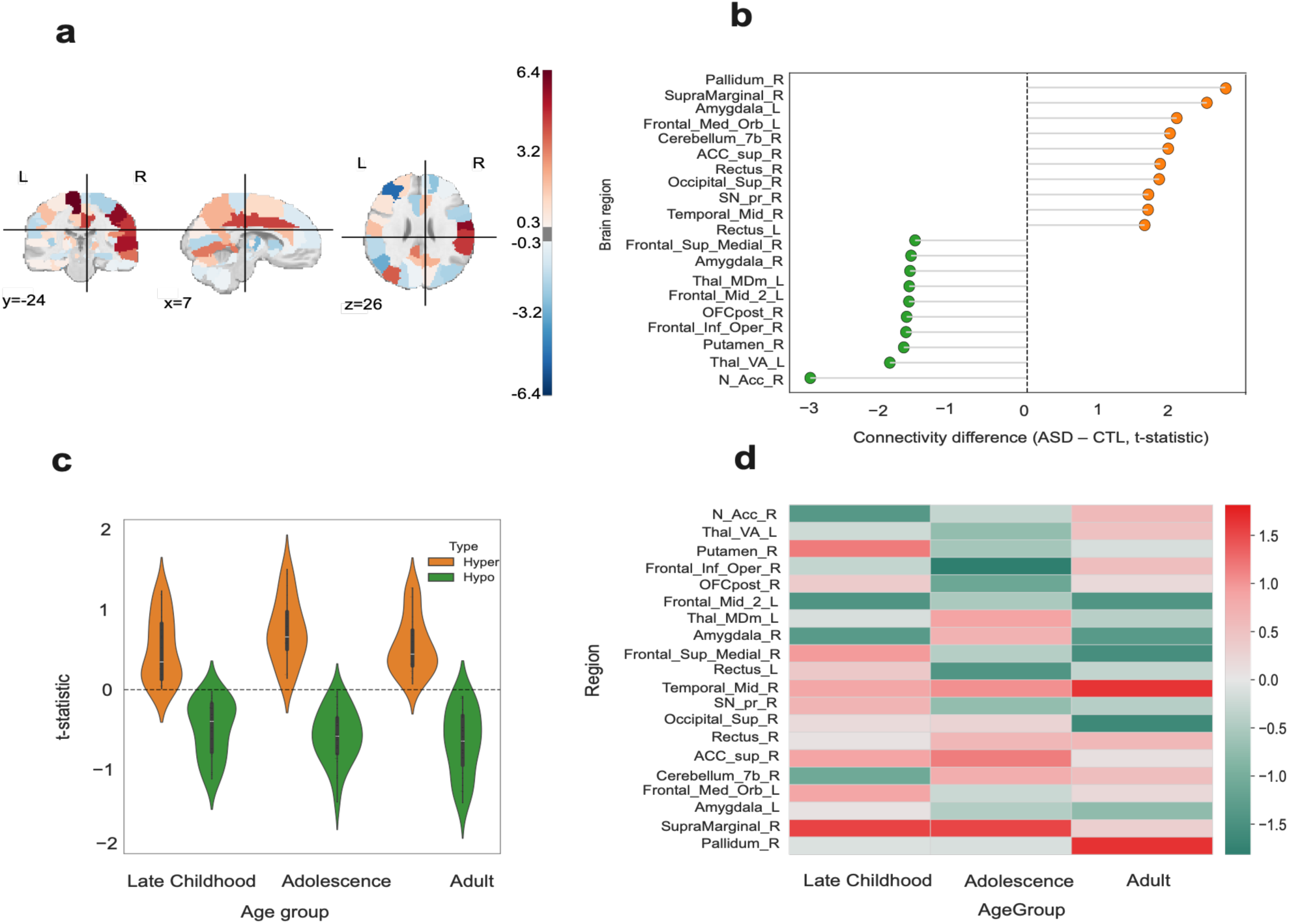
Functional hyper- and hypo-connectivity patterns in female ASD participants, highlighting sex-related differences in regional brain connectivity.

**Figure S11:**
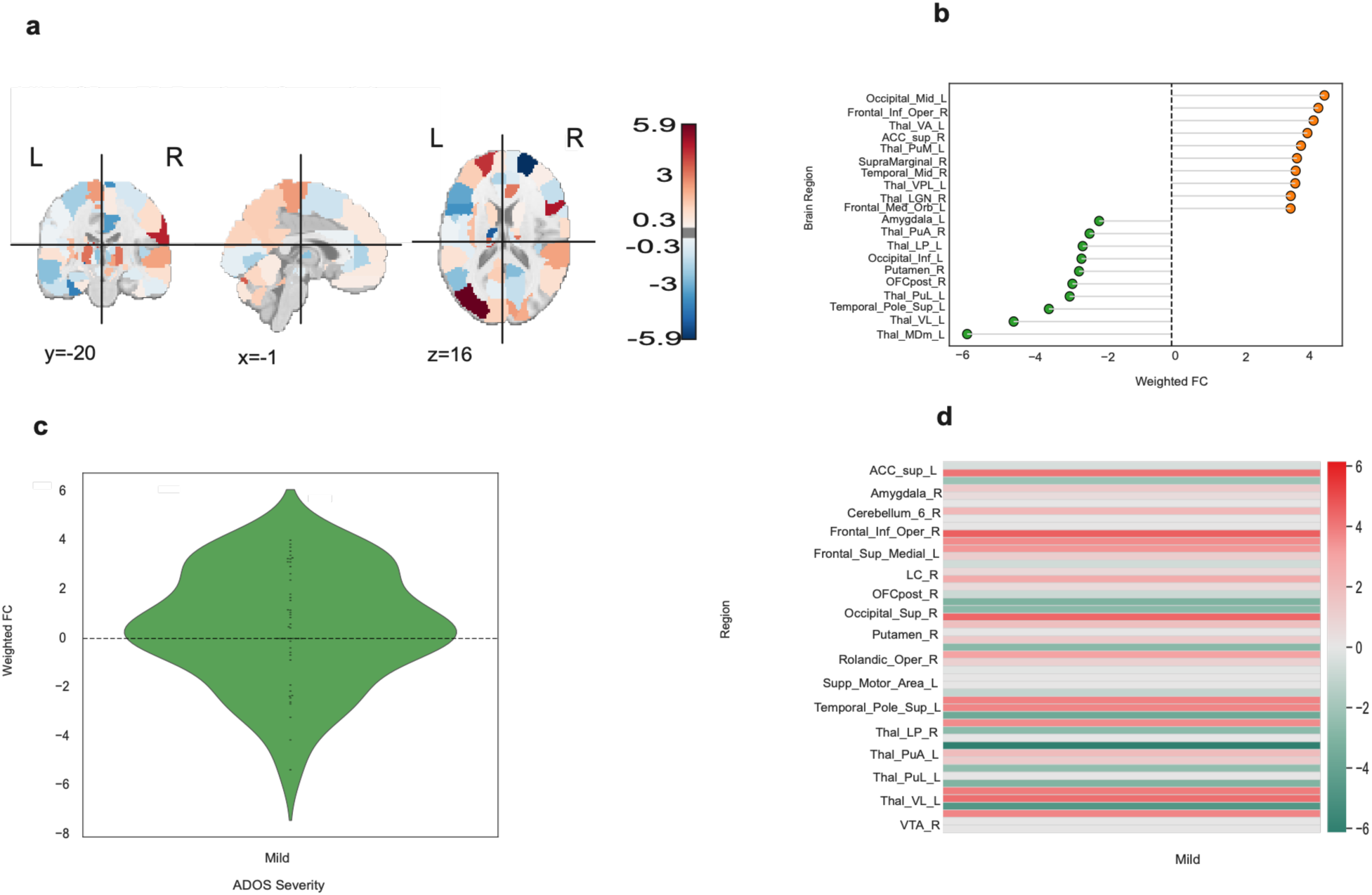
Functional hyper- and hypo-connectivity patterns in low symptom group, showing connectivity alterations associated with lower symptom severity.

**Figure S12:**
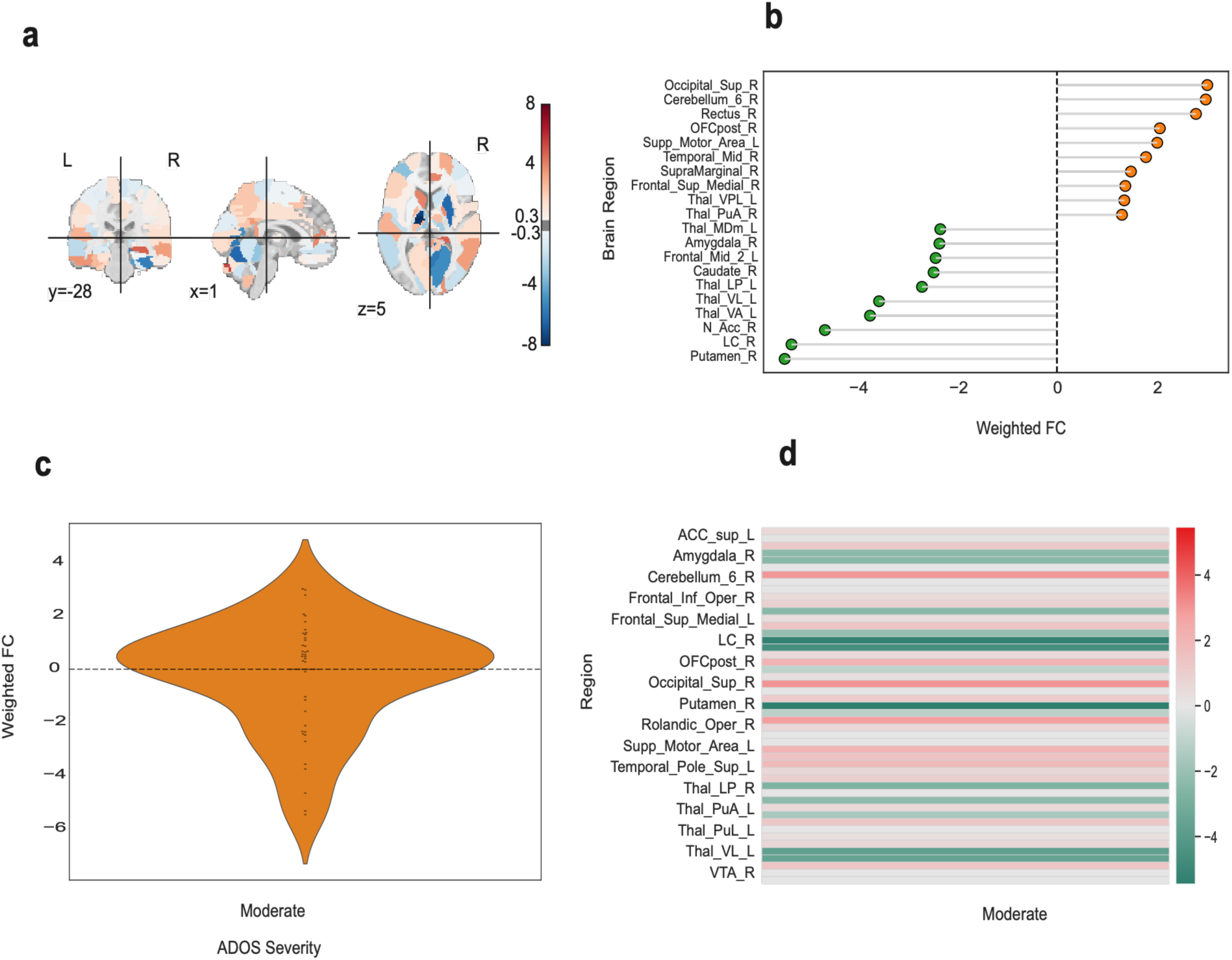
Functional hyper- and hypo-connectivity patterns in intermediate symptom group, demonstrating intermediate regional connectivity alterations.

**Figure S13:**
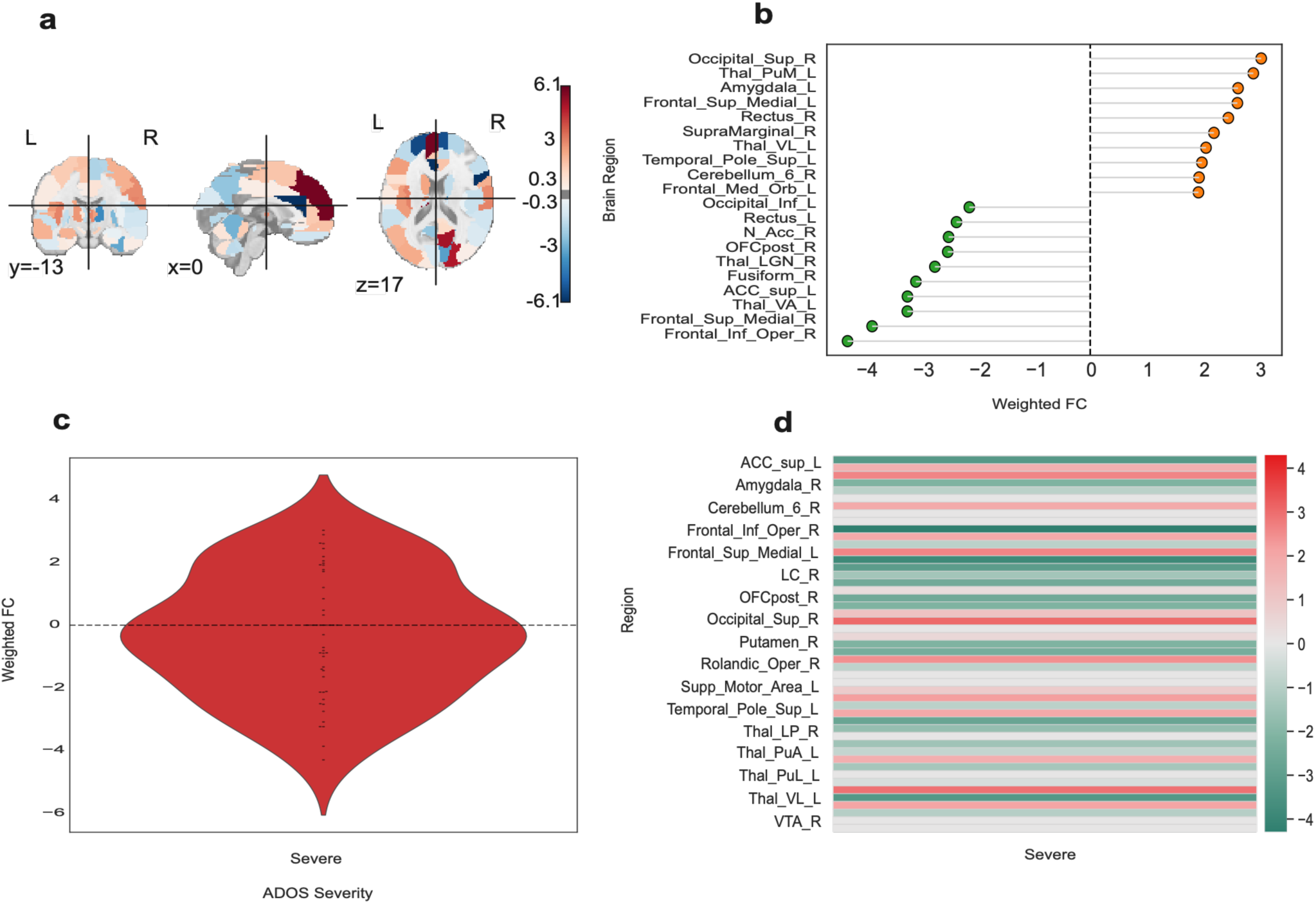
Functional hyper- and hypo-connectivity patterns in high symptom group, showing pronounced alterations across cortical and subcortical functional connectivity networks.

### 1.3 Correlation-Based Hyper- and Hypo-Connectivity Gene Association Analysis

**Figure S14:**
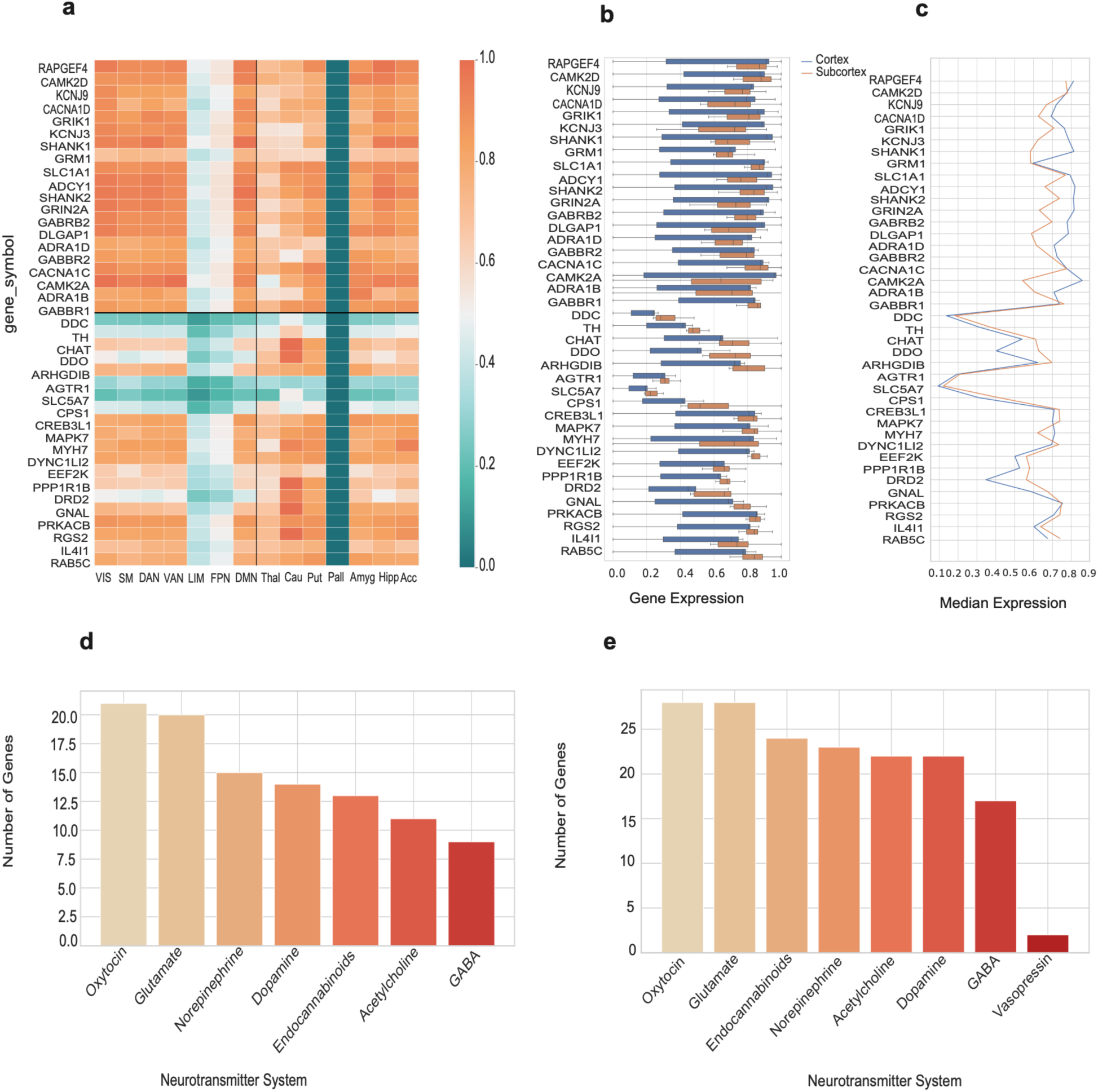
Correlation-based gene expression associations with hyper- and hypo-connectivity patterns in late childhood ASD, showing developmental molecular alterations linked with functional connectivity changes.

**Figure S15:**
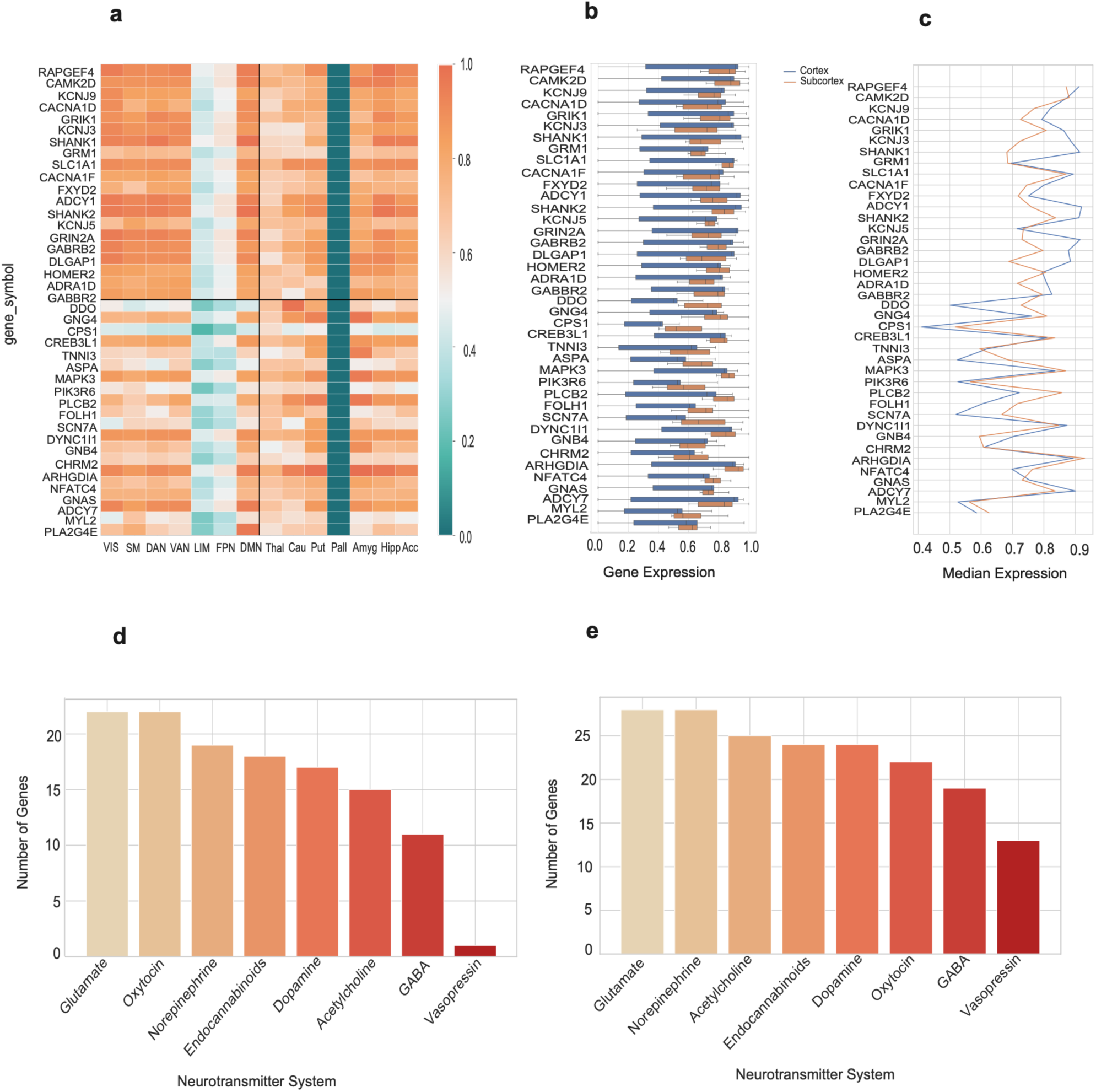
Correlation-based gene expression associations with hyper- and hypo-connectivity patterns in adolescent ASD, highlighting developmental differences in neurotransmitter-related molecular profiles.

**Figure S16:**
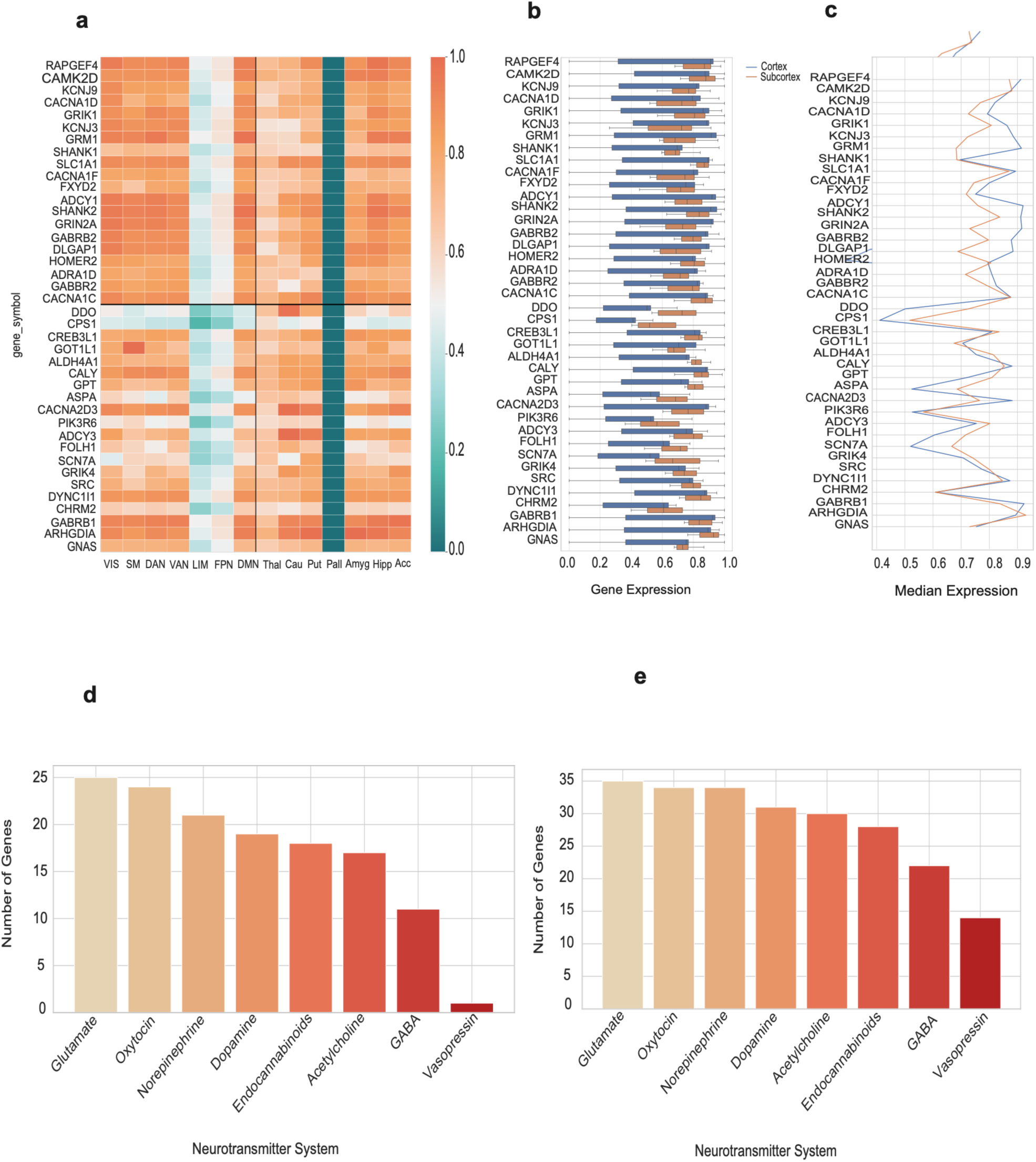
Correlation-based gene expression associations with hyper- and hypo-connectivity patterns in adult ASD, demonstrating persistent and stage-specific molecular connectivity alterations.

**Figure S17:**
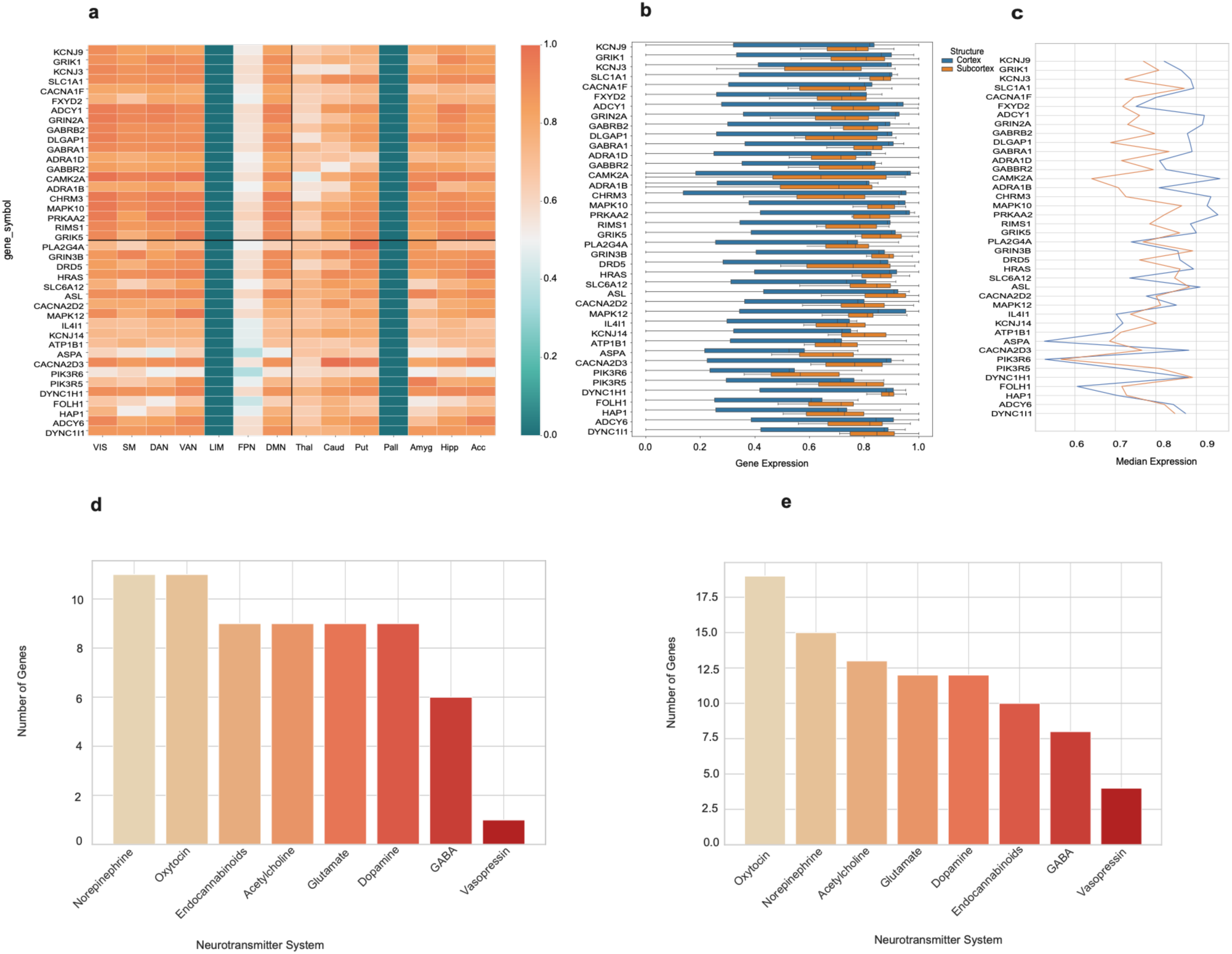
Correlation-based hyper- and hypo-connectivity gene associations in male ASD participants, showing sex-specific neurotransmitter-related molecular patterns.

**Figure S18:**
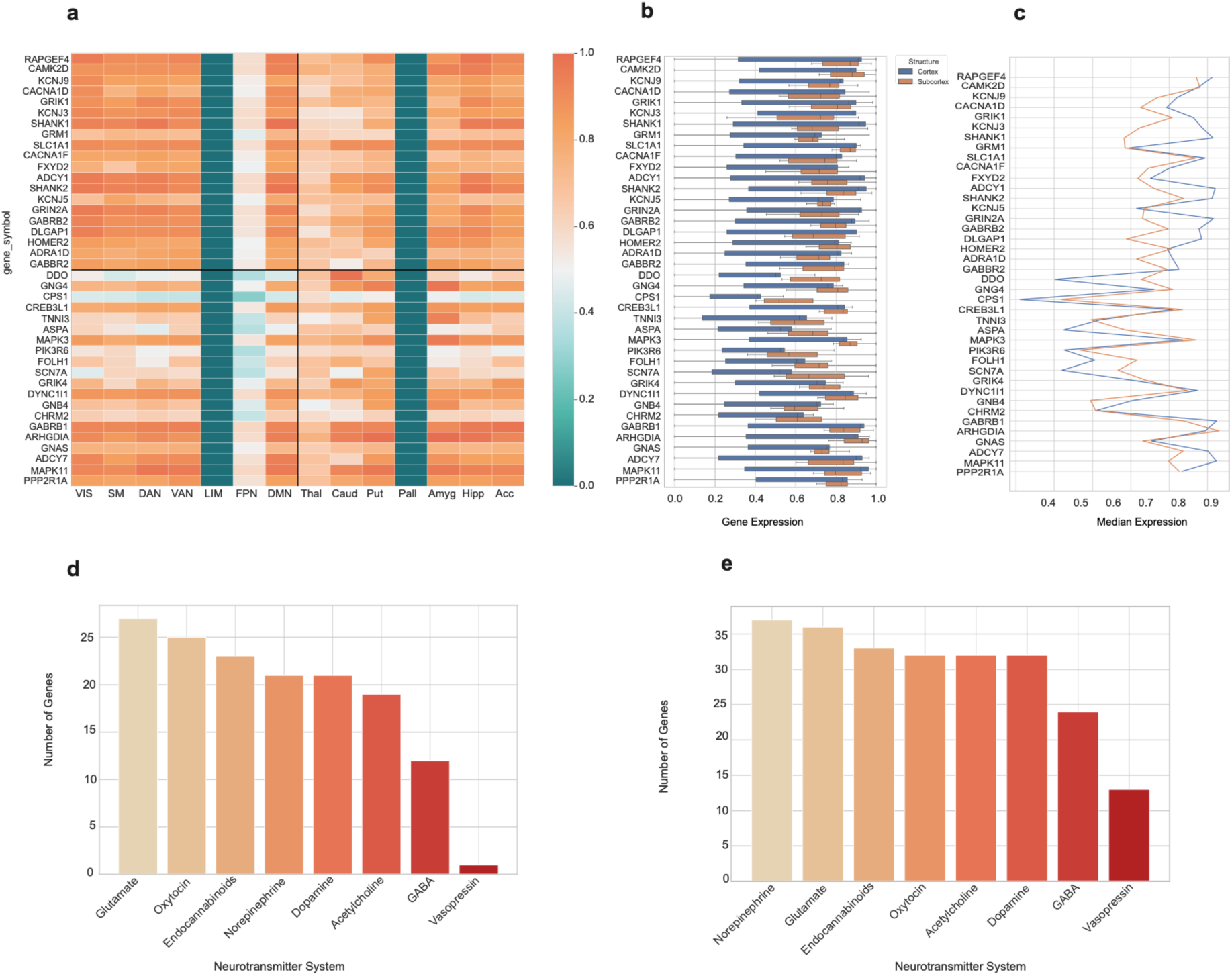
Correlation-based hyper- and hypo-connectivity gene associations in female ASD participants, highlighting sex-related molecular and neurotransmitter-associated differences.

**Figure S19:**
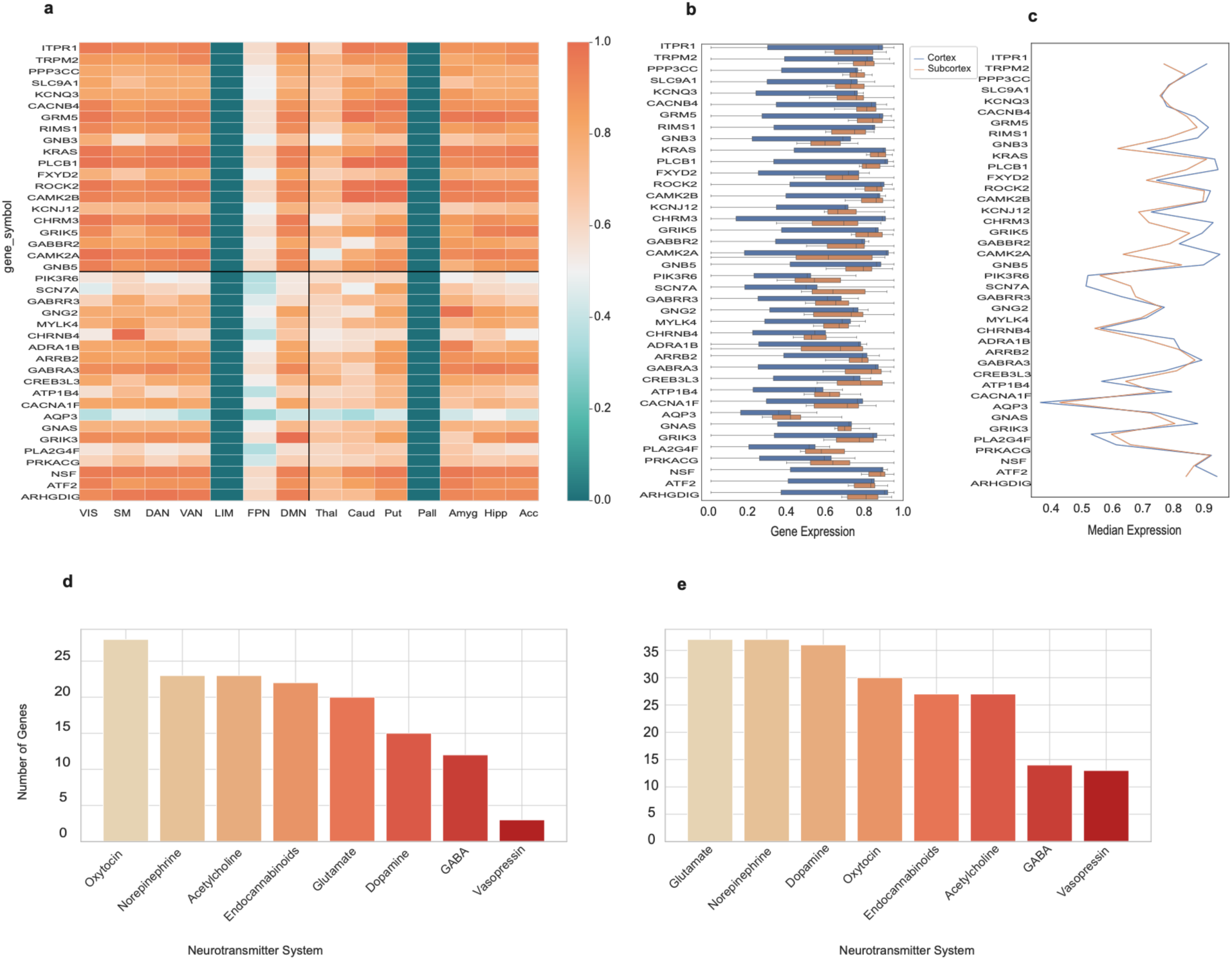
Correlation-based gene expression associations with hyper- and hypo-connectivity patterns in low symptom group, showing molecular alterations linked with lower symptom severity.

**Figure S20:**
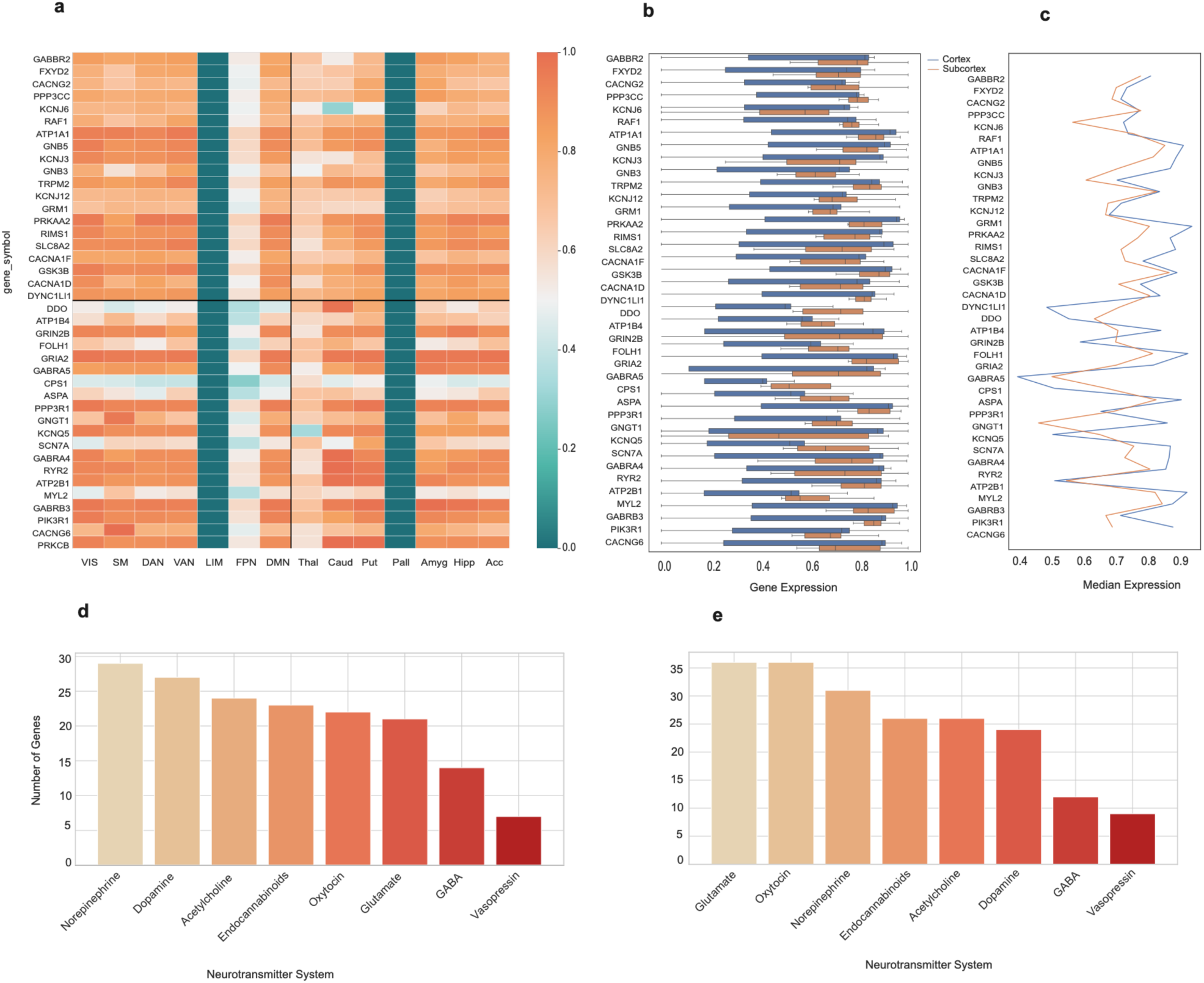
Correlation-based gene expression associations with hyper- and hypo-connectivity patterns in intermediate symptom group, demonstrating intermediate molecular connectivity alterations.

**Figure S21:**
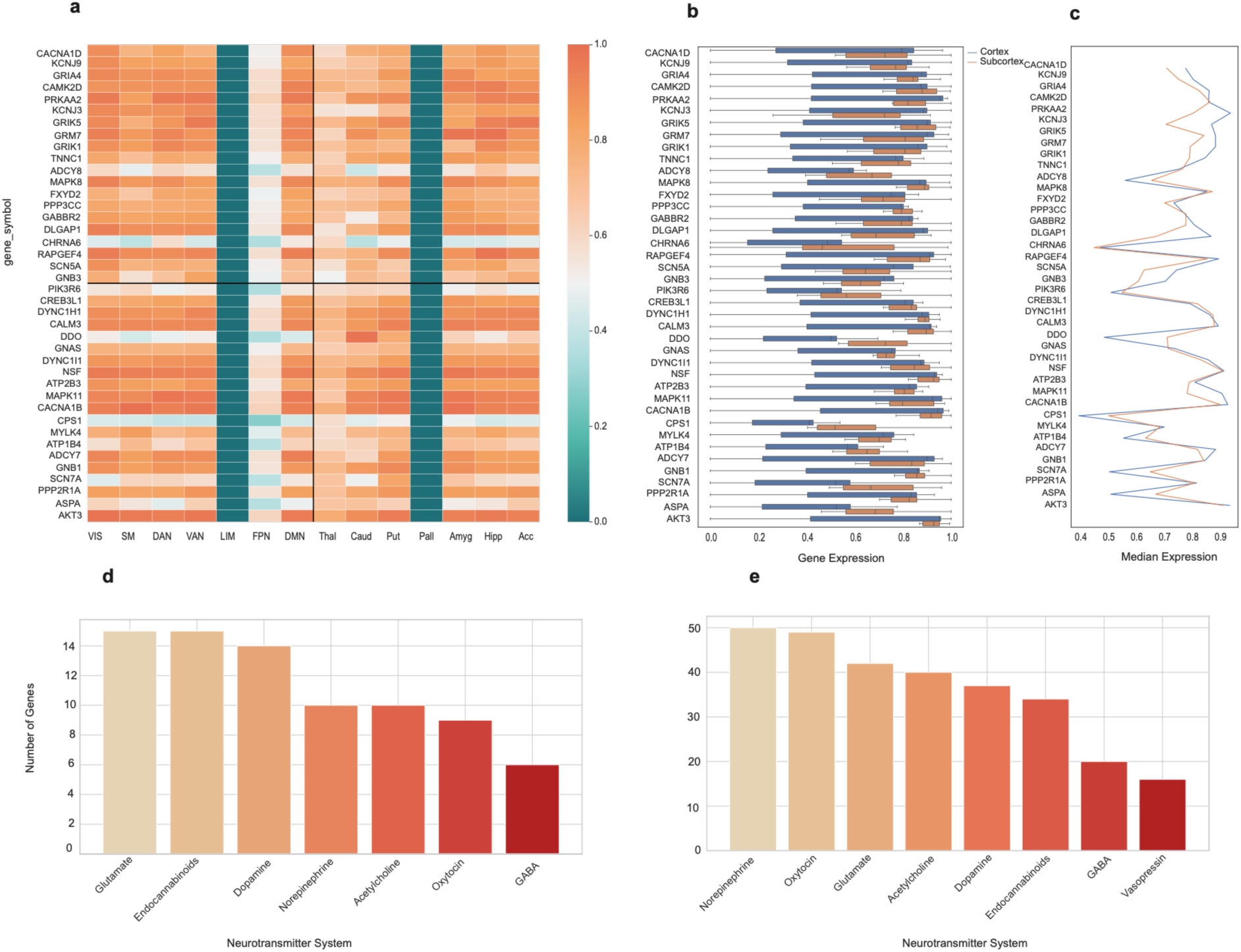
Correlation-based gene expression associations with hyper- and hypo-connectivity patterns in high symptom group, showing pronounced neurotransmitter-related molecular alterations associated with functional connectivity changes.

### 1.4 Functional Gradient Analysis

**Figure S22:**
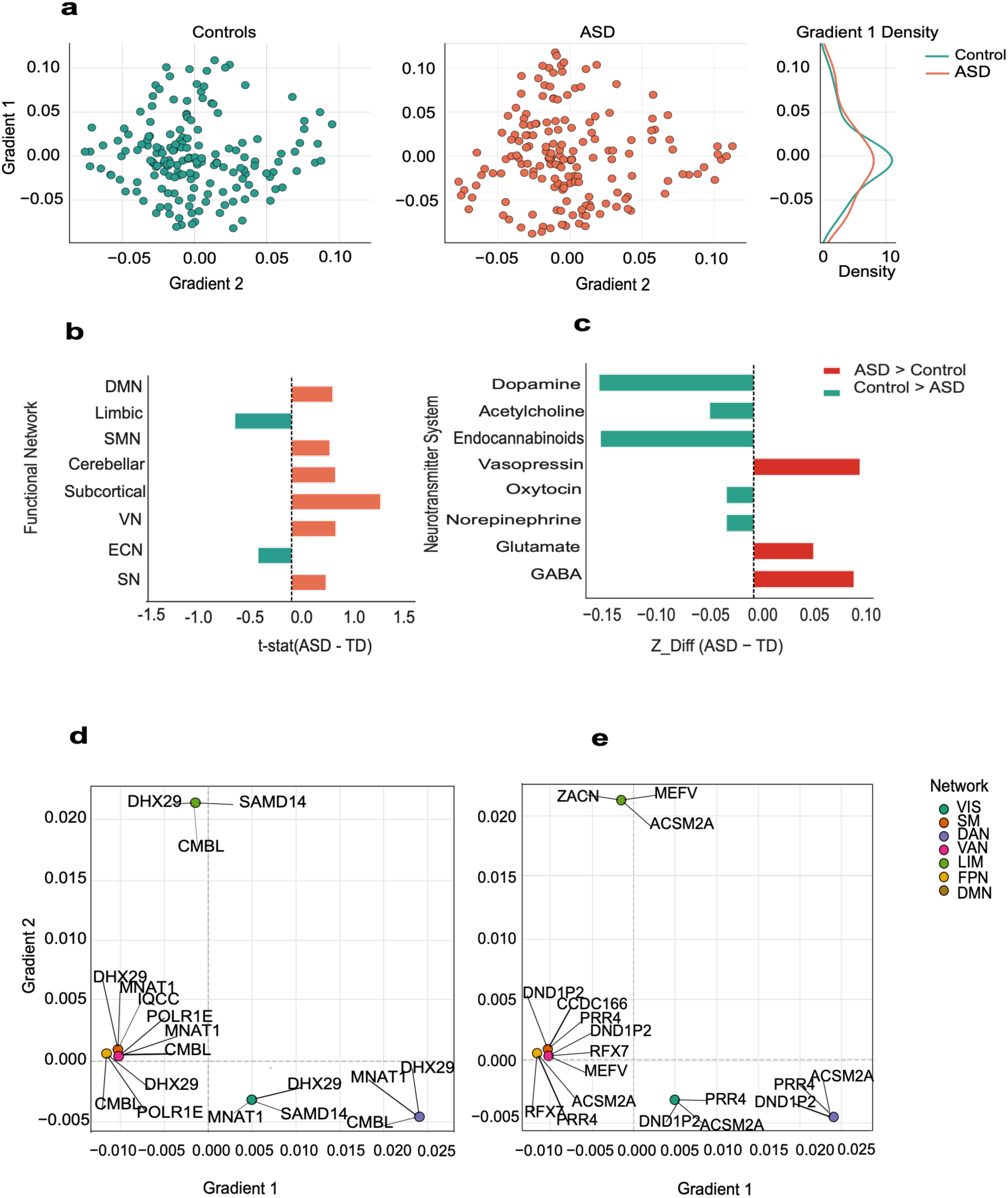
Functional gradient differences in late childhood ASD, highlighting developmental alterations in cortical organization, functional networks, and neurotransmitter-associated spatial patterns.

**Figure S23:**
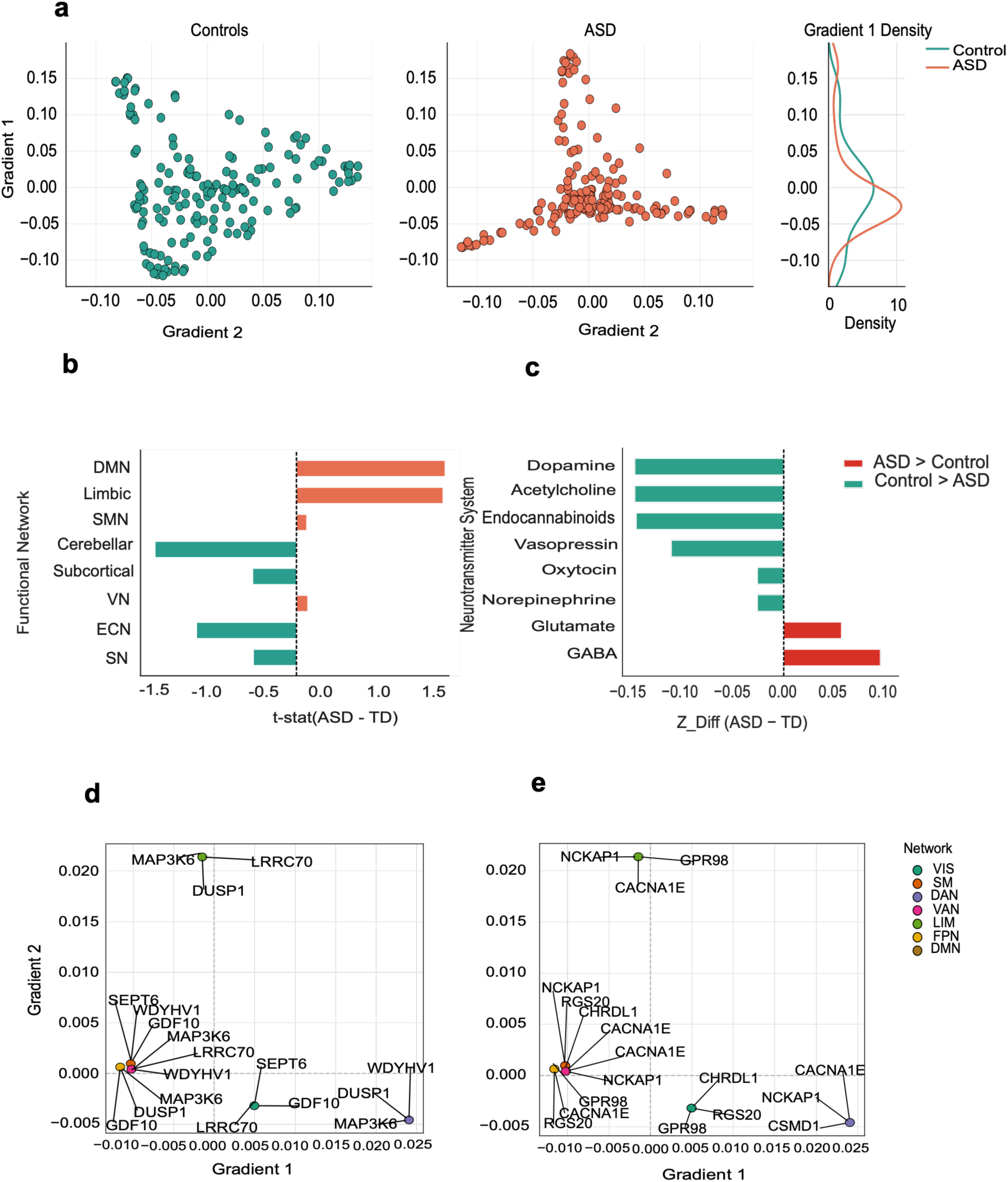
Functional gradient differences in adolescent ASD, showing age-related alterations in cortical hierarchy, network organization, and neurotransmitter-associated gradient patterns.

**Figure S24:**
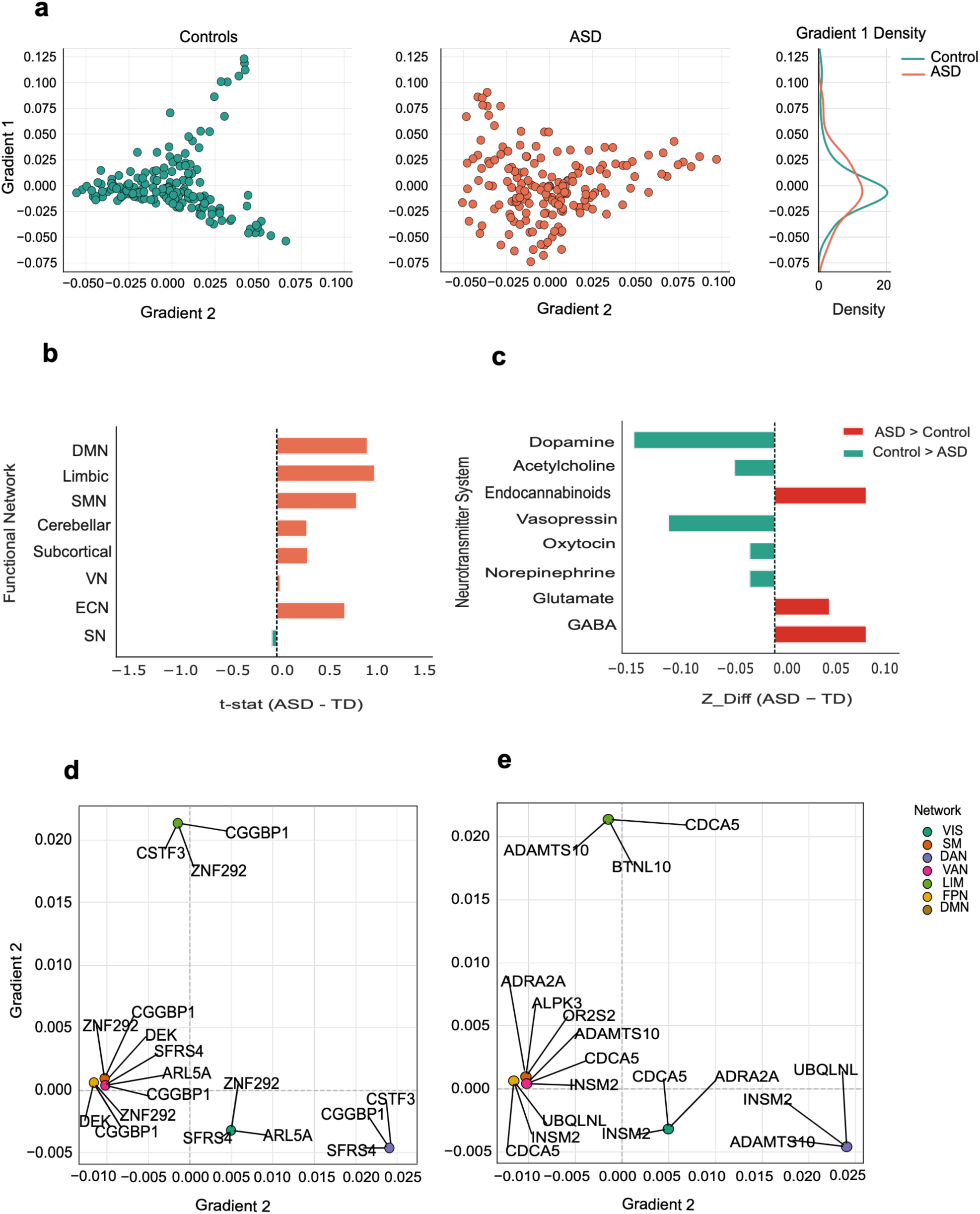
Functional gradient differences in adult ASD, demonstrating persistent and stage-specific alterations in cortical functional organization and neurotransmitter-associated spatial distributions.

**Figure S25:**
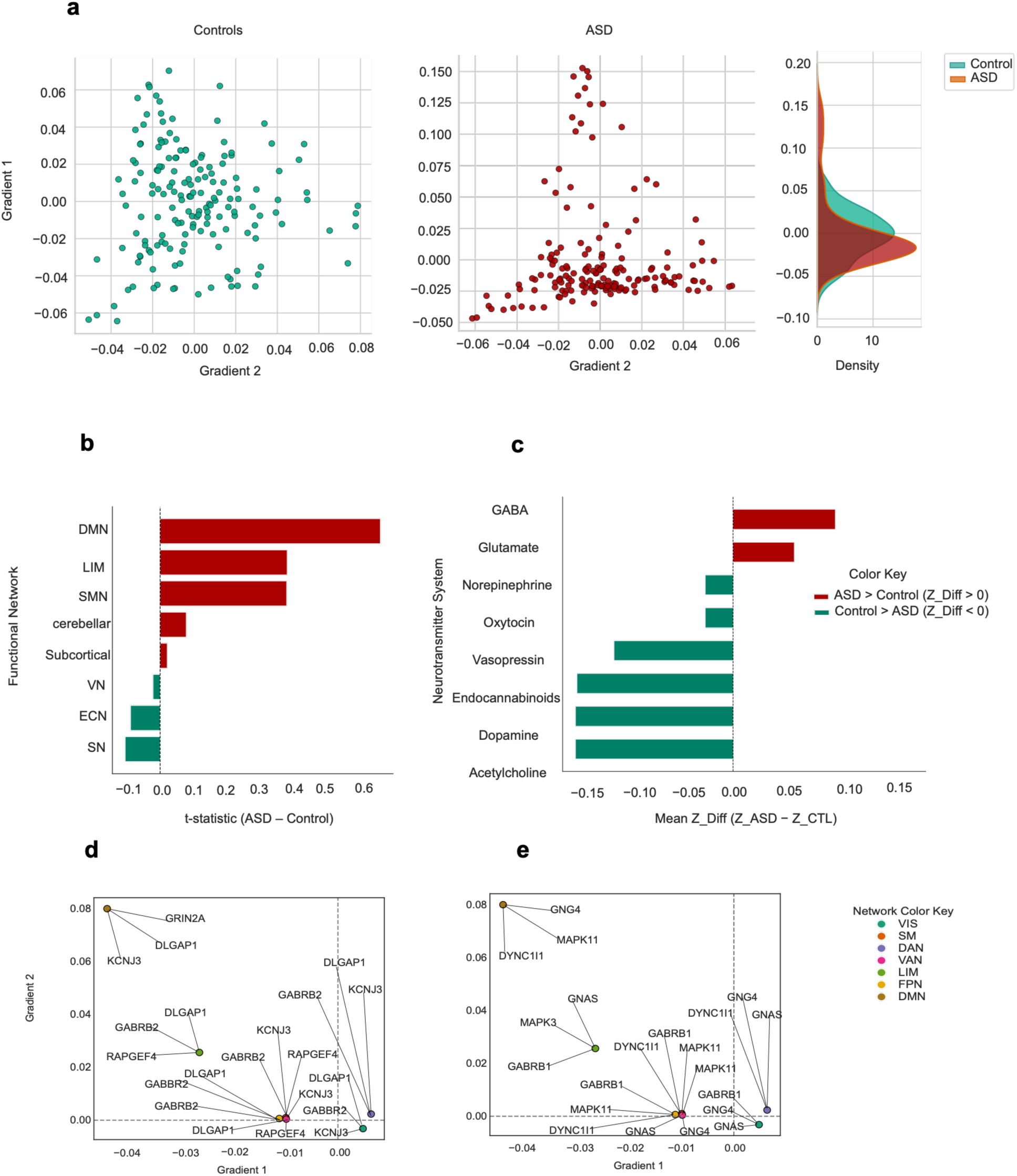
Functional gradient alterations in male ASD participants, showing sex-specific differences in cortical organization, network topology, and neurotransmitter-associated spatial patterns.

**Figure S26:**
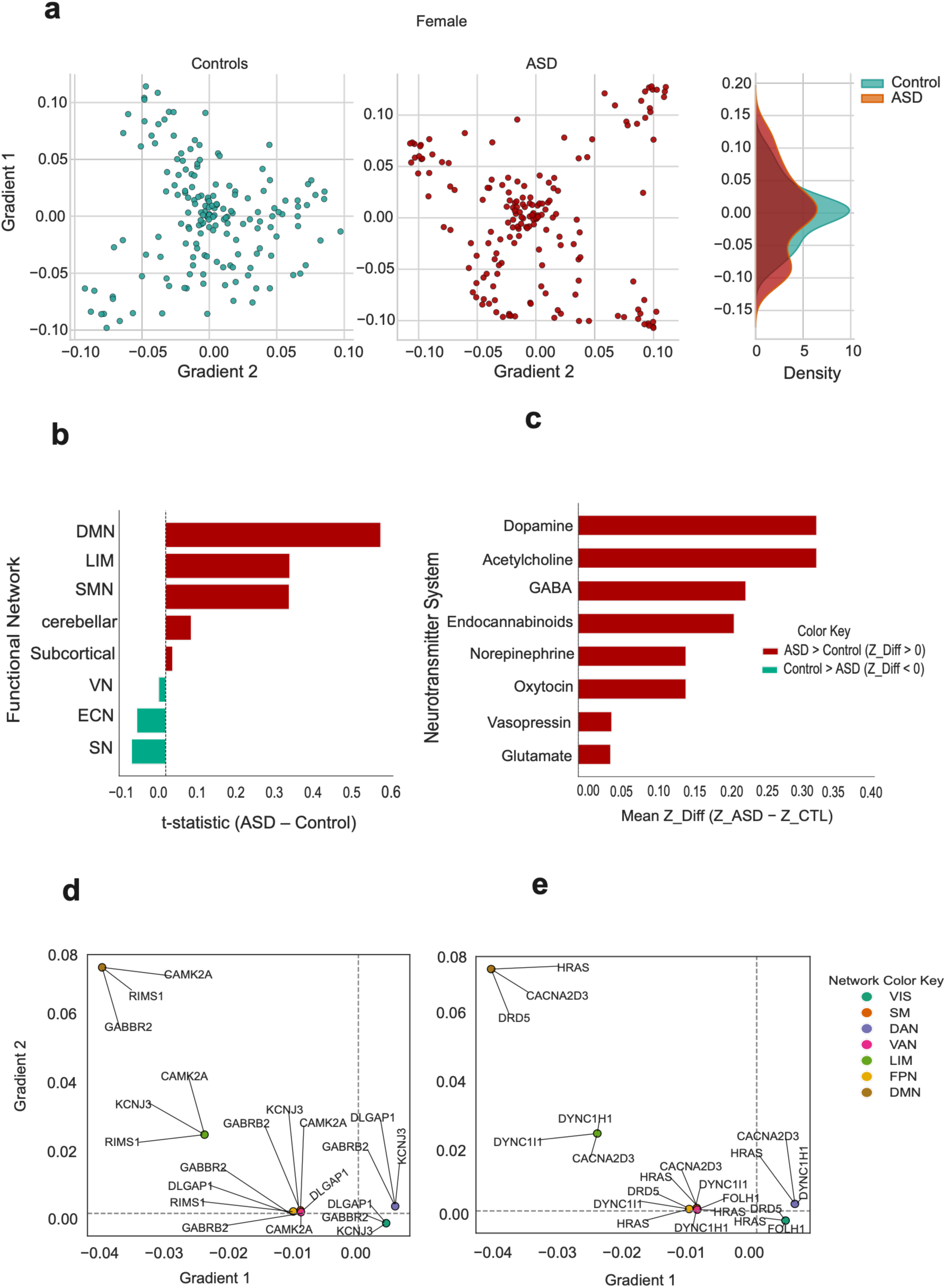
Functional gradient alterations in female ASD participants, highlighting sex-related differences in cortical hierarchy and neurotransmitter-associated functional organization.

**Figure S27:**
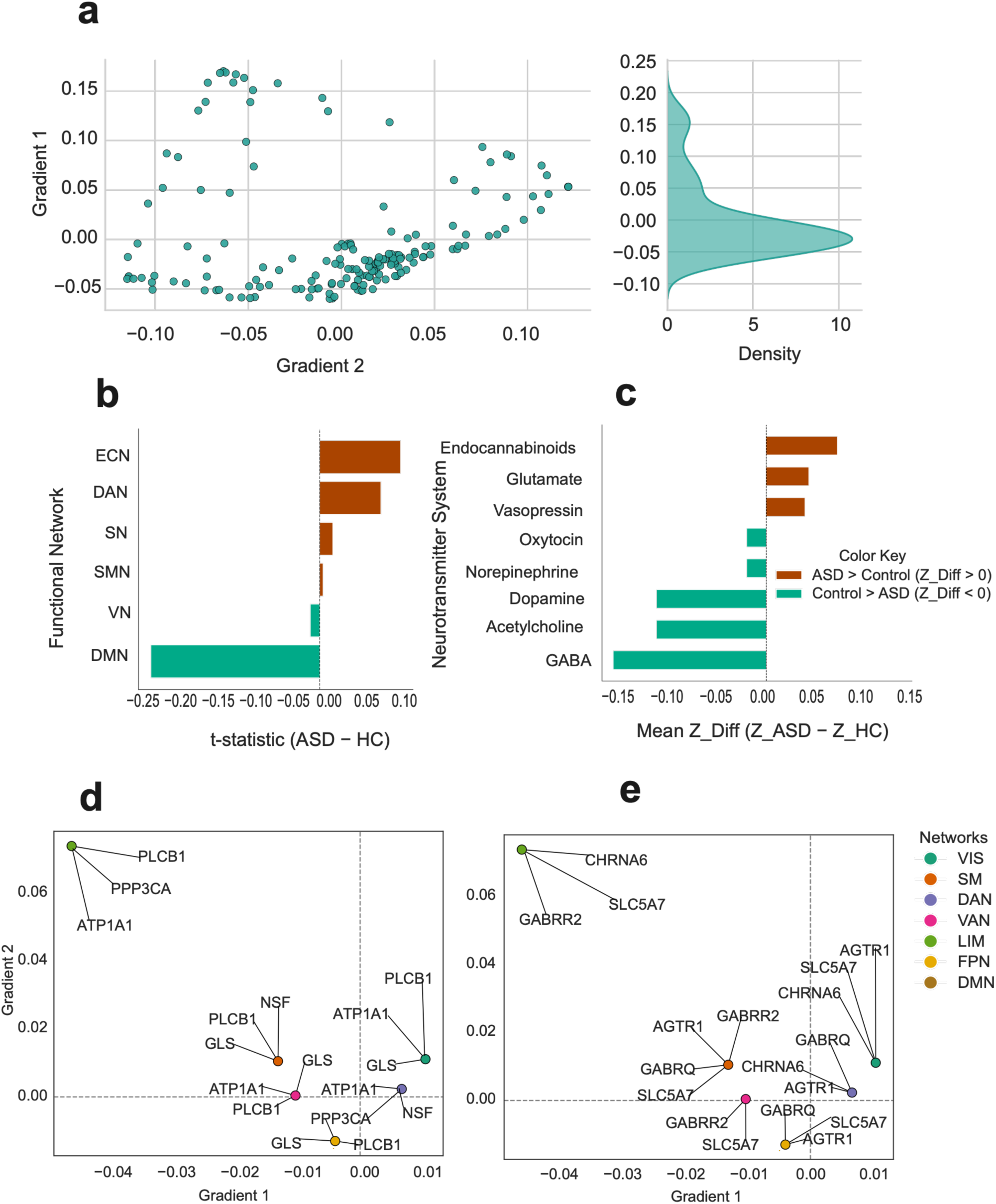
Functional gradient alterations in low symptom group, showing cortical organisational differences associated with lower symptom severity.

**Figure S28:**
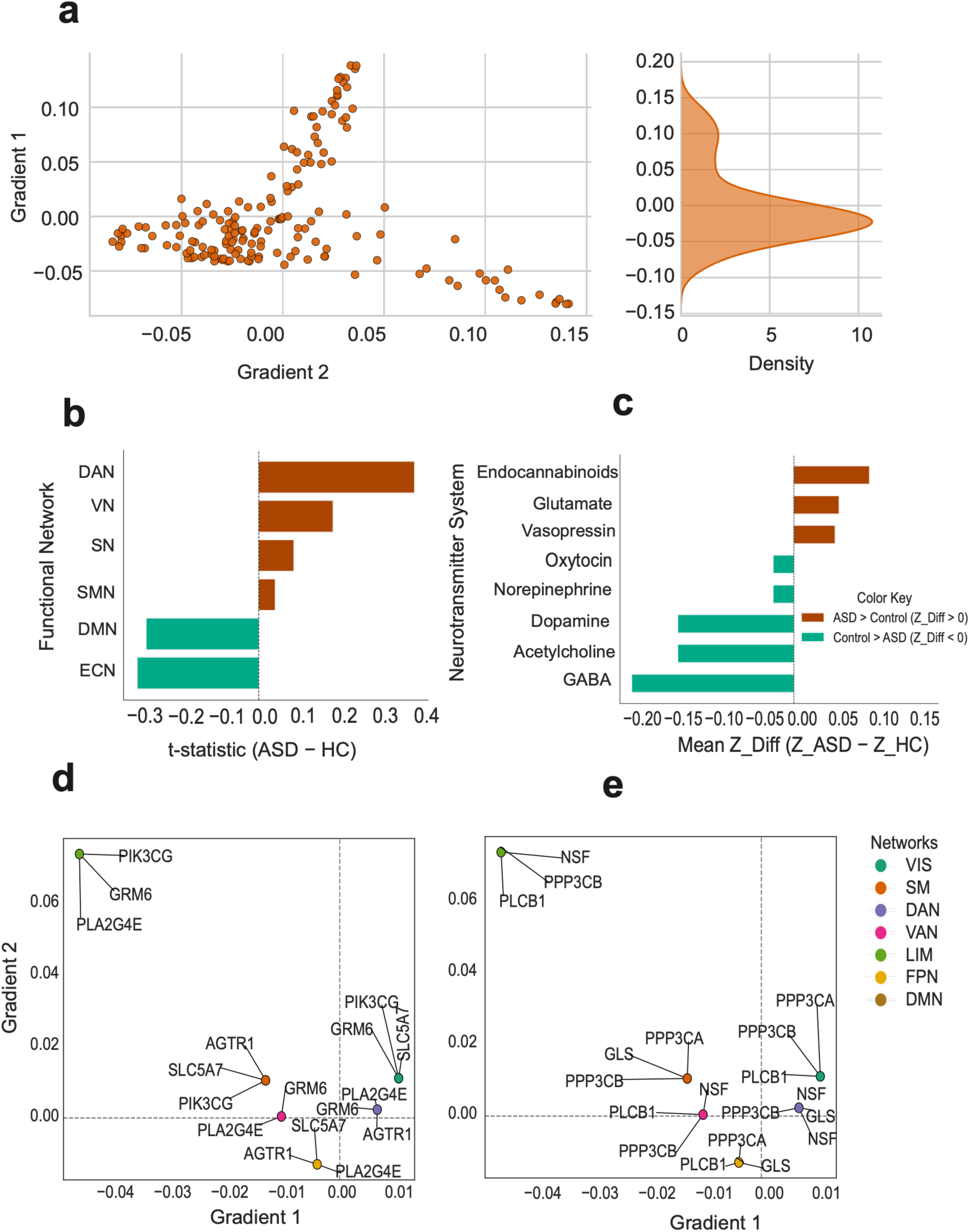
Functional gradient alterations in intermediate symptom group, demonstrating intermediate changes in cortical hierarchy and network-level organization.

**Figure S29:**
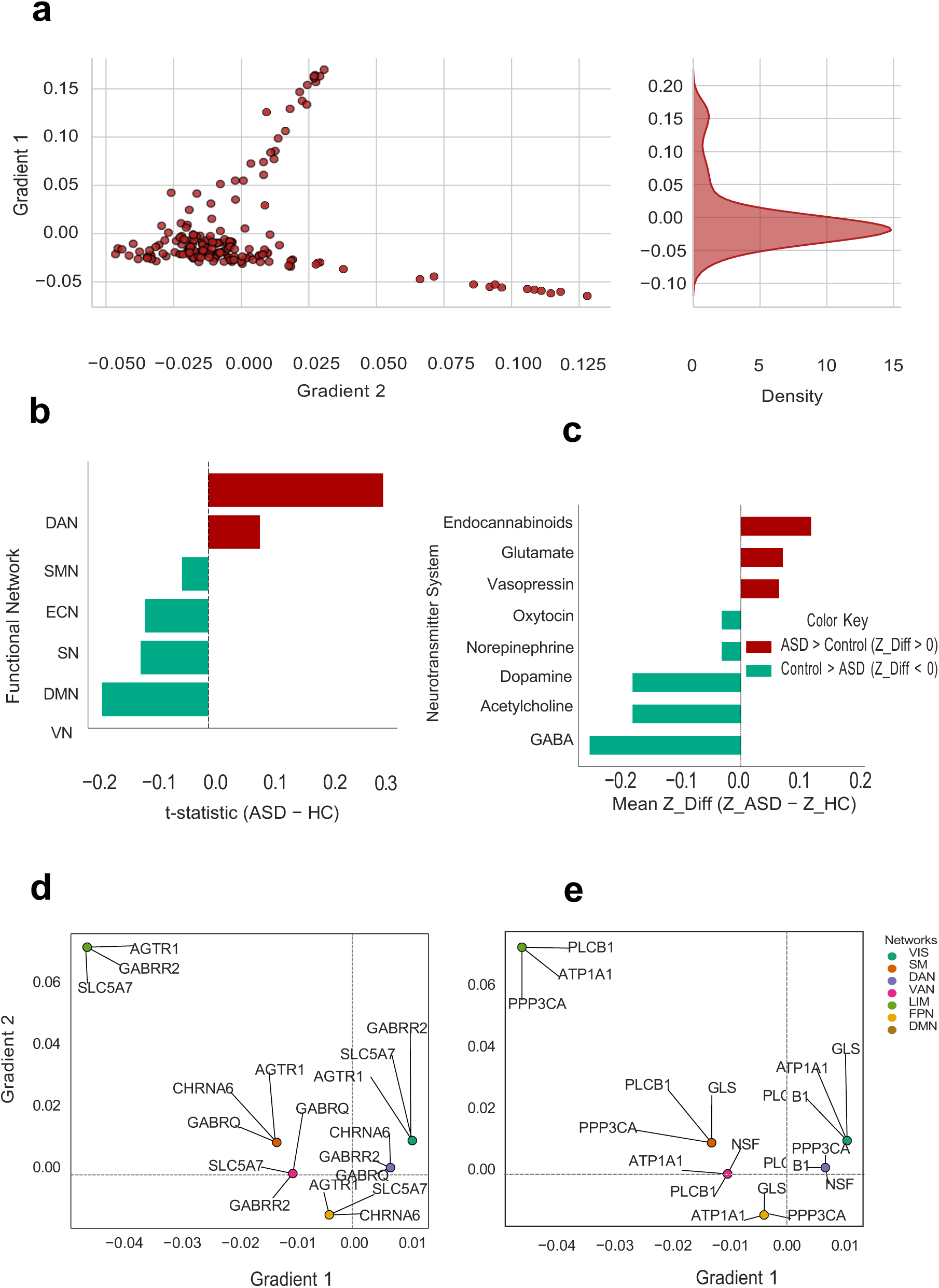
Functional gradient alterations in high symptom group, showing pronounced disruptions in cortical functional organization, network distributions, and neurotransmitter-associated spatial patterns.

**Figure S30:**
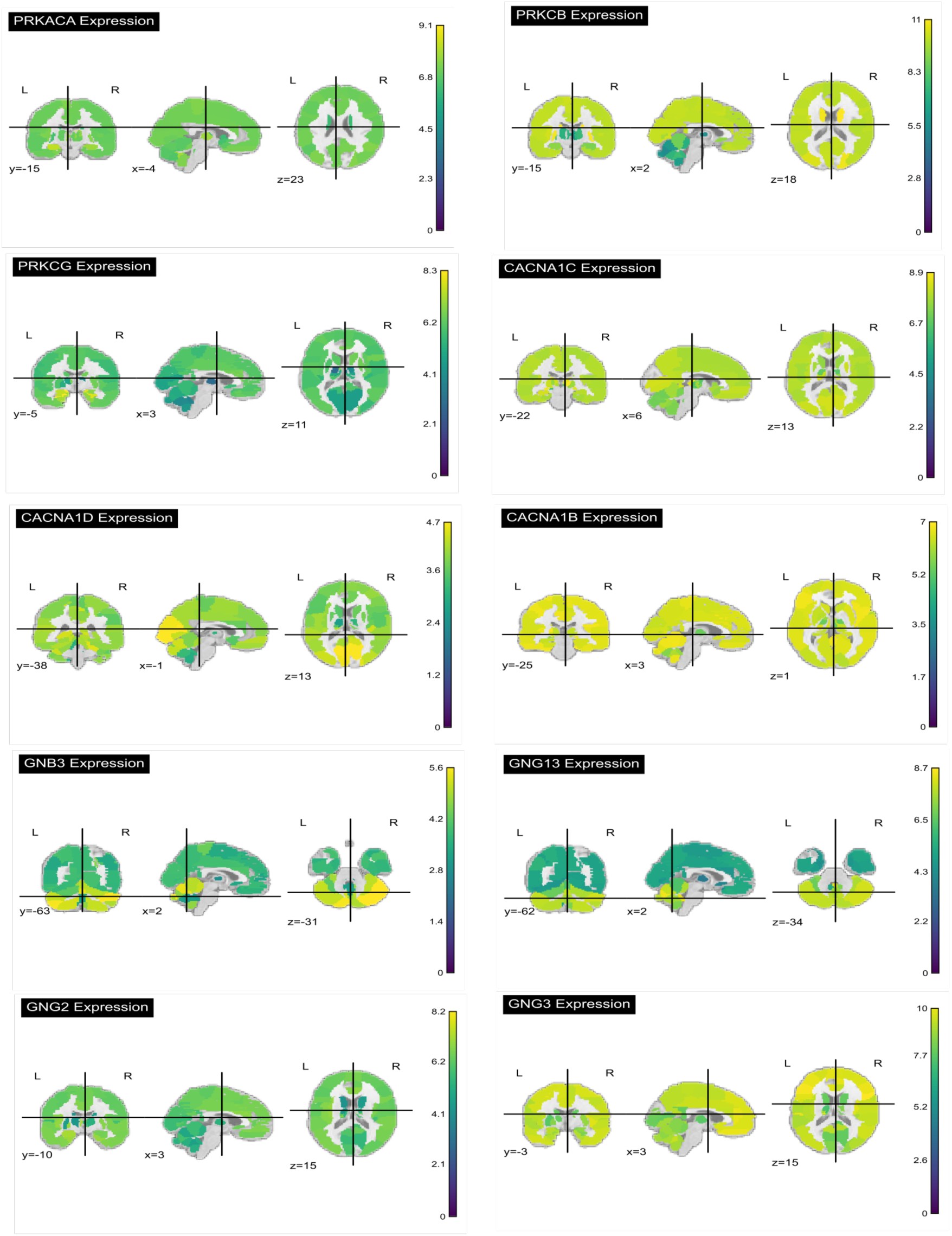
Spatial expression maps of the top ten genes with the highest pathway associations identified from the hierarchical clustering analysis of ASD-enriched pathways. Gene expression values from the Allen Human Brain Atlas were projected onto the AAL3 brain atlas and visualized in orthogonal (axial, coronal, and sagittal) views. The maps illustrate the anatomical distribution of each gene across cortical and subcortical brain regions.

**Figure S31:**
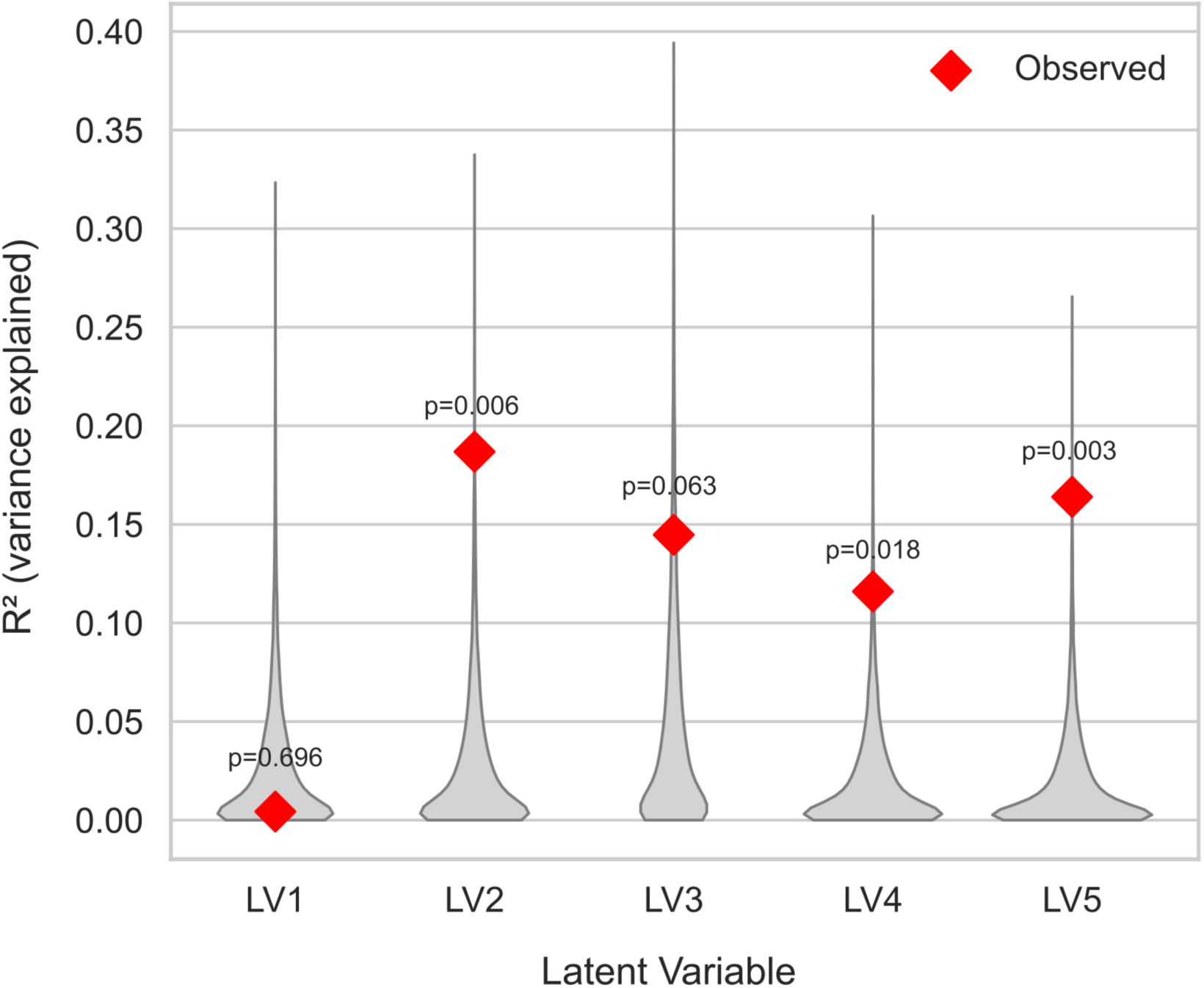
BrainSMASH spatial-null analysis of the PLSRegression model. Null distributions of variance explained (R²) generated from 10,000 BrainSMASH spatially constrained surrogate maps are shown for each latent variable. Red diamonds indicate the observed R² values. The majority of latent variables remained significant following BrainSMASH analysis, supporting the robustness of the multivariate imaging–transcriptomic associations after accounting for spatial autocorrelation.

**Figure S32:**
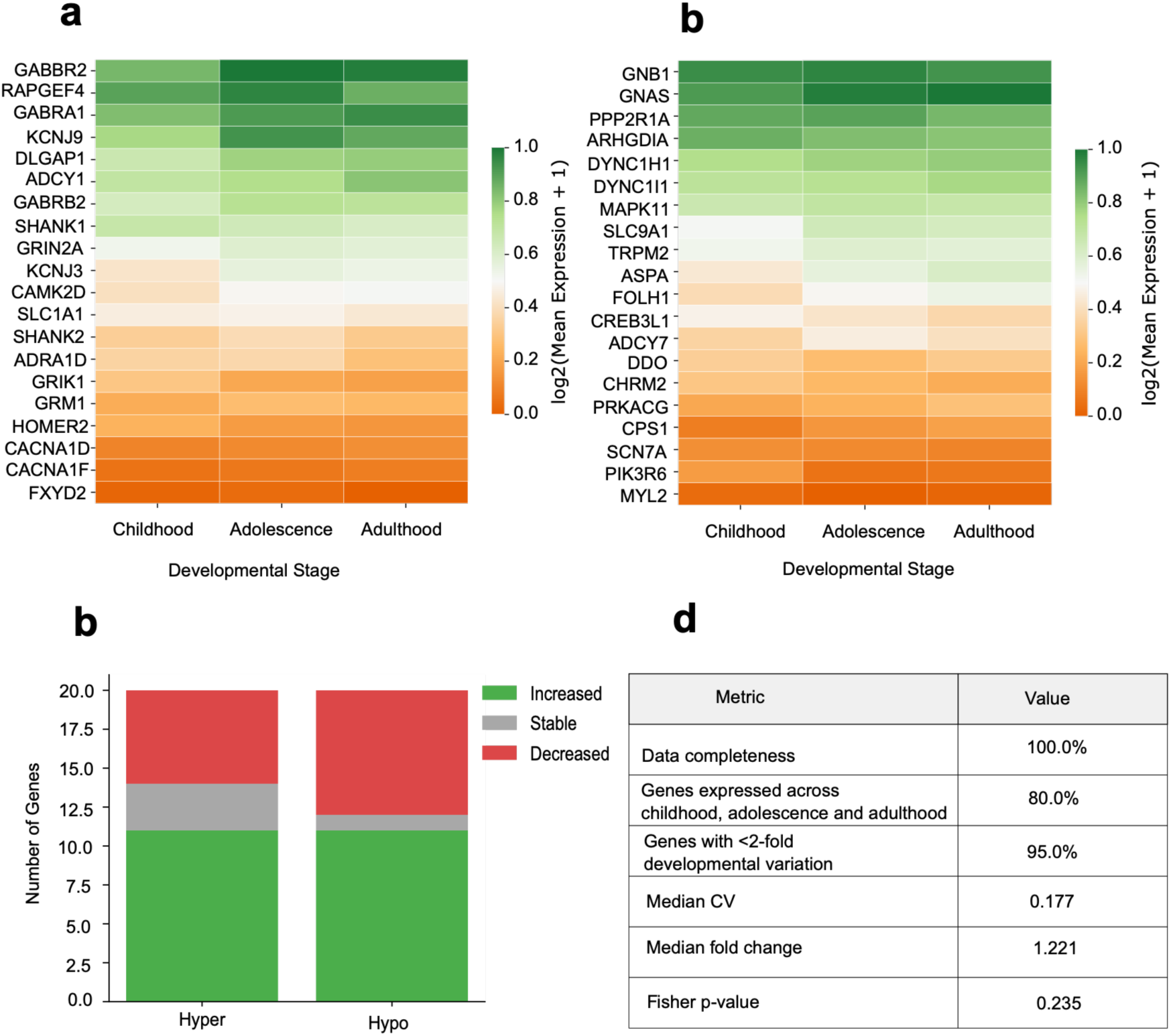
BrainSpan developmental sensitivity analysis of AHBA-derived transcriptomic signatures. (A) Developmental expression heatmap of the top 20 hyperconnectivity-associated genes identified using the Allen Human Brain Atlas (AHBA). (B) Developmental expression heatmap of the top 20 hypoconnectivity-associated genes. Expression values are shown as log_2_(mean expression + 1) across childhood, adolescence, and adulthood using the BrainSpan developmental transcriptomic atlas. (C) Summary of developmental expression trajectories for hyperconnectivity- and hypoconnectivity-associated genes. (D) Summary metrics of the developmental sensitivity analysis, including data completeness, developmental coverage, gene stability, median coefficient of variation (CV), median fold change, and Fisher’s exact test.

**Figure S33:**
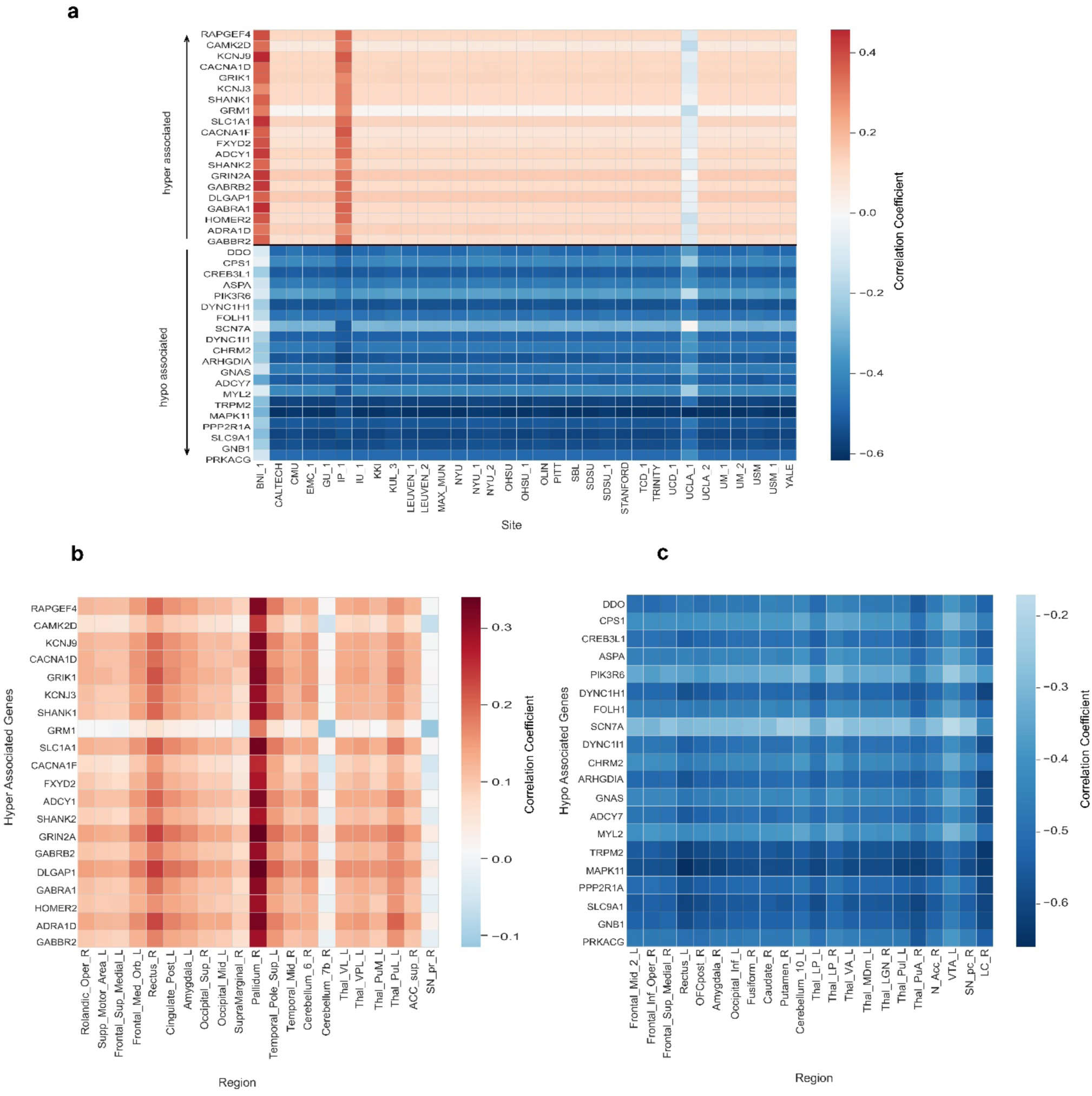
Robustness assessment of imaging–transcriptomic signatures using leave-one-site-out (LOSO) and leave-one-region-out (LORO) analyses. (a) LOSO analysis showing the stability of hyperconnectivity- and hypoconnectivity-associated gene signatures across ABIDE acquisition sites, with each column representing one excluded site. (b) LORO analysis of hyperconnectivity-associated genes following sequential exclusion of each significant brain region. (c) LORO analysis of hypoconnectivity-associated genes following sequential exclusion of each significant brain region. Colour scales represent correlation coefficients between regional functional connectivity alterations and AHBA gene-expression profiles. The consistent direction and magnitude of correlations across iterations demonstrate that the identified transcriptomic signatures are robust to both site-specific variability and regional sampling.

**Table 1:** Participant demographic and clinical characteristics.

| Characteristic | Category | N |
| --- | --- | --- |
| **Total participants** |  | **1,737** |
| **Diagnosis** | Autism spectrum disorder (ASD) | 885 |
|  | Typically developing controls (CTL) | 852 |
| **Sex** | Male | 1,439 |
|  | Female | 298 |
| **Age group\*** | Late childhood (6–12 years) | 575 |
|  | Adolescence (13–17 years) | 612 |
| | Adult ($\geq 18$ years) | 507 |
| **ADOS severity†** | High (ADOS $\geq 12$) | 264 |
|  | Intermediate (ADOS 8–11) | 197 |
| | Low (ADOS $< 8$) | 152 |
|  | Total with ADOS scores | 613 |

**Note:** \*Age groups were defined as late childhood (6–12 years), adolescence (13–17 years), and adulthood ( $\geq 18$ years). †ADOS = Autism Diagnostic Observation Schedule.
Symptom severity categories were defined using total ADOS scores: low ( $< 8$ ), intermediate (8–11), and high ( $\geq 12$ ). Valid ADOS total scores were available for 613 of the 885 participants with ASD.

**Table S1:** Top genes associated with neurotransmitter signaling pathways identified from the hierarchical clustering analysis of ASD-related enriched pathways. The table lists the ten genes with the highest number of pathway associations together with their corresponding neurotransmitter pathways and the total number of enriched pathways in which each gene is involved.

| Gene | Neurotransmitter Pathway | Number of Pathways linked in Hierarchical clustering |
| --- | --- | --- |
| PRKACA | Acetylcholine, Dopamine, Endocannabinoids, GABA, Glutamate, Norepinephrine, Oxytocin, Vasopressin | 7 |
| PRKCB | Acetylcholine, Dopamine, Endocannabinoids, GABA, Glutamate, Oxytocin | 7 |
| PRKCG | Acetylcholine, Dopamine, Endocannabinoids, GABA, Glutamate, Oxytocin | 7 |
| CACNA1C | Acetylcholine, Dopamine, Endocannabinoids, GABA, Glutamate, Norepinephrine, Oxytocin | 6 |
| CACNA1D | Acetylcholine, Dopamine, Endocannabinoids, GABA, Glutamate, Norepinephrine, Oxytocin | 6 |
| CACNA1B | Acetylcholine, Dopamine, Endocannabinoids, GABA | 5 |
| GNB3 | Acetylcholine, Dopamine, Endocannabinoids, GABA, Glutamate | 5 |
| GNG13 | Acetylcholine, Dopamine, Endocannabinoids, GABA, Glutamate | 5 |
| GNG2 | Acetylcholine, Dopamine, Endocannabinoids, GABA, Glutamate | 5 |
| GNG3 | Acetylcholine, Dopamine, Endocannabinoids, GABA, Glutamate | 5 |

**Table S2:**
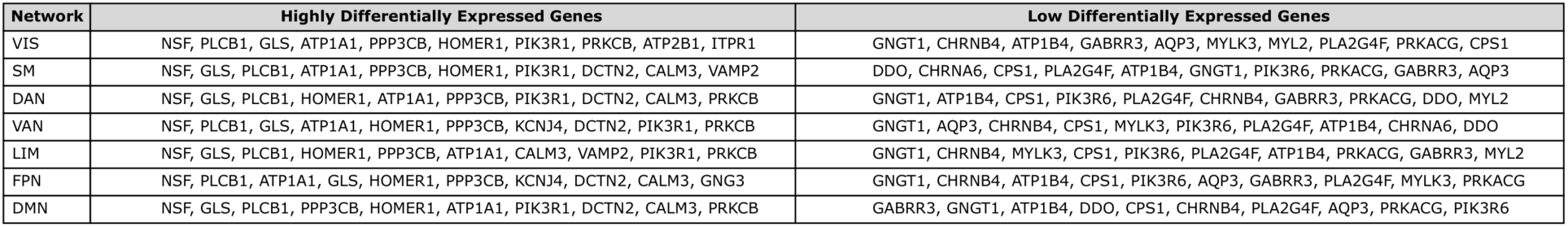
Network-wise gene expression profiles of FGSEA-selected genes across the seven canonical Yeo functional brain networks. For each network, the table lists the ten highest- and ten lowest-expressed genes based on their mean regional gene expression values.

**Table S3:**
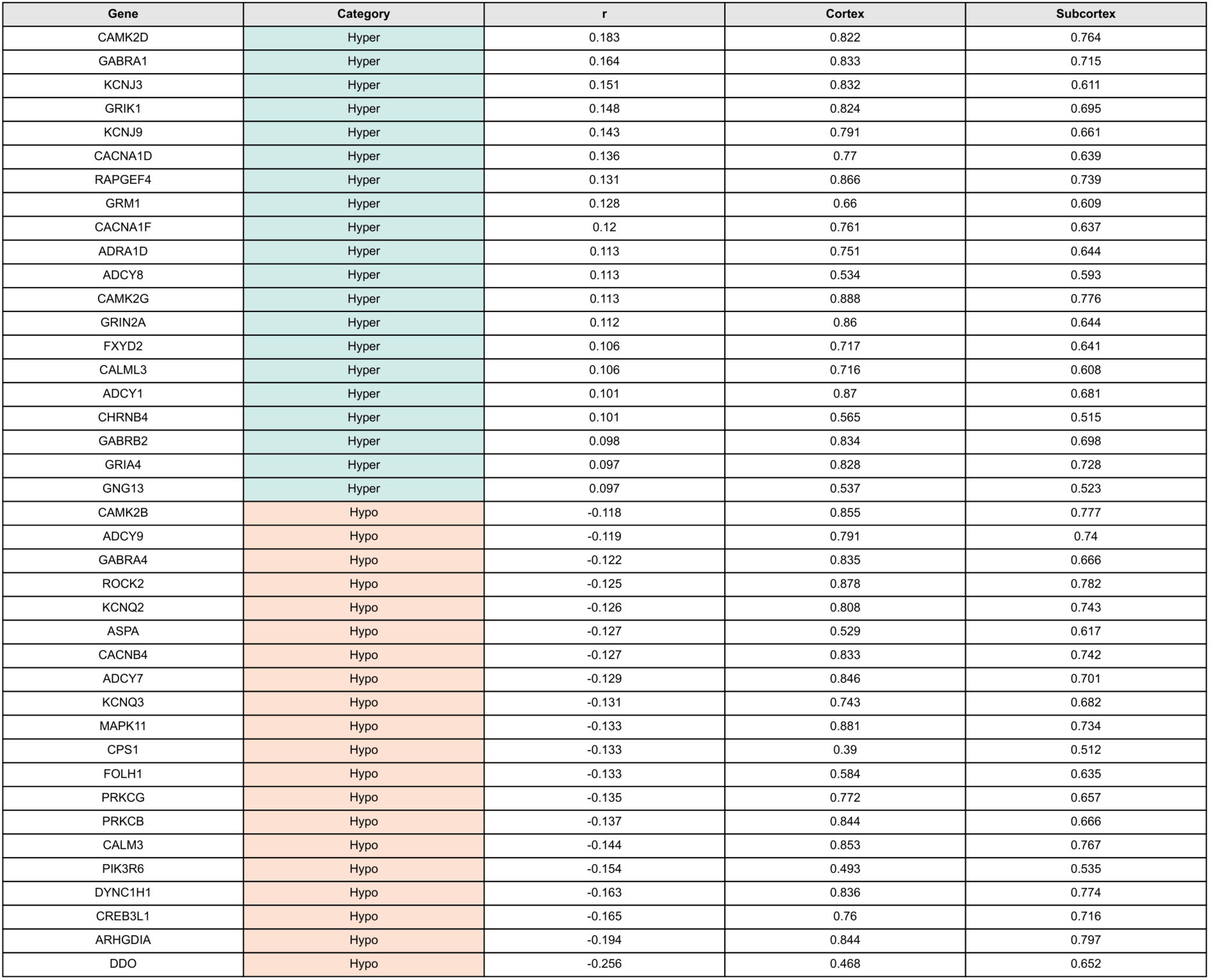
Top genes associated with ASD-related functional connectivity alterations identified through spatial correlation analysis. The table lists the top positively correlated (hyper-connectivity) and negatively correlated (hypo-connectivity) genes together with their Spearman correlation coefficients (*r*) and normalized cortical and subcortical gene expression levels.

**Table S4:** Developmental expression and stability of the 40 study genes in the BrainSpan atlas. The table summarizes developmental expression profiles of the top hyperconnectivity- and hypoconnectivity-associated genes across childhood, adolescence, and adulthood using the BrainSpan developmental transcriptomic atlas. For each gene, the coefficient of variation (CV), adult-to-child fold change, and expression stability status (defined as less than two-fold change across developmental stages) are reported.

| Gene | Connectivity Group | Childhood | Adolescence | Adulthood | CV | Adult/Child Fold Change | Stable (<2-fold) |
| --- | --- | --- | --- | --- | --- | --- | --- |
| ADCY1 | Hyperconnectivity | 26.59 | 30.55 | 33.93 | 0.12 | 1.28 | ✓ |
| ADCY7 | Hypoconnectivity | 1.87 | 3.57 | 2.93 | 0.31 | 1.57 | ✓ |
| ADRA1D | Hyperconnectivity | 6.28 | 6.52 | 4.05 | 0.24 | 0.64 | ✓ |
| ARHGDIA | Hypoconnectivity | 93.4 | 84.2 | 77.88 | 0.09 | 0.83 | ✓ |
| ASPA | Hypoconnectivity | 3.4 | 5.77 | 7.55 | 0.37 | 2.22 |  |
| CACNA1D | Hyperconnectivity | 1.1 | 1.74 | 1.91 | 0.27 | 1.73 | ✓ |
| CACNA1F | Hyperconnectivity | 0.35 | 0.36 | 0.36 | 0.02 | 1.03 | ✓ |
| CAMK2D | Hyperconnectivity | 11.87 | 13.91 | 14.52 | 0.1 | 1.22 | ✓ |
| CHRM2 | Hypoconnectivity | 1.16 | 1.01 | 0.82 | 0.17 | 0.71 | ✓ |
| CPS1 | Hypoconnectivity | 0.27 | 0.33 | 0.53 | 0.36 | 1.98 | ✓ |
| CREB3L1 | Hypoconnectivity | 3.61 | 3.32 | 2.18 | 0.25 | 0.6 | ✓ |
| DDO | Hypoconnectivity | 1.49 | 1.03 | 1.23 | 0.18 | 0.82 | ✓ |
| DLGAP1 | Hyperconnectivity | 24.98 | 33.33 | 33.54 | 0.16 | 1.34 | ✓ |
| DYNC1H1 | Hypoconnectivity | 40.9 | 52.77 | 60.33 | 0.19 | 1.48 | ✓ |
| DYNC1I1 | Hypoconnectivity | 30.51 | 36.81 | 42.12 | 0.16 | 1.38 | ✓ |
| FOLH1 | Hypoconnectivity | 2.46 | 3.76 | 5.31 | 0.37 | 2.16 |  |
| FXYD2 | Hyperconnectivity | 0.08 | 0.1 | 0.07 | 0.18 | 0.82 | ✓ |
| GABBR2 | Hyperconnectivity | 44.66 | 69.41 | 63.16 | 0.22 | 1.41 | ✓ |
| GABRA1 | Hyperconnectivity | 43.59 | 50.98 | 55.81 | 0.12 | 1.28 | ✓ |
| GABRB2 | Hyperconnectivity | 20.94 | 27.96 | 27.69 | 0.16 | 1.32 | ✓ |
| GNAS | Hypoconnectivity | 110.53 | 152.21 | 154.24 | 0.18 | 1.4 | ✓ |
| GNB1 | Hypoconnectivity | 143.27 | 144.84 | 138.08 | 0.02 | 0.96 | ✓ |
| GRIK1 | Hyperconnectivity | 5.04 | 3.28 | 3.19 | 0.27 | 0.63 | ✓ |
| GRIN2A | Hyperconnectivity | 14.93 | 18.5 | 18.2 | 0.12 | 1.22 | ✓ |
| GRM1 | Hyperconnectivity | 3.32 | 3.88 | 3.79 | 0.08 | 1.14 | ✓ |
| HOMER2 | Hyperconnectivity | 3.64 | 3.18 | 2.4 | 0.2 | 0.66 | ✓ |
| KCNJ3 | Hyperconnectivity | 12.3 | 16.55 | 16.16 | 0.16 | 1.31 | ✓ |
| KCNJ9 | Hyperconnectivity | 32.14 | 52.44 | 48.23 | 0.24 | 1.5 | ✓ |
| MAPK11 | Hypoconnectivity | 12.78 | 18.5 | 17.09 | 0.18 | 1.34 | ✓ |
| MYL2 | Hypoconnectivity | 0.04 | 0.01 | 0.02 | 0.66 | 0.69 | ✓ |
| PIK3R6 | Hypoconnectivity | 0.37 | 0.15 | 0.24 | 0.43 | 0.66 | ✓ |
| PPP2R1A | Hypoconnectivity | 99.63 | 103.58 | 89.46 | 0.07 | 0.9 | ✓ |
| PRKACG | Hypoconnectivity | 0.63 | 0.95 | 1.04 | 0.25 | 1.66 | ✓ |
| RAPGEF4 | Hyperconnectivity | 48.71 | 56.16 | 48.16 | 0.09 | 0.99 | ✓ |
| SCN7A | Hypoconnectivity | 0.33 | 0.3 | 0.29 | 0.07 | 0.88 | ✓ |
| SHANK1 | Hyperconnectivity | 25.0 | 22.03 | 20.63 | 0.1 | 0.83 | ✓ |
| SHANK2 | Hyperconnectivity | 6.14 | 6.56 | 5.14 | 0.12 | 0.84 | ✓ |
| SLC1A1 | Hyperconnectivity | 12.99 | 13.1 | 12.74 | 0.01 | 0.98 | ✓ |
| SLC9A1 | Hypoconnectivity | 4.69 | 8.47 | 8.24 | 0.3 | 1.76 | ✓ |
| TRPM2 | Hypoconnectivity | 4.73 | 7.45 | 7.16 | 0.23 | 1.51 | ✓ |

